# Somatic chromosomal integration of polydnavirus during parasitism triggered their germline infiltration in multiple lepidopteran families

**DOI:** 10.1101/2022.09.22.509082

**Authors:** Camille Heisserer, Héloïse Muller, Véronique Jouan, Karine Musset, Georges Périquet, Jean-Michel Drezen, Anne-Nathalie Volkoff, Clément Gilbert

**Author notes:** These authors contributed equally to this work (Camille Heisserer, Héloïse Muller).

## Abstract

Increasing numbers of horizontal transfer (HT) of genes and transposable elements are reported in insects. Yet the mechanisms underlying these transfers remain unknown. Here we first quantify and characterize the patterns of chromosomal integration of the polydnavirus (PDV) encoded by the Campopleginae *Hyposoter didymator* parasitoid wasp (HdIV) in somatic cells of parasitized fall armyworm (*Spodoptera frugiperda*). Polydnaviruses are domesticated viruses injected by wasps together with their eggs into their hosts in order to facilitate the development of wasp larvae. We found that six HdIV DNA circles integrate into the genome of host somatic cells. Each host haploid genome suffers between 23 and 40 integration events (IEs) on average 72 hours post parasitism. Almost all IEs are mediated by DNA double strand breaks occurring in the host integration motif (HIM) of HdIV circles. We show that despite their independent evolutionary origins, PDV from both Campopleginae and Braconidae wasps use remarkably similar mechanisms for chromosomal integration. Next, our similarity search performed on 775 genomes reveals that PDVs of both Campopleginae and Braconidae wasps have recurrently colonized the germline of dozens of lepidopteran species through the same mechanisms they use to integrate into somatic host chromosomes during parasitism. We found evidence of HIM-mediated HT of PDV DNA circles in no less than 124 species belonging to 15 lepidopteran families. Thus, this mechanism underlies a major route of HT of genetic material from wasps to lepidopterans with likely important consequences on lepidopterans.

## Introduction

Parasitoid wasps are a paraphyletic group of Hymenoptera that are classified as ectoparasites or endoparasite insects, depending on whether they develop on or within an arthropod host, respectively (Beckage and Drezen 2012). To ensure the developmental success and survival of their eggs and larvae within hosts, many parasitoid wasps use viral particles that are akin to gene or protein delivery agents (Herniou *et al*. 2013). These particles are injected together with wasp’s eggs in parasitized hosts, of which they alter the physiology, thus enabling the development of wasp larvae within the host body (Beckage and Gelman 2004). The structural components of these viral particles are encoded by genes originating from molecular domestication of viral genomes that were integrated into the genome of wasp’s ancestors (Figure 1).

**Figure 1:**
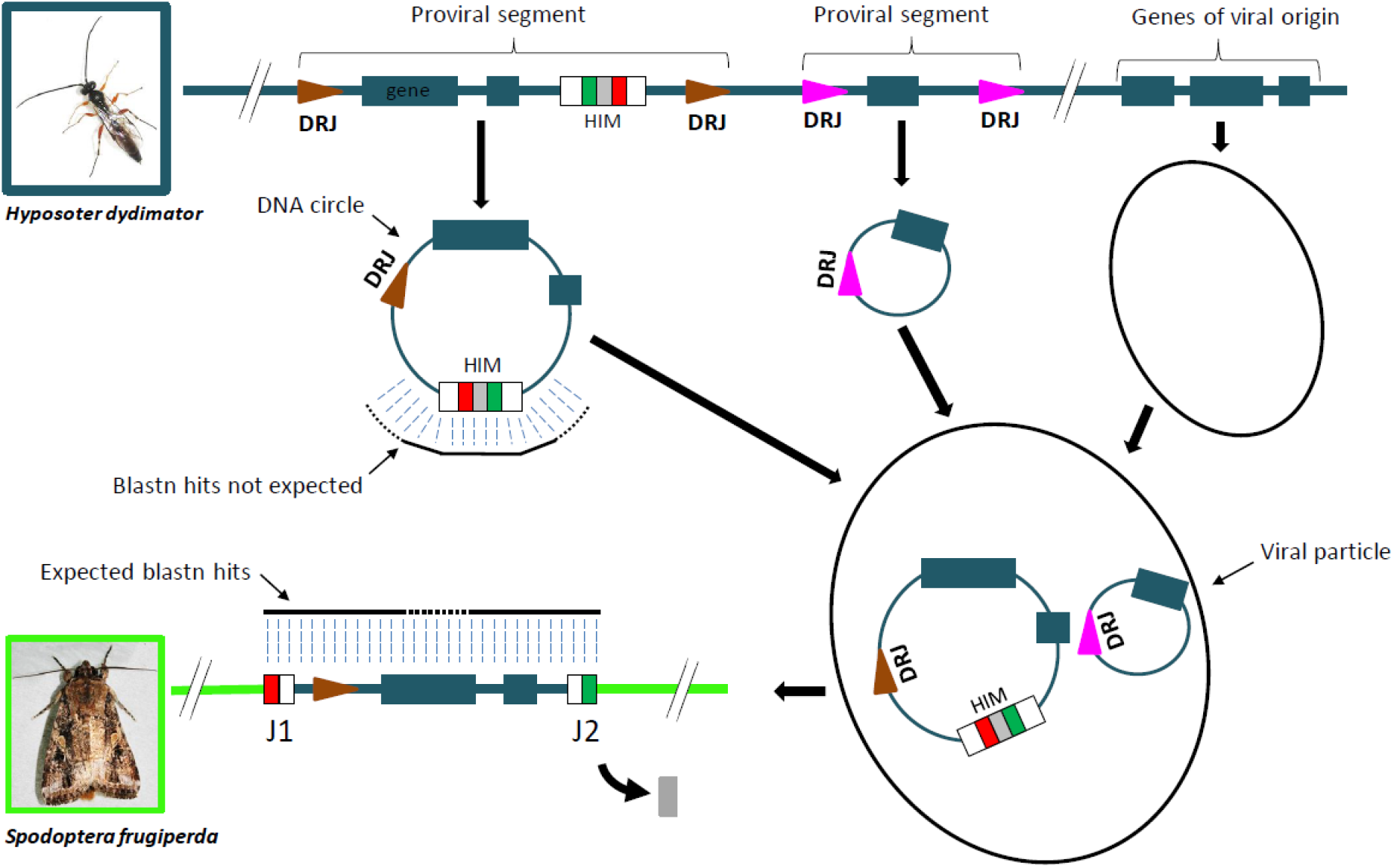
Genome structure and chromosomal integration of polydnaviruses. The genome and genes of the wasp (here *Hyposoter didymator*, Ichneumonidae) are shown in blue while that of the lepidopteran host (here *Spodoptera frugiperda*) is shown in green. The viral particle proteins of polydnaviruses are encoded by genes of viral origin that have been coopted by parasitoid wasps. DNA circles packaged into viral particles originate from proviral segments, which are amplified and circularized in the calyx of female wasps, likely through recombination at the Direct Repeat Junction (DRJ) motifs. Contrary to bracoviruses in which DRJ motifs are similar between proviral segments, DRJ motifs differ between ichnovirus segments, hence they are depicted with different colors (pink and brown). While ovipositing into larvae of their lepidopteran hosts, female wasps inject large amounts of PDV viral particles, which enter host somatic cells and deliver DNA circles to host cell nuclei. DNA circles possessing a Host Integration Motif (HIM) then undergo chromosomal integration catalyzed by wasp and host integrases (Wang *et al*. 2021). Integration involves a double strand break within the HIM and linearized circles newly integrated into host DNA are bordered by junction 1 (J1 in red) and junction 2 (J2 in green) motifs. The short intervening region between J1 and J2 (in grey; about 50 bp in most circles) is lost during the process. We predicted that if horizontal transfer of PDV sequences occur recurrently in lepidopteran hosts through HIM-mediated germline genome integration, few or no PDV sequence integrated in lepidopteran genomes should contain the intervening region. Thus we should recover few or no blastn hit covering this region. By contrast, most or all blastn hits covering the HIM region should end at the J1 or J2 motif, which lie at the predicted junction with host DNA. The picture of *H. didymator* and *S. frugiperda* were taken by Marie Frayssinet, INRAE DGIMI lab.

Such domestication events occurred multiple times independently in different hymenopteran lineages, yielding viral particles that differ in terms of structure, function and content (Volkoff *et al*. 2010; Beliveau *et al*. 2015; Pichon *et al*. 2015; Burke 2019; Di Giovanni *et al*. 2020; Burke *et al*. 2021; Mao *et al*. 2022). Given their viral origin, the particles packaging DNAs encoded by parasitoid wasps have been referred to as viruses and classified in their own viral family, the Polydnaviridae, or PDVs (Stoltz *et al*. 1984; Herniou *et al*. 2013). The two currently recognized genera of PDVs are the bracoviruses (BVs) and the ichnoviruses (IVs), respectively carried by wasps of the Braconidae and Ichneumonidae families that parasitize mainly lepidopteran hosts (Bézier *et al*. 2009; Beliveau *et al*. 2015; Burke *et al*. 2018; Volkoff and Cusson 2020). Bracoviruses result from the domestication of a nudivirus integrated in a braconid ancestor ∼100 million years ago (Ma) (Bézier *et al*. 2009), and are found today in all wasps belonging to the “Microgastroid complex” estimated to contain about 50,000 species (Murphy *et al*. 2008). Ichnoviruses derive from an unknown viral family (Volkoff *et al*. 2010) and are present in two related Ichneumonidae subfamilies containing about 4,000 species. It was questioned whether their distribution in Ichneumonidae is due to one or two independent integration events (Béliveau *et al*. 2015) but recent phylogenetic analyses favor the second hypothesis (Santos *et al*. 2022).

A remarkable feature of BVs and IVs is that they both package circular DNA molecules (Volkoff and Huguet 2021 ; Figure 1). These DNA circles arise from amplification and excision of so-called “proviral segments” located in the wasp genome. The segments contain genes of different origins that encode virulence factors and are expressed in the parasitized host throughout the development of the wasp larvae. In braconid wasp genomes, most BV segments are arranged in tandem in large loci. For example, *Cotesia congregata* harbors 35 proviral segments, 17 of which are clustered in an ∼2-Mb macrolocus located on chromosome 5 (Gauthier *et al*. 2021). Each of the segments is flanked by direct repeat junctions (DRJs), sequence motifs that all contain an AGCT tetramer motif at which circularization through site-specific recombination occurs (Drezen *et al*. 1997; Desjardins *et al*. 2008; Burke *et al*. 2015). Ichneumonid IV segments are generally more numerous than BV segments and are highly dispersed in wasp genomes. For example, *Hyposoter didymator* harbors 57 proviral segments that are all separated by megabase-long portions of wasp genome (Legeai *et al*. 2020). Like BV segments, IV segments are flanked by DRJs but contrary to BVs, which segments share similar extremities, the sequence of IV DRJs are specific of each segment. Circularization is thought to occur through homologous recombination between left and right DRJs, with breakpoints located at varying positions along the DRJs (Legeai *et al*. 2020).

While the different genomic organization of BV and IV proviral segments imply differences in the mechanisms regulating their amplification and circularization, DNA circles are produced in cells of the calyx, a special region of the female gonad, for both PDV types. They are then packaged into viral particles, released in the oviduct lumen and injected into the host during oviposition. Early cell line-based studies suggested that once in host cells, some DNA circles could persist as chromosomally integrated forms (Figure 1) (McKelvey *et al*. 1996; Volkoff *et al*. 2001; Gundersen-Rindal and Lynn 2003). This was confirmed using targeted Sanger and Illumina sequencing by showing that at least a subset of BV circles integrates into host hemocytes, and that integration likely involves an enzymatically regulated process through recognition of a specific motif of the DNA circles called host integration motif (HIM) (Beck *et al*. 2011; Chevignon *et al*. 2018). In agreement with this, a recent study found by functional analysis that in some lepidopteran hosts, a fraction of DNA circle integration events are catalyzed by a series of viral integrases, while host integrases are also involved in the integration of some circles (Zehua Wang *et al*. 2021). Using bulk Illumina sequencing of parasitized Mediterranean corn borer larvae (*Sesamia nonagrioides*, Lepidoptera, Noctuidae) we showed that chromosomal integration of BV circles from *C. typhae* occurs systematically in multiple host tissues and that only BV DNA circles containing HIM motifs integrate at measurable levels. Our quantitative approach also allowed us to estimate that each host haploid genome suffers between 12 and 85 BV DNA circle integration events depending on the tissue. Despite this high level of integration activity, we found that non-integrated circles also persist as much as integrated circles in parasitized hosts during at least half the development of wasp larvae (Muller *et al*. 2021). Recent bulk Illumina sequencing of parasitized diamondback moths (*Plutella xylostella*, Lepidoptera, Plutellidae) larvae also revealed that IV DNA circles from *Diadegma semiclausum* (DsIV) undergo massive integration into host hemocyte genomes during parasitism (Ze-hua Wang *et al*. 2021). Four DNA circles were found to integrate via a HIM motif, as observed for BV, but contrary to BV DNA circles, IV circles devoid of HIM were also found to integrate at substantial levels through DNA double strand breaks occurring at varying locations along the circles (Ze-hua Wang *et al*. 2021). The finding that both IV and BV DNA circles integrate via a shared HIM-mediated mechanism despite their independent origin is remarkable and several scenarios have been proposed to explain the possible evolutionary ties between these two polydnavirus lineages (Muller *et al*. 2021; Ze-hua Wang *et al*. 2021).

In the present study, we first apply the approach developed in Muller *et al*. (2021) on the *C. typhae*/*S. nonagrioides* system to comprehensively characterize and quantify DNA circle integration of yet another polydnavirus, the Hyposoter didymator IV (HdIV), in two tissues of parasitized fall armyworms (*Spodoptera frugiperda*, Lepidoptera, Noctuidae). Our results further underline the striking similarities in integration patterns between IV and BV, despite that these domesticated endogenous viruses are derived from viruses belonging to different families. We then assess whether HIM-mediated IV and BV chromosomal integration occurred in the germline of their hosts in the past by screening the genome of 775 lepidopteran species. We found sequences from IV and BV DNA circles in no less than 124 species belonging to 15 different lepidopteran families, suggesting that HIM-mediated wasp-to-host horizontal transfer of PDV DNA circles occurred recurrently during the evolutionary history of lepidopterans.

## Results

### Quantifying *Spodoptera frugiperda* and *Hyposoter didymator* genomic material

In order to characterize genome-wide patterns of HdIV integration during parasitism of the fall armyworm (*S. frugiperda*), we Illumina-sequenced whole DNA extracted from fat bodies and hemocytes of parasitized larvae. We obtained from 321 to 403 million trimmed reads depending on the tissue and time post-parasitism (hemolymph 24h and 72h [H24 and H72], fat body 24h and 72h [FB24h and FB72h]) (Table 1).

**Table 1:** Summary of the number of reads and chimeric reads obtained from *Spodoptera frugiperda* larvae parasitized by *Hyposoter didymator*.

|  | Accession number (SRA) | Total number of trimmed reads | Reads mapped on caterpillar genome | Number of chimeric reads | RPM on caterpillar genome | Number of IEs |
| --- | --- | --- | --- | --- | --- | --- |
| H24h | **SRX15279414** | 371 695 498 | 320038242 (86%) | 2 861 | 9 | 2 449 |
| H72h | **SRX15279415** | 321 231 671 | 263127889 (82%) | 4 903 | 19 | 4 398 |
| FB24h | **SRX15279416** | 403 147 682 | 346759613 (86%) | 2 516 | 7 | 2 065 |
| FB72h | **SRX15279417** | 347 218 922 | 299552940 (86%) | 3 325 | 11 | 2 908 |

H: hemolymph; FB: fat body. Numbers between brackets are percentages of total sequencing reads mapped on each reference genome. IEs: Integrations events (host-wasp junctions with the same coordinate in the host genome but covered by more than one chimeric read are counted as one integration event).

The majority of these reads (from 77% to 81%) aligned to the *S. frugiperda* genome (384 Mb), which was covered almost entirely (98%) in all samples. The average sequencing depth on the *S. frugiperda* genome varied from 101X for the H72h sample to 132X for the FB24h sample (Figure 2). By contrast, only 0.3% to 15% of the 226-Mb *H. didymator* genome was covered depending on the sample, at very low average sequencing depths (1 to 2X), most likely corresponding to the DNA of *H. didymator* eggs. By contrast, the average sequencing depth of the *H. didymator* wasp regions annotated as ichnovirus proviral segments by Legeai *et al*. (2020) was much higher than the rest of the wasp’s genome: 914X and 631X in the hemolymph (24h and 72hpp) and 1,220X and 386X for the fat body (24h and 72hpp) (Figure 2). This higher depth is consistent with the presence of many integrated and/or non-integrated forms of HdIV circles in the caterpillar after parasitization, since large amounts of virus particles are introduced with the parasite egg.

**Figure 2:**
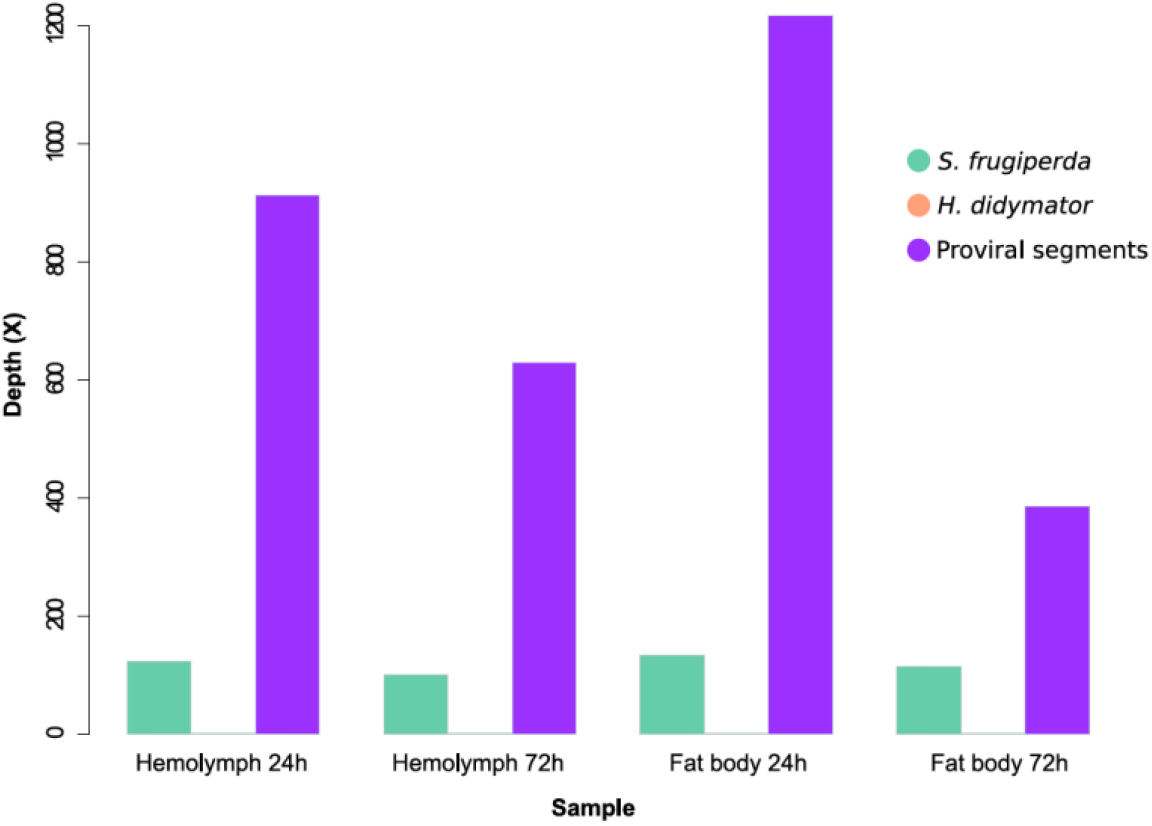
Mean sequencing depth over parasitized host (*Spodoptera frugiperda*) and parasitoid wasp (*Hyposoter didymator*) genomes.

### Quantification and characterization of integrated HdIV into *Spodoptera frugiperda* somatic genomes

To quantify chromosomal integration of HdIV circles into parasitized *S. frugiperda* cells, we searched for chimeric reads containing HdIV–*S. frugiperda* junctions (see methods). Depending on the sample, we detected from 2,516 to 4,903 chimeric reads (Table 1). Even after discarding PCR duplicates, a same integration event (IE), as defined by a given set of coordinates in the HdIV and *S. frugiperda* genome, might be covered by more than one chimeric read. This may occur when a given IE is duplicated through cell division. We thus estimated the number of independent IEs by counting as one event all reads with identical or nearly identical coordinates for both genomes. We found that the vast majority of IEs were covered by only one read and in total, we counted between 2,065 and 4,398 IEs (Table 1; Figure 3). The number of HdIV junctions covered by more than one reads went from 1 to 121 depending on the tissue (highest number of reads covering a junction = 3).

**Figure 3:**
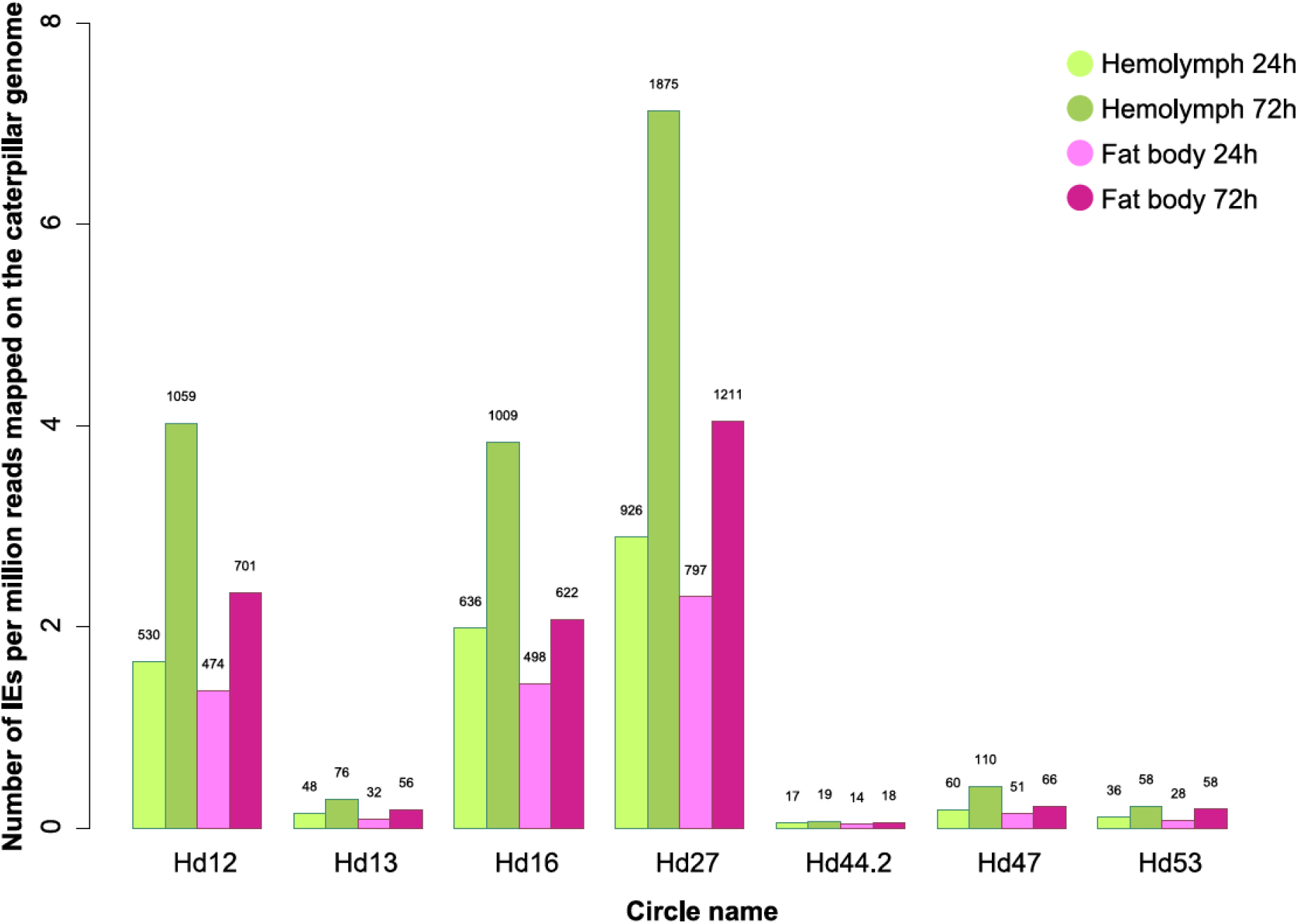
Number of HdIV chromosomal integration events per HdIV circle in parasitized *Spodoptera frugiperda* larvae. IE: Integration events (host-wasp junctions with the same coordinate in the host genome but covered by more than one chimeric read are counted as one integration event)

We found that seven out of the 57 HdIV circles had more than 10 chimeric reads mapping to them in all four DNA samples, strongly suggesting these seven segments underwent chromosomal integration. The seven segments form two groups depending on whether they have generated more or less than one IE per million reads mapped on the host (*S. frugiperda*) genome (respectively group 1: Hd12, Hd16 and Hd27 and group 2: Hd13, Hd44.2, Hd47 and Hd53) (Figure 3). Comparing normalized IEs per segment revealed a higher number of IEs in the hemolymph than in the fat body and a higher number of IEs at 72h than at 24hpp. These trends hold for all seven integrated segments and are in agreement with earlier findings on BVs (Beck *et al*. 2007; Muller *et al*. 2021).

Next, we followed the rationale exposed in Muller *et al*. (2021) and assessed how many IEs, on average, occurred per sequenced haploid genome. To do so, we divided the relative number of IEs (IEs per million reads mapped to the host genome) for each integrated circle by the read length and we multiplied it by the *S. frugiperda* genome size. We counted a total of 18, 40, 13 and 23 IEs per cell, on average, in the Hemocyte 24h and 72h, and the Fat body 24h and 72h samples, respectively.

### Mechanisms underlying integration of HdIV circles

To assess whether HdIV circle integration involves recombination within HIM as observed for BVs, and/or other mechanisms, we generated plots of chimeric read depths along all seven circles found integrated into *S. frugiperda* genomes (Figure 4; Supplementary Figure 1). The vast majority of chimeric reads (96.3 to 100% depending on the circle) map to a narrow region (in white on Figure 4) and most of the positions along each HdIV circle are not mapped by any chimeric reads. Depending on the circle, the length of the mapped region is between 92 and 120 bp. We arbitrarily delineated these specific regions in the HdIV circles following Muller *et al*. (2021), by identifying the two positions along the circles with the highest number of chimeric reads and by selecting 30 bp upstream of the left-most one and 30 bp downstream of the right-most one. We then verified that these regions align to the recently described HIM motifs in various IVs (Ze-hua Wang *et al*. 2021) (Figure 4). Zooming on the HIM also confirmed the presence of two peaks of chimeric reads, as observed in BVs (Muller *et al*. 2021) and corresponding to the J1 and J2 motifs, at which double strand breaks most often occur during linearization and integration of the circles (Chevignon *et al*. 2018) (Figure 4; Supplementary Figure 1). Thus, our results show that as for BVs, most if not all chromosomal integrations of HdIV circles are mediated by ichnovirus HIMs.

**Figure 4:**
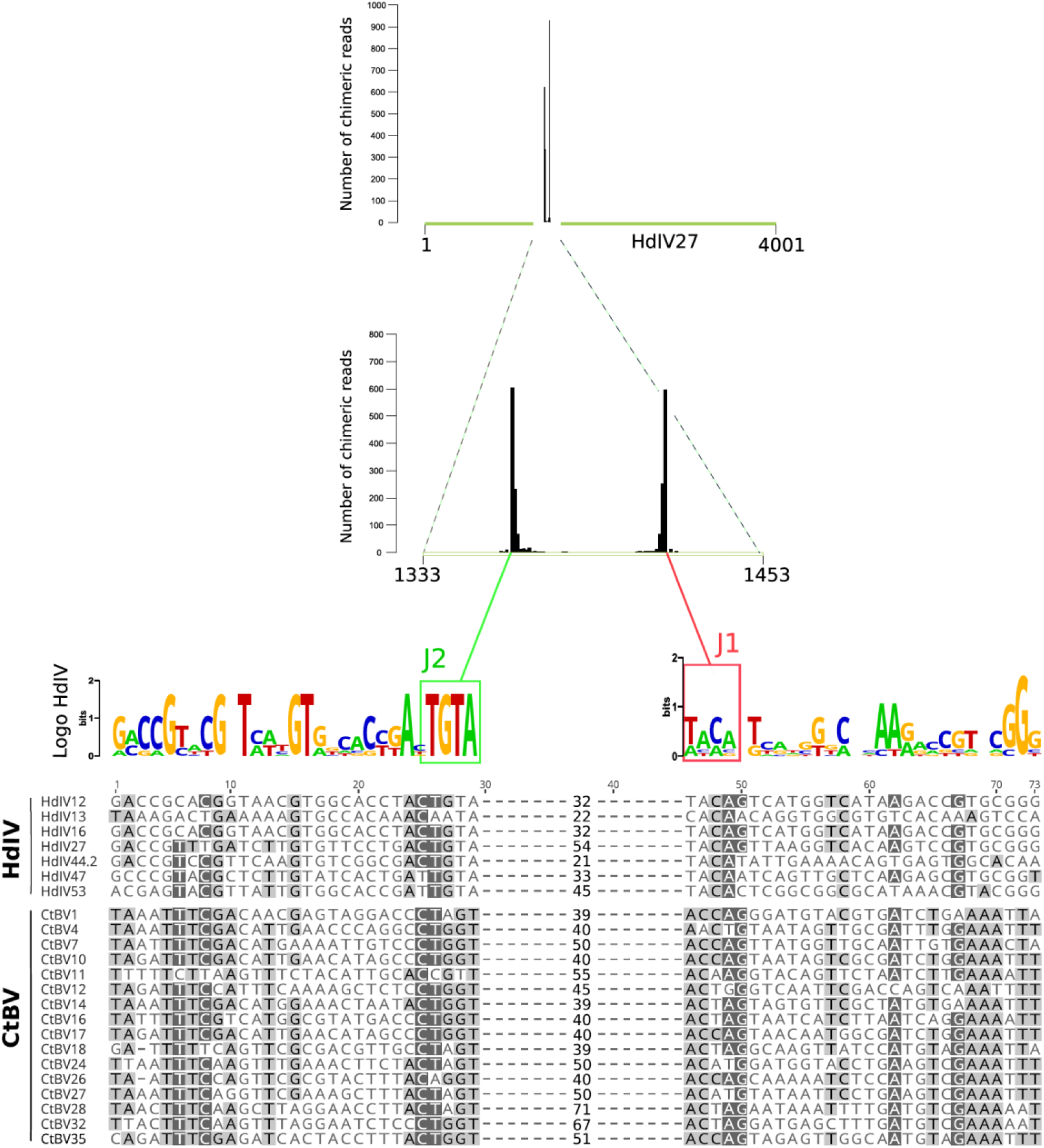
Polydnavirus DNA circles undergo HIM-mediated chromosomal integration. Top graph: map of chimeric reads along Hyposoter didymator ichnovirus circle 27 (Hd27) (green) in hemocytes showing that almost all chimeric reads map to the HIM (white). The circle is oriented from the 5’ DRJ to the 3’ DRJ. Below is a magnified view containing the 120-bp HIM, showing the two regions with many chimeric reads, called J2 (left) and J1 (right), corresponding both to the borders of Hd27 sequences integrated in parasitized host DNA. Below is shown the sequence logo of J2 and J1 from HdIV circles only (not including CtBV circles) generated with weblogo.berkeley.edu, and the alignment of the HIMs of the 7 HdIV circles that integrated into the *S. frugiperda* genome. An alignment of the CtBV circles described in Muller *et al*. (2021) is also shown below to allow comparison between bravovirus and ichnovirus HIM. Grey shading indicates the level of sequence conservation. Numbers in the middle of the alignment indicate the number of nucleotides that are present between the J1 and J2 motifs and that are lost upon integration of PDV circles in host genomes. This region is not conserved between circles and was thus removed to facilitate the reading of the figure.

To further characterize the mechanism involved in the integration within HIM regions and in particular whether homology between viral and host sequences could be involved in the integration process, we studied the distribution of microhomology lengths detected at the junction between HdIV and the *S. frugiperda* genome. We only investigated chimeric reads mapping to the HIM because the remaining chimeric reads represent only a very small fraction of all chimeric reads (0.8%). As previously done in our study on *Cotesia typhae* BV (Muller *et al*. 2021), we studied separately the major part of chimeric reads which map to the J1 and J2 motifs from those mapping outside of these motifs (but still within the HIM). We artificially generated chimeric reads following the method described in Peccoud *et al*. (2018) to compare the number of observed microhomology lengths with the number of expected microhomology lengths if junctions occurred randomly between HdIV HIMs and the *S. frugiperda* genome. For chimeric reads mapping to the J1 or J2 motifs, the analysis revealed a strong excess of 2- and 3-bp microhomologies and a slight excess of 4-bp microhomologies (Figure 5). The pattern was similar for chimeric reads mapping outside J1 and J2, though the excess of 3-bp microhomologies is less marked and that of 4-bp microhomologies no longer present. Interestingly, we observe a strong depletion of 0- and 1-bp microhomologies compared to the pattern expected by chance. This indicates that junction involving blunt-ended host-wasp sequences or 1-bp microhomology between them almost never occur.

**Figure 5:**
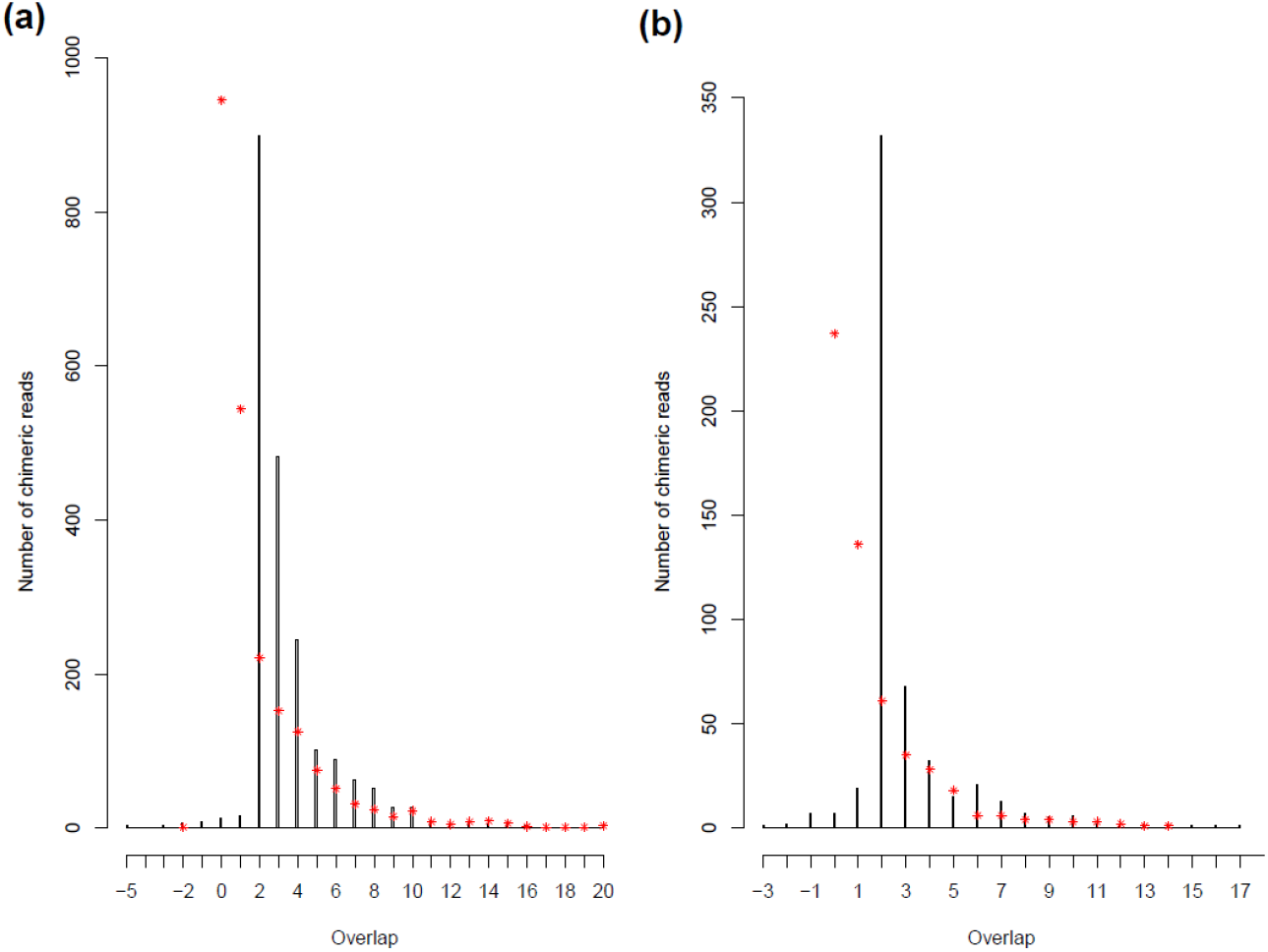
Distribution of microhomology lengths at wasp-host junctions in chimeric reads. Black bars correspond to the numbers of observed chimeric reads for each microhomology length. Red asterisks correspond to the expected numbers of chimeric reads for each microhomology length. (a) Distribution of microhomology lengths for HdIV-host junctions mapped in J1 or J2. (b) Distribution of microhomology lengths for HdIV-host junctions mapped within HIM but outside J1 or J2.

### Persistence of integrated versus non-integrated circles

We evaluated the quantity of integrated versus non-integrated HdIV circles during parasitism, as approximated by sequencing depth. We found that except for circle Hd27 which showed a very high sequencing depth (e.g., >6,000X in hemolymph 24h p.p.), the ranges of sequencing depths of integrated HdIV circles were similar to that of non-integrated HdIV circles (Figure 6). For example, in hemolymph 24h p.p., integrated circles (in blue on Figure 6) were sequenced at depths varying between 243X and 3400X while non-integrated circles (in red) were sequenced at depths varying between 93X and 4164X. At the later timepoint during parasitism (72h p.p.), the overall quantity of circles (integrated and non-integrated) decreases, with for example about three times less circles in the fat body at 72h p.p. than at 24h p.p (Figure 2). Interestingly, this decrease is overall stronger for non-integrated circles than for integrated ones.

**Figure 6:**
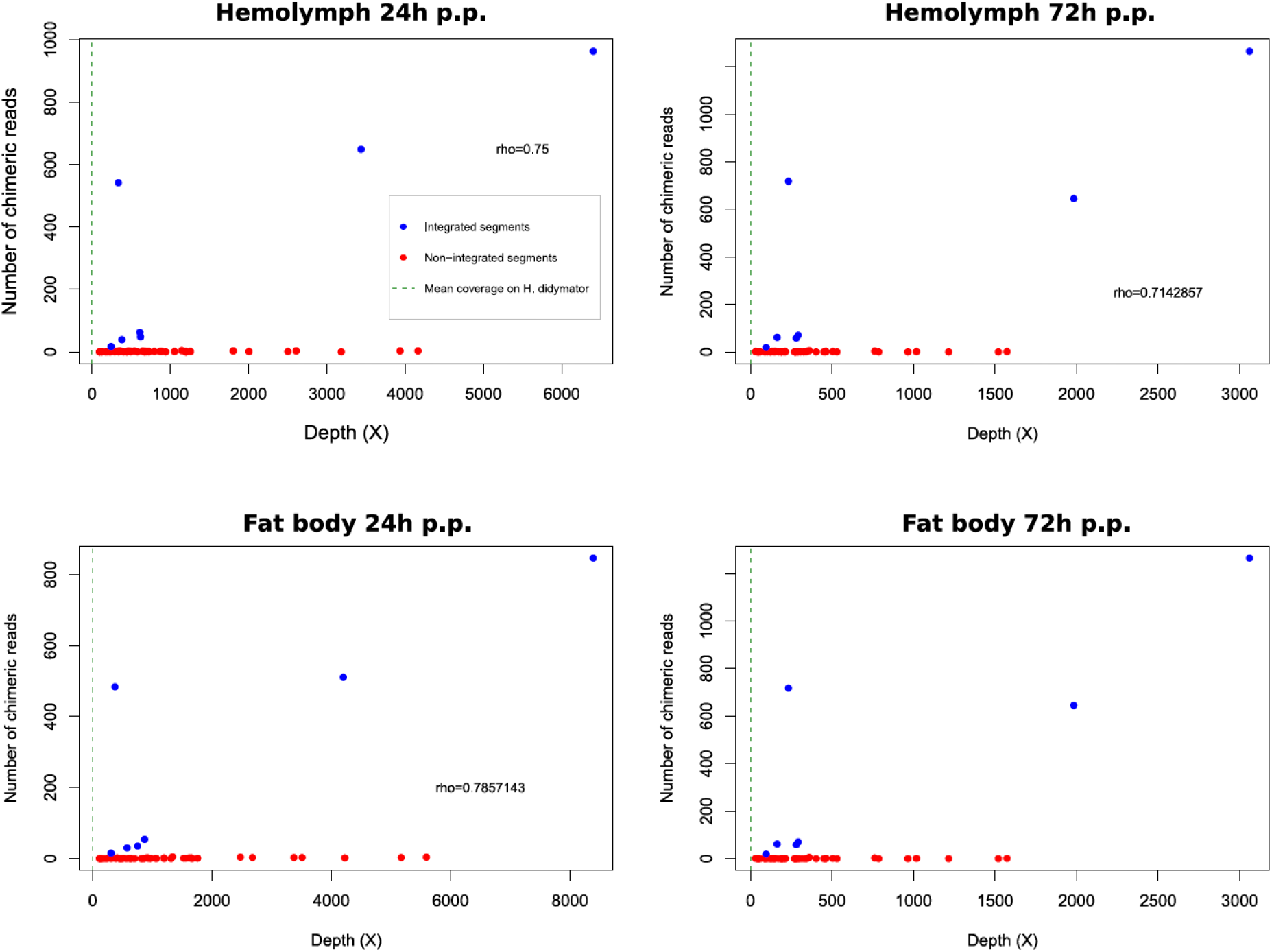
Plot of the sequencing depth versus the number of chimeric reads for each of the HdIV circle in hemolyphm 24h and 72h post parasitism (p.p.), and in fat body 24h and 72h p.p. Blue dots represent circles that do integrate into the S. frugiperda genome, and red dots represent circles that do not integrate. The identification numbers of the segments are shown near each blue dot. The vertical green dashed line shows the average sequencing depth over the H. didymator genome. The Spearman rho value indicates the correlation between sequencing depth and the number of integration events for circles that do integrate into the *S. frugiperda* genome.

This is well-illustrated by the fact that sequencing depth over Hd16 is lower than that of some non-integrated circles at 24h but it becomes higher than non-integrated circles at 72h p.p., a trend holding in both hemolymph and fat body (Figure 6). The capacity to integrate for a circle may thus increase its ability to persist in large amount during parasitism.

### Distribution of wasp circles throughout the genome of *Spodoptera frugiperda*

We investigated whether the integrations of *H. didymator* viral circles occur randomly along the caterpillar genome or whether there is an integration bias. To achieve this, we have split the *S. frugiperda* genome into 100,000-bp windows and assessed whether some windows were subject to more integrations than expected by chance. We found that the numbers of HdIV integrations per window did not follow a Poisson distribution with p-values <0.001 for all samples indicating that these integrations are not randomly distributed along the *S. frugiperda* genome). Under the Poisson distribution, we do not expect any windows with more than 6 or 7 IEs depending on the samples. However, for example, we detected in hemolymph samples a window with 14 IEs (H24h) and another with 18 (H72h) IEs. These results suggest that segments do not integrate completely randomly in the caterpillar genome, where IEs seem to be slightly concentrated in specific genomic regions.

### HIM-mediated integrations of polydnavirus DNA circles in multiple lepidopteran genomes

This analysis together with that of Wang *et al*. (2021) on DsIV, that of Wang *et al*. (2021) on CvBV and our earlier study on CtBV (Muller *et al*. 2021) shows that PDV circles undergo massive HIM-mediated chromosomal integration in host tissues during parasitism. We thus sought to assess whether horizontal transfer of PDV circles occurred from wasp to lepidopterans through HIM-mediated chromosomal integration in the germline of lepidopteran hosts, followed by vertical transmission in host populations. For this, we used HIM-containing HdIV and CtBV circles to perform blastn similarity searches on 775 lepidopteran genomes available in Genbank as of December 2021. Our search yielded 4,648 and 49,079 lepidopteran sequences longer than 300 bp that showed similarity to HIM-containing HdIV and CtBV circles, respectively, with an e-value lower than 0.0001. We reasoned that if some of these sequences were integrated through double strand breaks within HIM, then they should have been initially bordered by the J1 and J2 motifs (Figure 1) and may have since conserved these extremities, or at least one of them (J1 or J2). Moreover, the region between J1 and J2 which is lost during integration should not be retrieved during the analyses (see above and Chevignon *et al*. 2018 and Muller *et al*. 2021). In agreement with these expectations, no less than 2,213 of the 4,648 IV sequences were found in lepidopteran genomes starting or ending within the HIM, and none of them contained the entire region lying between the J1 and J2 motifs. Regarding the 49,079 blastn hits involving CtBV, we found 174 sequences starting or ending within the HIM and only 3 containing the entire region lying between the J1 and J2 motifs. Our blastn searches did not allow us to directly retrieve full length PDV circles bordered by J1 on one side and J2 on the other side. This may be because integrations are ancient and were followed by rearrangements and/or degradation of the circles. Another reason behind the fragmentary nature of the blastn hits may be that the donor wasp species was distantly related to *H. didymator* or *C. typhae*. While complete, circles from such distant species may be too divergent to align over their entire length with no interruption to those of *H. didymator* or *C. typhae*. Importantly, we manually curated several lepidopteran PDV sequences and were able to reconstruct several examples of full-length integrated circles bordered by J1 and J2 (Supplementary Dataset 1). In the following, we refer to fragments of IV or BV sequences as retrieved by our blastn search, *i*.*e*., fragments longer than 300 bp and bordered either by the J1 or J2 motif, to describe patterns of HIM-mediated integration of PDV circles into lepidopteran genomes.

### Numbers and distribution of polydnavirus DNA circles in the lepidopteran tree

IV fragments bordered by J1 or J2 motif were found for four out of the seven HIM-containing HdIV circles in a total of 87 lepidopteran species (out of 775) that belong to 11 out of the 35 lepidopteran families included in our search (Supplementary Table 1; Figure 7). BV fragments bordered by J1 or J2 motif were found for 14 out of 16 HIM-containing CtBV circles in a total of 60 lepidopteran species belonging to 11 families, including 7 (Geometridae, Hesperiidae, Lycaenidae, Noctuidae, Nymphalidae, Pieridae, Riodinidae), which also had IV fragments integrated in their genome (Supplementary Table 2; Figure 7). Altogether, PDV fragments (IV + BV) were retrieved in a total of 124 species with 23 having both IV and BV in their genome. These 124 species are from 15 lepidopteran families, which belong to seven superfamilies (Noctuoidea, Geometroidea, Papilionoidea, Pyraloidea, Tortricoidea, Zygaenoidea, Yponomeutoidea, Gelechioidea) that diverged more than 100 million years ago (MYA), and cover a large diversity of lepidopterans (Figure 7) (Kawahara *et al*. 2019).

**Figure 7:**
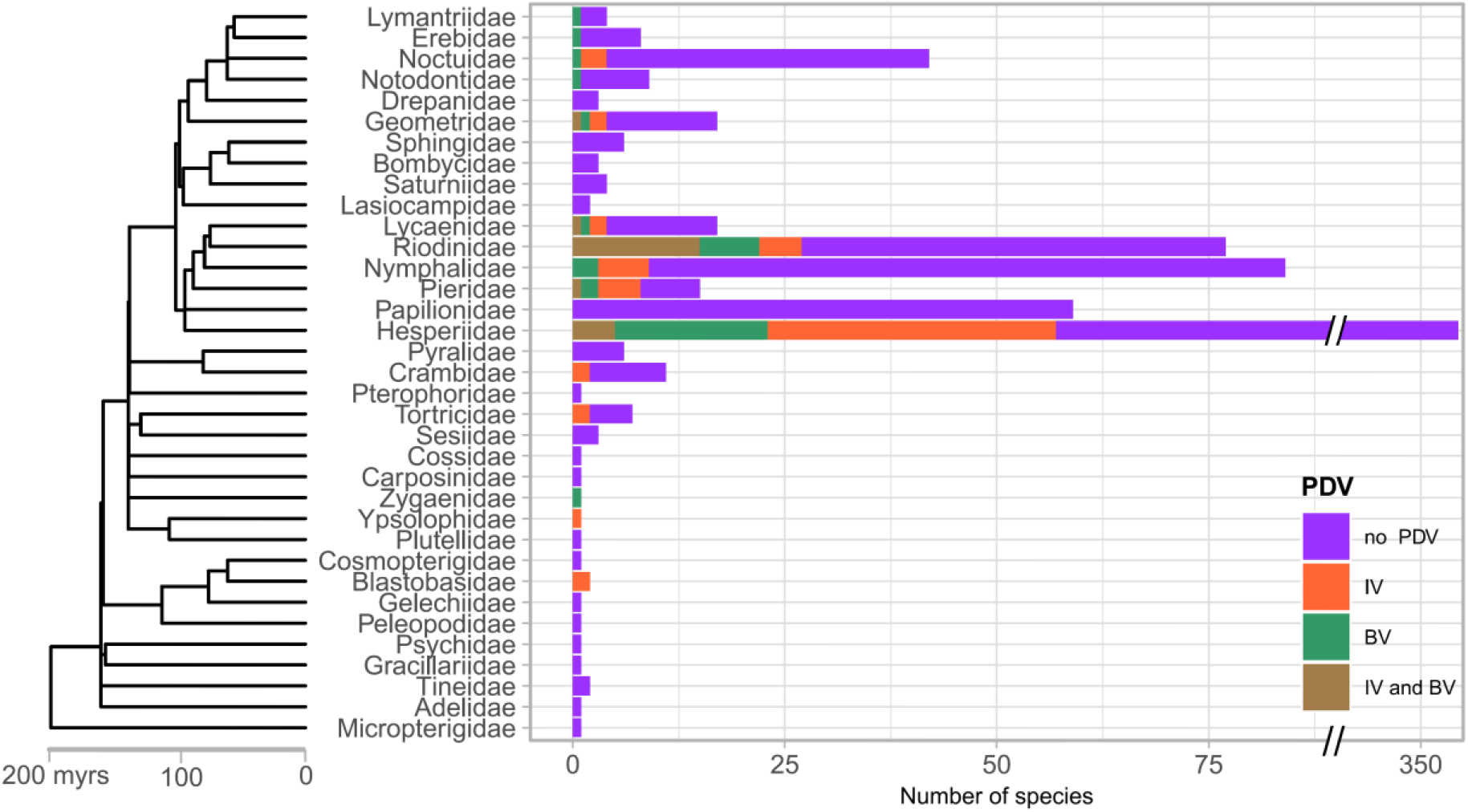
Numbers of genomes per lepidopteran families in which J1- or J2-bordered bracovirus (BV) and ichnovirus (IV) fragments were found. Details on the numbers and type of PDV circles in each lepidopteran species can be found in figure 8 and supplementary tables 3 and 4. The tree and divergence times were recovered from timetree.org (Kumar *et al*. 2017).

Lepidopteran IV fragments retained for this study are between 300 and 4110 bp (median = 621 bp) and show between 65 and 96% nucleotide identity to HdIV circles (median = 78.5%). BV fragments are between 304 and 2860 bp (median = 515 bp) and show between 65 and 88% nucleotide identity to CtBV circles (median = 77%). All PDV fragments bordered by J1 or J2 motifs as well as their alignment coordinates and percent identity to CtBV or HdIV are provided in Supplementary Tables 1 and 2.

Numbers of PDV fragments are heterogenous both in terms of circles and lepidopteran species. In terms of lepidopteran species, numbers of IV fragments are generally relatively low, with 52 species having only one or two fragments and 76 out of the 87 species having less than 10 (Supplementary Table 3). However, three species have high (81 in *Blastobasis adustella* and 80 in *B. lacticollela*, Blastobasidae) to very high (1756 in the wainscot hooktip *Ypsolopha scabrella*, Ypsolophidae) numbers of IV fragments (Supplementary Table 3, Figure 8). Numbers of BV fragments are also generally relatively low, with 48 out of the 60 species having only one or two J1- or J2-bordered BV fragments (Supplementary Table 4). Four species have ten or more (up to 17) BV fragments (*Calephelis nemesis, C. perditalis, Apodemia duryi* and *Boloria selene*) (Supplementary Table 4, Figure 8). The number of both IV and BV fragments per lepidopteran family was positively correlated with the number of genomes surveyed per family (both Pearson’s r = 0.98 ; p-values < 0.000001).

**Figure 8:**
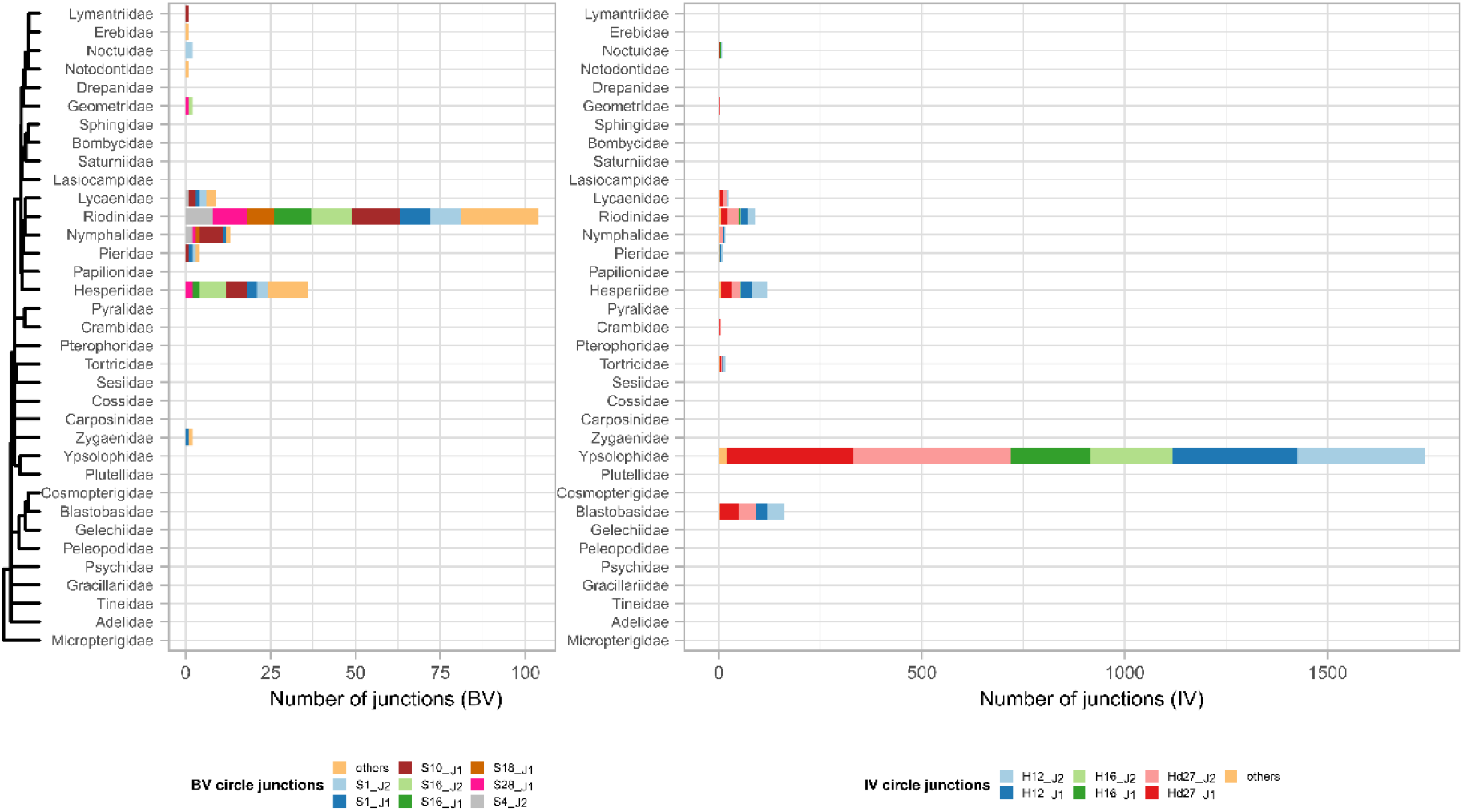
Numbers of J1- or J2-bordered bracovirus (BV) and ichnovirus (IV) fragments per lepidopteran families. Details on the numbers and type of PDV circles in each lepidopteran species can be found in supplementary tables 3 and 4. Junctions types for which less than 8 (BV) or 10 (IV) fragments were found in all families were grouped together in the “others” category.

In terms of circles, we found a much higher number of IV fragments corresponding to Hd12/16 (634 J1 and 595 J2) and Hd27 (498 J1 and 422 J2) than to Hd47 (28 J1 and 35 J2) and Hd44.2 (1 J2) (Figure 8; Supplementary Table 3). This mirrors the pattern observed for somatic integrations, with Hd12, Hd16 and Hd27 being highly integrated and Hd47 being the mostly highly integrated among lowly integrated circles (Figure 3). Numbers of IV J1 and J2 junctions are similar, which is consistent with the fact that during HIM-mediated integration, each integrated IV circle is linearized through double-strand break within the HIM and thus ends flanked by the J1 motif on one side and J2 motif on the other side (Figure 1). In terms of BV circles, the number of fragments found in lepidopteran genomes did not match with the relative abundance of integrated circles in parasitized hosts. For example, while we found the highest number of BV fragments for CtBV circles 16 (21 J2 and 13 J2) and 10 (31 J2) (Supplementary Table 4, Figure 8), these two circles are among the least integrated circles in somatic genomes of parasitized *S. nonagrioides* individuals (Figure 6 in Muller *et al*. 2021).

CtBV circle 1, which is by far the most highly integrated circle in parasitized host tissues is among the most integrated circles in lepidopteran genomes but the number of C1 fragments (16 J1 and 17 J2) are close to those of several other circles (Supplementary Table 4, Figure 8). Contrasting with IV, for which similar amounts of J1-bordered and J2-bordered fragments were recovered for each circle in lepidopteran genomes, we often found different numbers of J1 and J2 BV fragments. In fact, we found only J1-bordered fragments (no J2-bordered fragments) for 9 BV circles (Supplementary Table 4, Figure 8). We believe that this disequilibrium is unlikely to have biological underpinnings because when we lowered the length threshold used to filter blastn hits to 200 bp, *i*.*e*. when we retain all BV-like sequences longer than 200 bp instead of 300 bp, the number of J1- and J2-bordered BV fragments are more similar for some circles (not shown). Yet we chose to keep the 300 bp threshold to retain blastn hits in order to maximise specificity and to ensure recovering large enough PDV fragments to conduct phylogenetic analyses.

### Sequencing depth supports germline integration of polydnavirus DNA circles

Several features of the lepidopteran PDV fragments strongly suggest that they do not result from contamination or from chromosomal integrations in somatic genomes of parasitized individuals. Polydnavirus DNA circles are known to occur in three forms: 1) proviral segments flanked by DRJs in the genome of wasps, in which J1 and J2 motifs are next to each other, separated by a short sequence (about 50 bp) in the HIM, 2) circular sequences in the wasp ovaries and in host larvae, in which J1 and J2 motifs are also next to each other in the HIM, and 3) sequences bordered by J1 and J2 motifs integrated in parasitized hosts’ somatic genomes. The fact that PDV fragments we report here in lepidopteran genomes are bordered at one of their extremity by the J1 or J2 motif, unlike in the wasp genome in which these motifs lie next to each other, argues against the possibility that these fragments result from contamination by wasp DNA that could have occurred during DNA extraction and/or sequencing. However, these fragments could correspond to somatic integrations that could have occurred in the individuals that were used for whole genome sequencing (many lepidopteran genomes are obtained from a single individual). This would imply that these individuals were parasitized by a wasp prior to sequencing. An important difference between somatic and germline integrations of PDV circles is that while the former are present only in a subset of somatic cells, the later should be present in all cells of an individual that would have received them from its parents. In agreement with the presence of PDV IEs in a subset of somatic cells during parasitism, 98% of the CtBV IEs we identified in parasitized *S. nonagrioides* larvae were covered by only one chimeric read (Muller *et al*. 2021). The pattern is similar here for HdIV, with between 97.2 and 99.9% of HdIV IEs covered by only one read, depending on the sample. In contrast with somatic IEs, germline integrations should be covered by multiple reads because they are expected to be present in all cells of the sequenced lepidopteran individual. Furthermore, sequencing depth over the integration should be similar between the PDV and lepidopteran flanking genomic region. To verify this, we recovered raw Illumina reads for the eight lepidopteran species having ten or more IV fragments, and for the six species having five or more BV IEs, and we computed average sequencing depth and number of chimeric reads for all IV or BV junctions in each species. In agreement with our predictions, we found that the average number of chimeric reads per species was always higher than one (Figure 9 and 10). In fact, it was higher than five for five out of six species harbouring BV fragments (Figure 10) and higher than 10 for seven out of eight species harbouring IV fragments (Figure 9). Importantly, the average number of chimeric reads per IE was always close to the average sequencing depth at the PDV integration point. Thus, variations in numbers of chimeric reads between species have no biological underpinnings but are due to variation in sequencing depth. Furthermore, we found that for most species, average sequencing depth per species was homogeneous across PDV/lepidopteran junctions (Figure 9 and 10).

**Figure 9:**
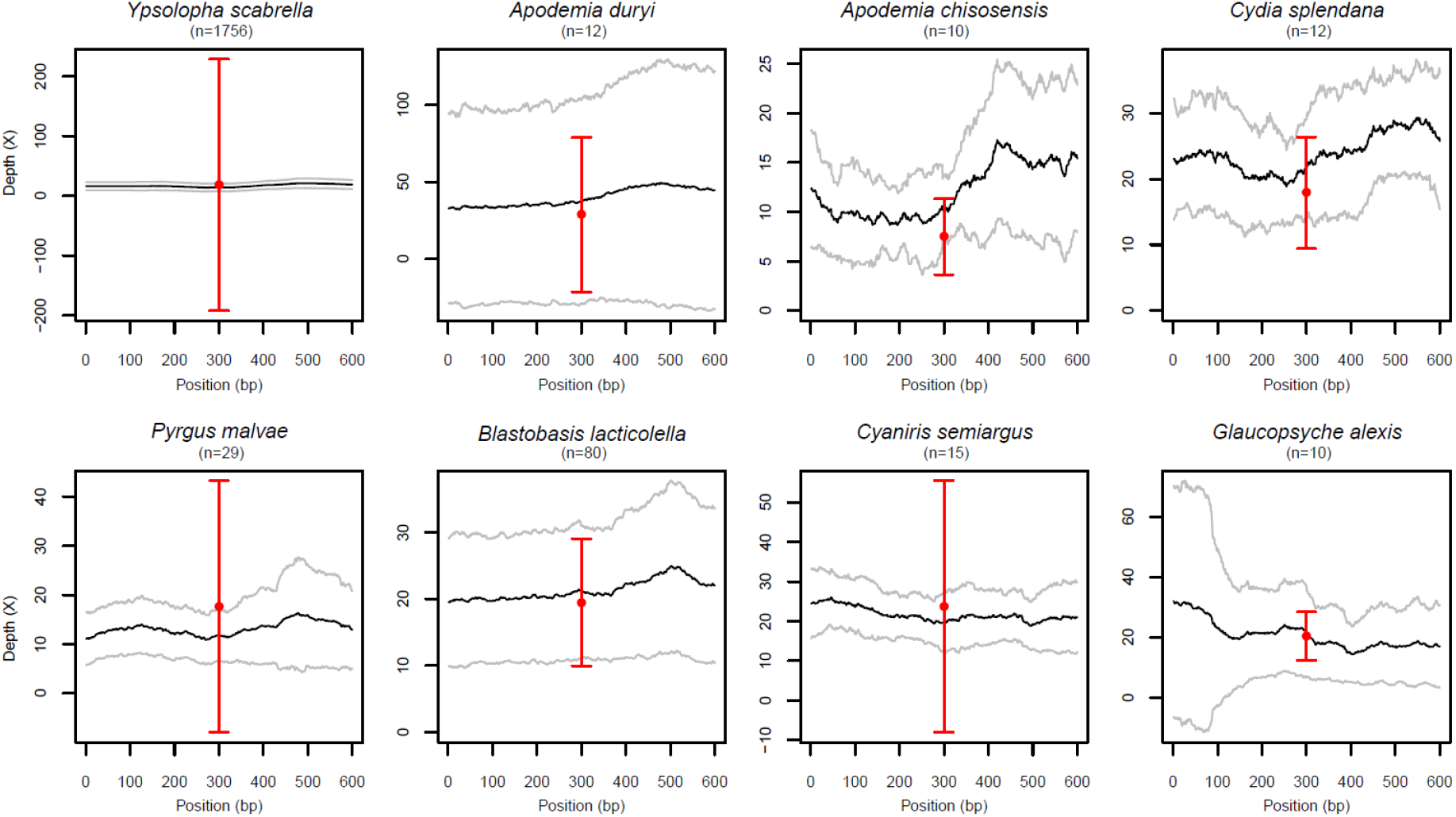
Sequencing depth and numbers of chimeric reads support HIM-mediated chromosomal integration of ichnovirus fragments in the germline of lepidopteran species. Eight lepidopteran species in which 10 or more J1- or J2-bordered ichnovirus fragments were found were selected to compute average sequencing depth over all junctions. The X axis indicates the genome position along the junction, the ichnovirus fragments being located between position 1 and 300 bp and the flanking lepidopteran genome regions being between 301 and 600 bp. The black line indicates average sequencing depth over 300 bp upstream and downstream of the junctions, with grey lines indicating standard deviation. Red dots indicate average numbers of chimeric reads supporting junctions in each species. Standard deviation is also shown.

**Figure 10:**
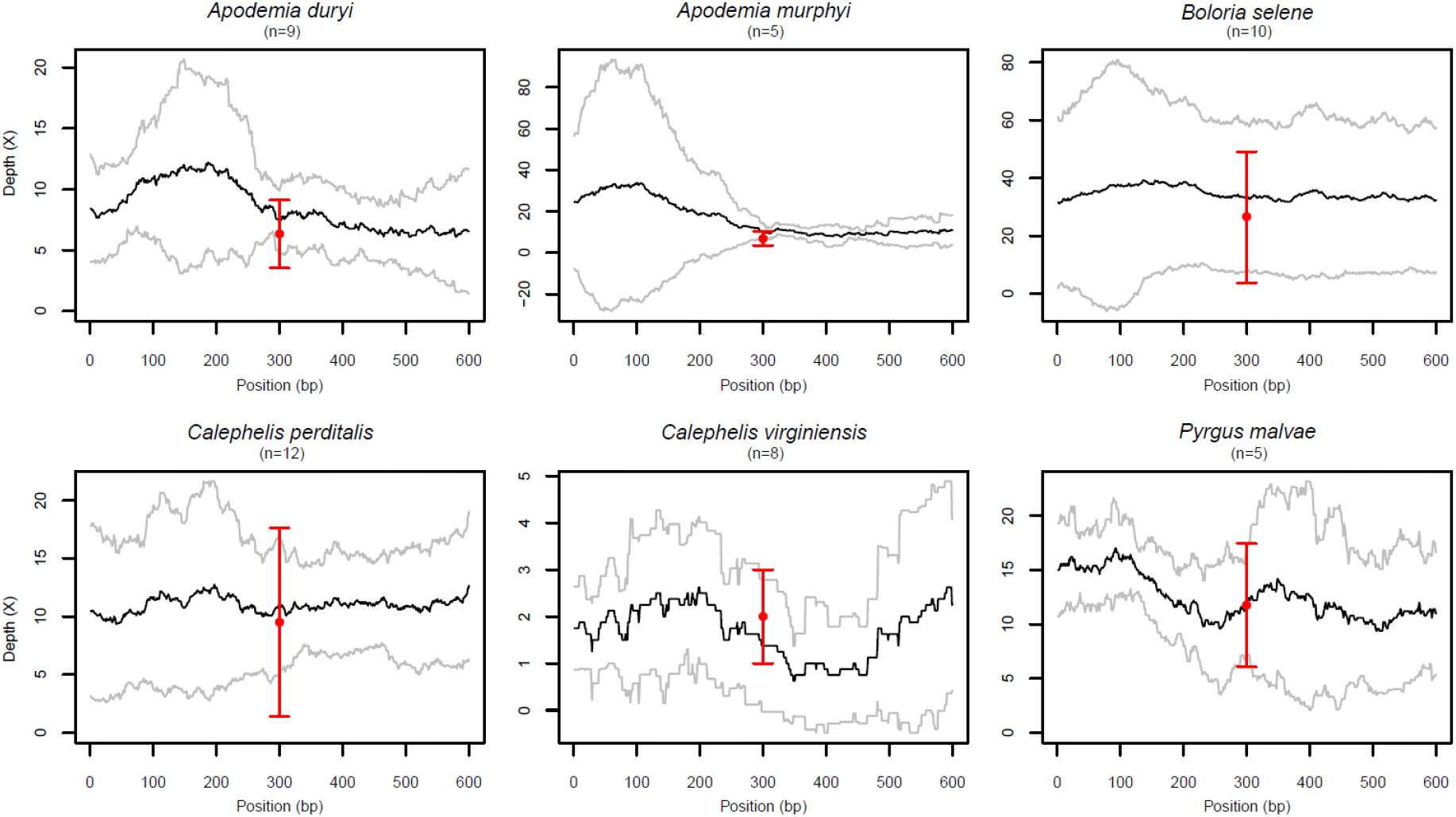
Sequencing depth and numbers of chimeric reads support HIM-mediated chromosomal integration of bracovirus fragments in the germline of lepidopteran species. Six lepidopteran species in which 5 or more J1- or J2-bordered bracovirus fragments were found were selected to compute average sequencing depth over all junctions. The X axis indicates the genome position along the junction, the bracovirus fragments being located between position 1 and 300 bp and the flanking lepidopteran genome regions being between 301 and 600 bp. The black line indicates average sequencing depth over 300 bp upstream and downstream of the junctions, with grey lines indicating standard deviation. Red dots indicate average numbers of chimeric reads supporting junctions in each species. Standard deviation is also shown.

For two species (*Apodemia duryi* and *A. murphyi*), we noticed a particularly high increase in average sequencing depth over BV circles compared to the flanking lepidopteran genomic sequence (Figure 10), which could result from the presence of unintegrated BV circles in these species. However, reads produced by unintegrated circles are expected to cover such circles relatively homogeneously, which should here translate into a sharp increase in depth on the circle side starting from the very junction. Yet, here the increase in depth is progressive. We note that the quality of the assembly of the two species is low as both have N50 lower than 500 bp. It is thus possible that the increase in sequencing depth over the circle side in *Apodemia* species may be caused by the fact that several integrated circles have not been included in the assembly. Furthermore, the average number of chimeric reads in the two species (6.3 and 7.0) is consistent with the presence of most BV IEs being present in all cells. Thus, while we are unable to provide a definitive explanation for the higher depth on BV circles compared to flanks for these two species, our observations are not consistent with the presence of unintegrated circles. Overall, we contend that these results indicate that most J1/J2 bordered PDV fragments we found in lepidopteran genomes result from germline integration that were then transmitted vertically in host populations.

An independent confirmation of this reasoning was obtained by assessing experimentally the presence of J1 and J2 extremities of insertions related to the HdIV12 circle and to the Cotesia congregata bracovirus circles 1 and 17 in different individuals of two very common butterflies: the small white and the green vein white (*Pieris rapae* and *Pieris napi*). Twelves individuals of each species were collected from three different locations in France. We obtained specific bands by PCR amplification of the junctions between flanking and viral sequences for all individuals (Figure 11).

**Figure 11:**
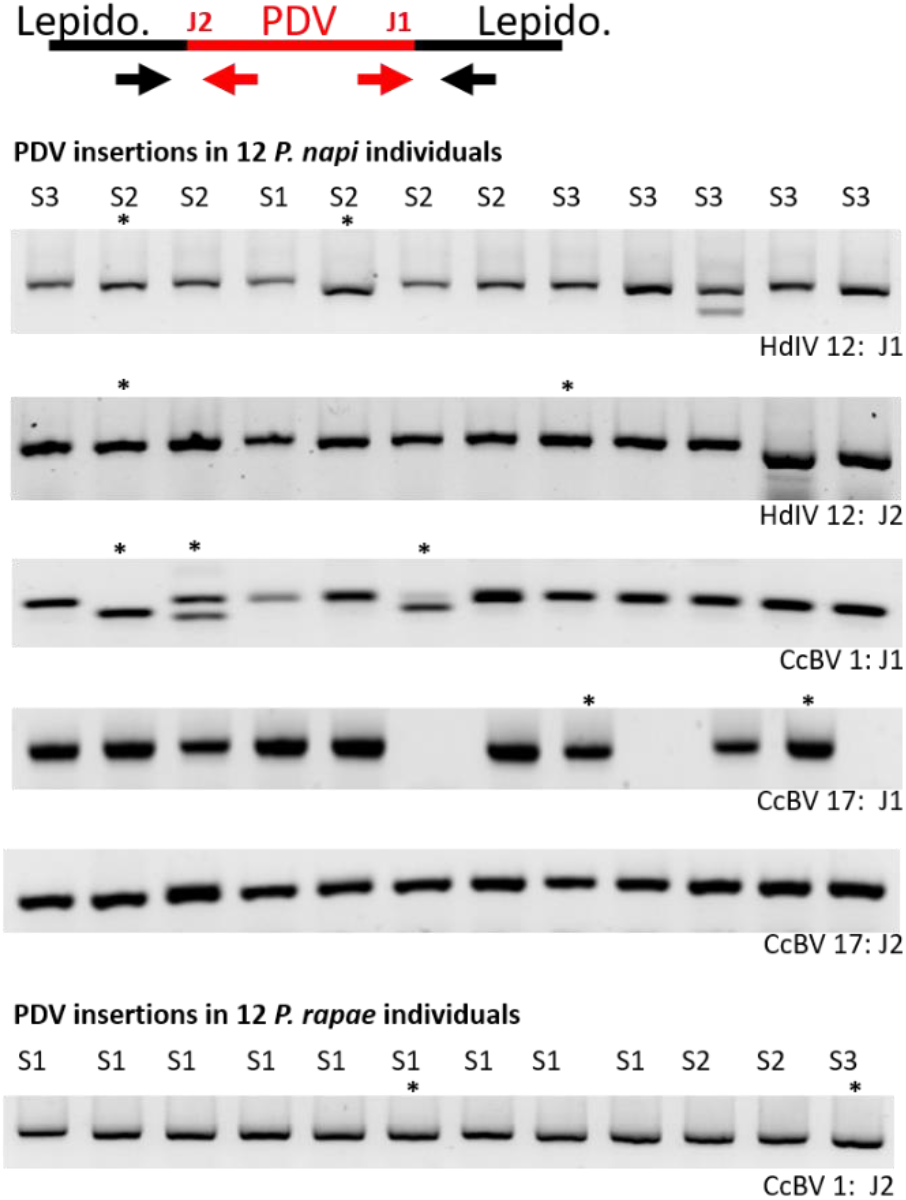
Experimental verification of polydnavirus-lepidopteran junctions in populations of two butterfly species (*Pieris napi* and *P. rapae*). The top illustration shows schematic a HIM-mediated insertion of a polydnavirus (PDV) circle bordered by the J1 and J2 motifs, as well as the position of the PCR primers (arrows at the bottom) designed on each side of the junction. Below are pictures of electrophoresis gels showing presence or absence of PCR bands obtained with primers targeting five PDV-*P. napi* junctions and one PDV-*P. rapae* junction in twelve individuals of each species sampled in three different geographical locations (S1, S2, S3, see methods). Asterisks indicate PCR products that were Sanger-sequenced. The sequences are provided in Supplementary Dataset 2. For each verified junction, the type of PDV circle is indicated at the bottom right of the gel. HdIV: Hyposoter didymator ichnovirus. CcBV: Cotesia congregata bracovirus. The genomic coordinate of each junction is provided in Supplementary Dataset 2. Note that polymorphism was observed in *P. napi* for CcBV1 J1 containing PCR products (2 individuals having two bands) and in CcBV17 J1 (amplification was reproducibly not obtained from the DNA of three individuals) probably reflecting a still ongoing erosion of these regions. CcBV1 insertions in *P. napi* and *P. rapae* have different flanking regions (See Supplementary Dataset 2).

Sequencing of the PCR products confirmed that J1 and J2 constituted actually the extremities of the four insertions identified from *P. rapae* and *P. napi* genomes, which thus appear to be fixed in the population and not a particular feature of the individuals used for genome sequencing. All sequences are provided in Supplementary Dataset 2. The *P. napi* HdIV 12 J2 sequence corresponds to the blastn hit number 167 in Supplementary Table 1. The other five verified junctions were not recovered in the bioinformatic search because this experimental study was performed independently by GP, JMD and KM in 2018. The study differed from the bioinformatic survey performed in 2022 in that it used CcBV as a query bracovirus instead of CtBV and it included two genomes of *P. napi* instead of one (see methods and Supplementary Dataset 2).

### Integration dynamics of polydnavirus fragments during lepidopteran evolution

Some PDV circle integrations are likely to be ancient and thus shared with other species as found for a 5-million years old bracovirus integrations shared by the monarch and related species of the Danaina subtribe (Gasmi *et al*. 2015). To gain insights into the timing of different integration events of PDV fragments in lepidopteran germlines, we first searched for PDV fragments that are orthologous between species, *i*.*e*., fragments located at the same position in the genome of two or more species. Our approach allowed us to identify five BV fragments and eight IV fragments shared by two or more species at the same genomic position (Table 2).

**Table 2.**
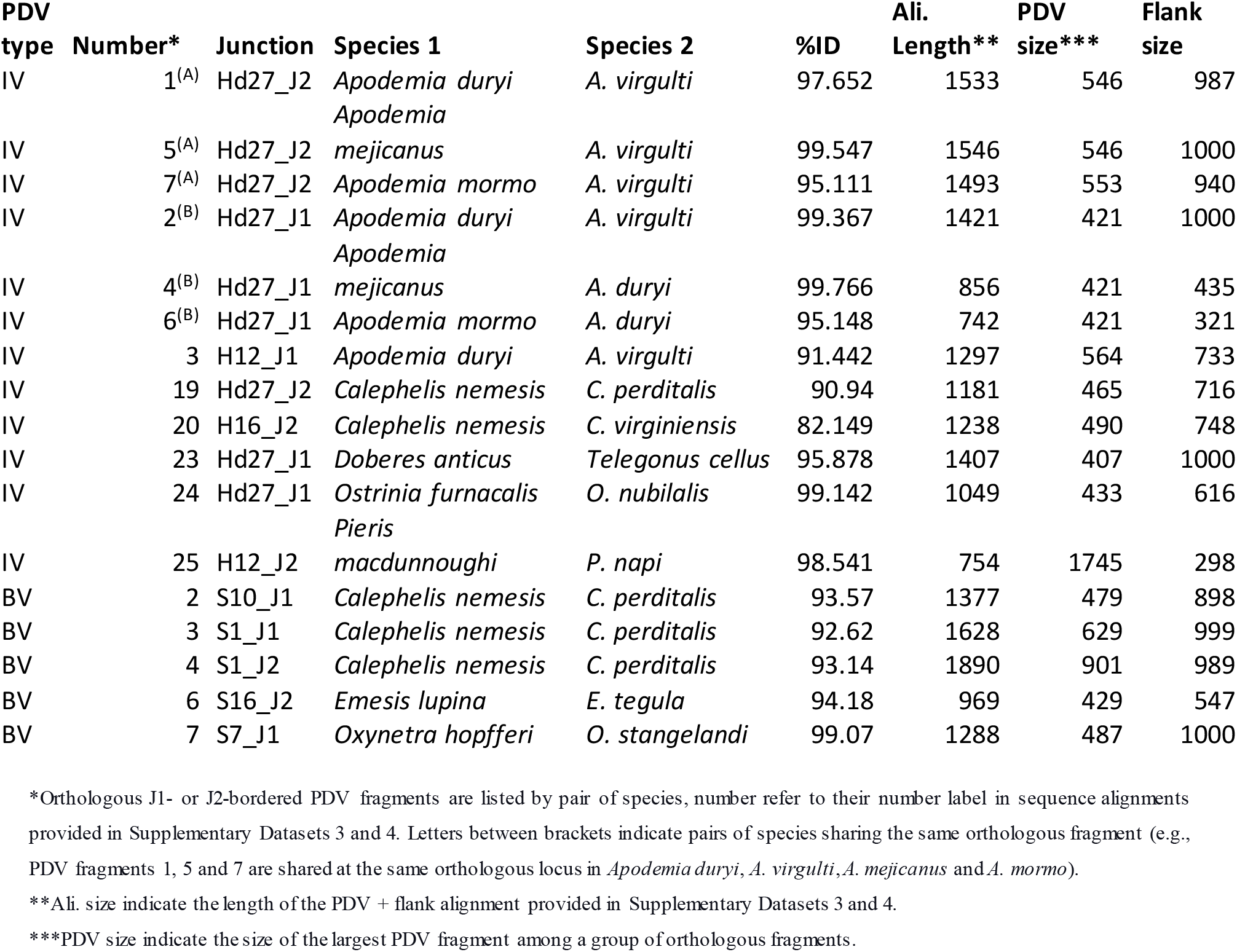
Characteristics of J1- or J2-bordered PDV fragments shared at orthologous loci between two or more lepidopteran species.

One IV orthologous fragment, found in *Calephelis* metalmarks may be as old as the monarch integration as it shows only 82.1% nucleotide identity between *C. nemesis* and *C. virginiensis* and the split between the two species has been dated at 4.9 MYA (Table 2, Supplementary Dataset 3) (Cong *et al*. 2017). By contrast, all other cases involve species for which divergence time is unknown but is likely recent because the species are congeneric and/or nucleotide identity between orthologous fragments is high (from 90.9 to 99.3%; mean = 96.6%). Regarding IV orthologous fragments, two are shared between four very closely related species of metalmark butterflies (genus *Apodemia*, Riodiniidae) (Zhang *et al*. 2019), one of them is shared between the green-veined white (*Pieris napi*, Pieridae) and its close relative *P. macdunnoughi* (Chew and Watt 2006) and another one is shared between the European corn borer (*Ostrinia nubilalis*, Crambidae) and the Asian corn borer (*O. furnacalis*), which diverged recently (Table 2; Supplementary Dataset 3) (Bourguet *et al*. 2014). In terms of BV orthologous fragments, three are shared between *C. nemesis* and *C. perditalis* metalmark butterflies, one is shared between two species of firetips (*Oxynetra hopfferi* and *O. stangelandi*) and one is shared between *Emesis lupina* and *E. tegula* (Table 2; Supplementary Dataset 4) (Cong *et al*. 2017).

In addition to orthologous fragments shared between species, we found several paralogous IV fragments sharing the same immediate flanking region genome (Supplementary Table 5; Supplementary Dataset 5). Numbers of such paralogous fragments vary from two in the Kamehameha butterfly *Vanessa tameamea*, (Nymphalidae) to 83 in the wainscot hooktip *Y. scabrella* (Ypsolophidae), with a maximum of three sequences in a given paralogous group (Supplementary Table 5). The finding of paralogous sequences sharing flanks indicates that some IV fragments were duplicated after integration. Most duplications are likely recent as identity levels within these paralogous groups are generally high (from 82.7 to 100%; mean = 97.2%; median = 99.2%).

We next generated multiple alignments of PDV fragments inserted in Lepidopteran genomes and PDV circles from various wasps, that we submitted to phylogenetic analyses. As expected in these phylogenies, PDV fragments found to be orthologous between two or more lepidopteran species group together. This is the case for example of the four Hd27 J1 fragments flanked by the same genomic region in four *Apodemia* species (Table 2; Figure 12). By contrast, when multiple PDV fragments were found in genomes of a given lepidopteran family, genus, or species, they generally do not cluster together in a monophyletic group (Figure 12 and Supplementary Figure 2). For example, Hd27 J1 fragments found in various Hesperiidae species (dark green in Figure 12) fall in at least 6 different clusters intermingled with fragments found in other lepidopteran families. Similarly, Hd27 J1 fragments from various species of the *Apodemia* genus, as well as those from the species *Ypsolopha scabrella* are polyphyletic, forming multiple clusters in the tree (Figure 12).

**Figure 12:**
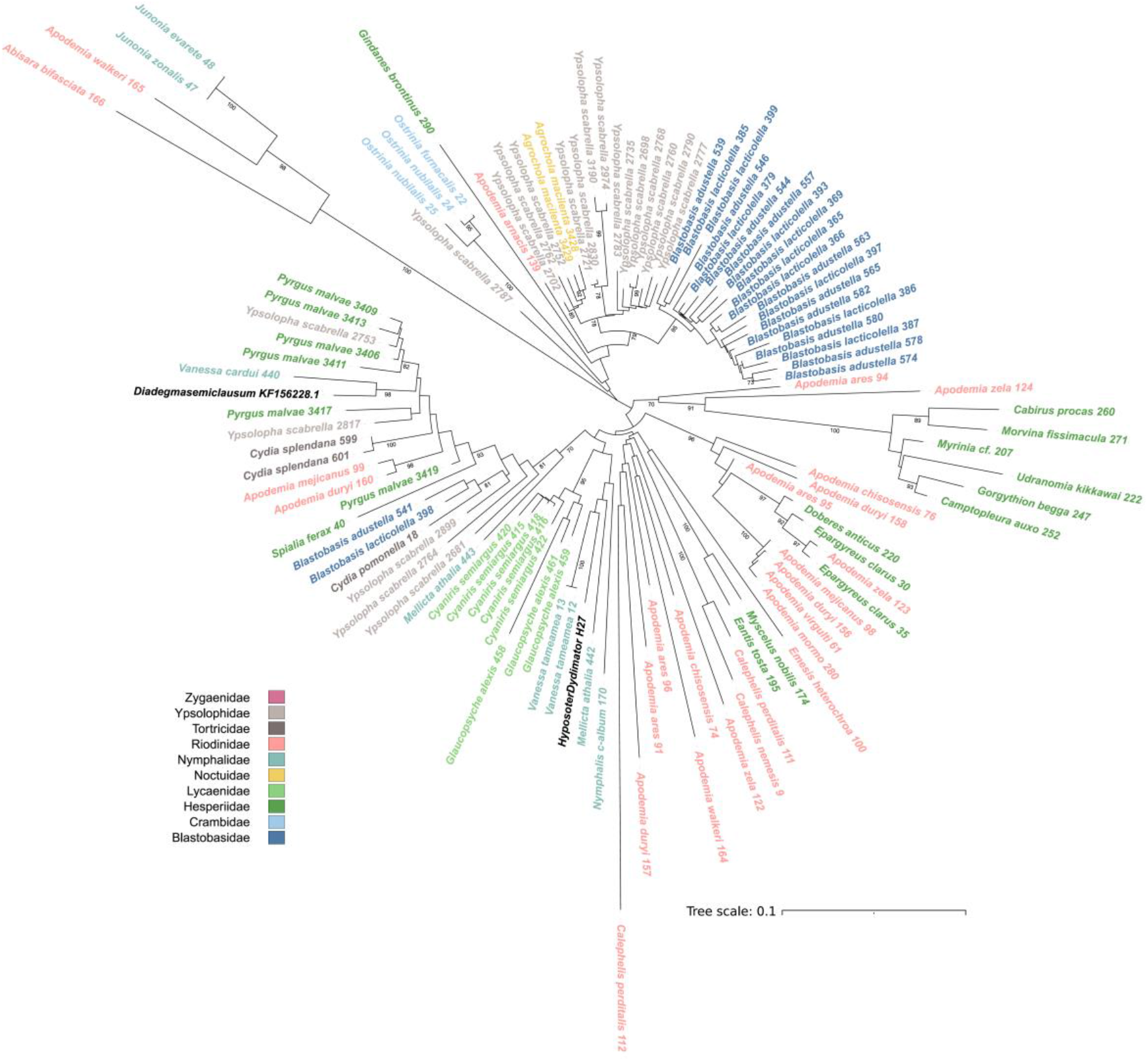
Unrooted phylogeny of IV fragments homologous to the J2 junction of the HdIV27 circle found in lepidopteran genomes. All fragments similar to the J2 extremity of the HdIV27 circle found in lepidopteran genomes were aligned together with HdIV27 and a homologous circle from *Diadegma semiclausum* (both are in black). Prior to alignment, the numerous fragments found in *Ypsolopha scabrella, Blastobasis adustella* and *Blastobasis lacticolella* were clusterized at 95% nucleotide identity threshold. Species names are coloured according to lepidopteran family and the number associated to each species name is a unique identifier that allows retrieving the sequence in Supplementary Table 1 (column “Number”). Numbers on branches are bootstrap values higher than 70%. The tree was built using the Maximum Composite Likelihood model of substitution available in MEGA 11 (Tamura *et al*. 2021) with 100 bootstrap replicates.

Furthermore, PDV sequences from different contemporary wasps are also generally scattered in the phylogenies. This is the case for example of the Hd27 J1 sequence from the wasp *Diadegma semiclausum*, which falls more closely related to lepidopteran Hd27 J1 than to the Hd27 J1 sequence of *H. didymator* (Figure 12). Similarly, H12/H16 J1 sequences from *H. fugitivus* and *D. semiclausum* are scattered in the phylogeny, falling closer to lepidopteran H12/H16 J1 fragments than to H12 and H16 J1 sequences of *H. didymator* (Supplementary Figure 2). This result suggests that lepidopteran species have suffered IV integrations from multiple, relatively distantly related donor wasps and/or that integrations from a given wasp lineage occurred at different times throughout the evolutionary history of the species.

## Discussion

### Preferential integration of HdIV in hemocytes of *Spodoptera frugiperda*

In this study, we have shown that seven out of the 57 IV circles of the Ichneumonidae wasp *H. didymator* undergo chromosomal integration into two tissues (hemocytes and fat body) of parasitized fall armyworms (*S. frugiperda)*. Three of the circles integrate massively into these somatic cells and we estimate that on average, each host haploid genome suffers between 13 and 40 IEs 72h p.p. These numbers are similar to those reported earlier for *Cotesia typhae* bracovirus (CtBV) which typically undergoes 12 to 85 IEs per host haploid genome on average, depending on the tissue (Muller *et al*. 2021). As observed for CtBV, HdIV integrations are more numerous in the hemolymph than in the fat body. This is also in line with earlier observations made for *Microplitis demolitor* BV (Beck *et al*. 2007). Given that hemocytes are individual cells that circulate through the entire body of caterpillars, they may be more accessible to PDVs than cells of other tissues, such as the fat body, that are akin to cell aggregates in which only external layers may be immediately reachable by PDVs. Alternatively, or in addition, PDVs may have evolved to preferentially target hemocytes, the main mediators of insect immune response (Lavine and Strand 2002; Jiang *et al*. 2010).

### What role for PDV circle integration in parasitism success?

A question emerging from earlier studies of PDVs relates to the role of circle integration during parasitism (Beck *et al*. 2011; Benoist *et al*. 2017; Chevignon *et al*. 2018; Muller *et al*. 2021). In our study of CtBV, we found that similar quantities of integrated circles (*i*.*e*. circles possessing a HIM motif) and nonintegrated circles (no HIM motif) can persist over at least 7 days after oviposition, that is about half the time needed for *C. typhae* larvae to emerge from their host (Muller *et al*. 2021). Thus, it seemed that integration did not play so much of a role in ensuring persistence of BV circles in the context of this earlier study. However, host tissues parasitized by *C. typhae* were sequenced at only one time point in Muller *et al*. (2021), such that we could not assess how the overall quantity of integrated versus non-integrated circles evolves during parasitism. Here, we show that the number of IEs increases with time after oviposition. While we did not perform replicates to test whether this increase is statistically supported, we observe that it holds both for the hemolymph and fat body, which may be considered pseudo-replicates. We deduce that this increase in IEs through time is due to continuous integration of circles from 24h p. p. to 72h p. p., rather than to caterpillar cell replication, since we did not detect more IEs covered by multiple reads (as a result of cell division) at 72h p. p. than at 24 h p. p. Mirroring this increase in integrated circles, we found that the sequencing depth over HdIV, and thus the overall quantity of IV circles, decreases through time (Figure 2). The decrease is particularly striking in the fat body, whereby HdIV sequencing depth is divided by three between 24h and 72h p. p. (Figure 2). The combined observation of an increase in integrated IV circles and a decrease in overall circle quantity implies that the quantity of non-integrated circles decreases quite sharply through time after oviposition, and faster than that of integrated circles. In this context, integration may be seen as a way for the PDVs to persist in larger amounts in host cells throughout the entire duration of wasp progeny development.

### Remarkable similarity in ichnovirus and bracovirus chromosomal integration patterns

This study is the second one investigating integration of IV DNA circles using bulk Illumina sequencing of parasitized host tissues. Like in Wang *et al*. (2021), we show that the majority of IEs occur through double strand breaks located in the HIM. We recently investigated chromosomal integration of a bracovirus (CtBV) during parasitism using the same approach as here and found that CtBV circles also almost exclusively integrated through double strand breaks occurring within the HIM (Muller *et al*. 2021). This confirms the central role of these motifs in mediating circle integration in host somatic cells and ensuring parasitism success. Applying the same method to characterize circle integration of an IV and a BV (Muller *et al*. 2021) allows us to directly compare the two systems. Patterns of chromosomal integrations turn out to be highly similar between HdIV and CtBV, in terms of nature - all IEs are mediated by a HIM motif having a similar structure with conserved inverted repeats separated by a short stretch of sequence not conserved -, tissue tropism (hemocytes as preferential target), and numbers of IEs per host cell. This similarity suggests the mechanisms involved in integration could be highly related, which is unexpected given that IV and BV have an independent origin, deriving from domestication of viruses belonging to different viral families (Gimenez *et al*. 2020; Gauthier *et al*. 2021; Gilbert and Belliardo 2022).

The mechanisms underlying PDV circle integration are not fully understood. In the case of BV, three candidate wasp genes of nudiviral origin coding for Tyrosin recombinases (*vlf-1* and *int-2 homologues*) have been proposed to operate together ensuring the integration of HIM-containing circles (Chevignon *et al*. 2018) and were shown by functional assays to be involved in the integration of some circles of Cotesia vestalis bracovirus (Wang *et al*. 2021). Moreover, there is evidence showing that some BV circles also rely on different lepidopteran retroviral integrases for chromosomal integration (Wang *et al*. 2021). In the case of IV, circle integration has not been studied at the functional level yet, but the viral machinery does not include a gene encoding a protein with a known integrase domain (Legeai *et al*. 2020), which suggests that host integrases could be involved. Indeed, site-specific recombinases/integrases such as retroviral integrases and transposases are abundant in eukaryote genomes. Another possibility is that among the genes of unknown function deriving from the IV ancestor, there is a recombinase from a still uncharacterized family that has not yet been recognized as such. It is noteworthy that the proteins encoded by the conserved gene set of ichnoviruses do not contain any recognizable conserved domain (Volkoff *et al*. 2010). The striking similarities between BV and IV circles integration mechanisms could be explained by the fact that they were shaped by similar structural constraints, despite their independent origin. In fact, the J1 and J2 motifs are much alike inverted terminal repeats of DNA transposons (ITRs). However, unlike ITRs which lie at the extremities of DNA transposon copies, these inverted repeats are located next to each other, only separated by the short stretch of nucleotides deleted during integration. Once a transposase binds to the ITRs, the formation of the transpososome relies on properties of the DNA separating the ITRs, including a minimum length (Hickman *et al*. 2018) (Supplementary Figure 3). Similarly, to form an integration complex starting from a PDV circle, a minimum sequence length between the ITRs is most probably required, which would explain the presence of the intervening DNA between the J1 and J2 motifs (Supplementary Figure 3). The fact that the IV and BV HIM only show very low similarity in primary sequence may be due to the involvement of recombinases belonging to distantly related families. Thus, the action of similar constraints could be sufficient to explain the similarity of the HIM structure of BV and IVs. Confirmation of this hypothesis requires to better characterize the molecular mechanisms involved in BV and IV circles integration. Another possibility would be that recombinases of the same family, which remain to be identified, may act as major players in circle integration of both BVs and IVs. In this case, horizontal transfer events of HIM sites might be invoked to explain that the two PDVs genera share the same mechanism despite their different origin. The similarity in J1 and J2 motifs as well as in other positions of the HIM between BVs and IVs may be viewed as supporting this scenario (Figure 4). Horizontal transfer events have been proposed as an explanation of the common phylogenetic signature shared by the virulence factors V-ANKs of BV and IVs (Falabella *et al*. 2007; Ceirqueira de Araujo *et al*., in press).

Finally, though as for HdIV, Wang *et al*. (2021) found a majority of DsIV IEs occurring at HIM in *Plutella xylostella* hemocytes, they also identified large numbers of IEs involving random positions along both HIM-bearing circles and circles devoid of HIM. This led them to conclude that two distinct mechanisms were involved in DsIV chromosomal integration. Here the number of IE falling outside HIM or in circle devoid of HIM is so low that the biological significance of these integrations in terms of parasitism success in the *H. didymator*/*S. frugiperda* system is questionable. Interestingly, whatever the system (CtBV, DsIV, HdIV), there is an enrichment for microhomologies between the host genome and IV circles at the virus-host junction site, suggesting that host DNA repair mechanisms may be involved in at least a subset of IEs (Muller *et al*. 2021; Wang *et al*. 2021). In fact, the fraction of junctions involving microhomology is higher for IV than for BV as contrary to BV, for IVs we observe almost no junction involving blunt-ended wasp and host sequences or 1-bp microhomology between them.

### Widespread germline infiltrations of polydnaviruses in multiple lepidopteran families

Earlier studies uncovered a number of BV circle fragments in the genome of various species of lepidopterans and proposed that these sequences were horizontally transferred from wasp through HIM-mediated integration in the germline genome of lepidopterans (Schneider and Thomas 2014; Gasmi *et al*. 2015; Di Lelio *et al*. 2019). However, none of the horizontally transferred BV sequences reported in these studies contained the J1 or J2 motif of the HIM. This could be explained either because the transfers were ancient and the HIM was degraded faster than the rest of the circles, or because these transfers occurred through a mechanism not involving a double strand break within the HIM. The finding that CcBV (Chevignon *et al*. 2018), DsIV (Ze-hua Wang *et al*. 2021), CvBV (Zehua Wang *et al*. 2021), CtBV (Muller *et al*. 2021) and HdIV (this study) undergo massive HIM-mediated chromosomal integration in host somatic cells during parasitism, together with the recent availability of hundreds of lepidopteran whole genome sequences led us to assess the extent to which this mechanism fostered wasp-to-lepidopteran HT during evolution. We uncovered dozens of J1- or J2-bordered IV and BV fragments in a total of 124 lepidopteran species belonging to 15 families. In addition to show that sequencing depth was homogenous over lepidopteran/PDV circle junctions for most of these fragments, we also found that they were supported by multiple chimeric reads, contrasting with somatic PDV circle insertions that are almost all covered by only one chimeric read. Furthermore, we PCR-validated typical extremities of HIM mediated junctions of two IV and two BV integrations in multiple individuals of two species of *Pieris* butterflies and uncovered multiple integrations shared at orthologous loci between two to four lepidopteran species. Altogether these results indicate that HIM-mediated germline infiltration of PDV circles is widespread in lepidopterans, the main hosts of braconid and ichneumonid parasitoid wasps.

### Limitations of the current study and perspectives

The number of J1- or J2-bordered PDV circle fragments we report here is necessarily underestimated because we only used HdIV and CtBV circles, for which HIM were annotated using the same approach (Muller *et al*. 2021; This study), as queries to perform our similarity searches. In addition, we used relatively conservative criteria to retain blast hits for downstream analysis (at least 300-bp long and e-value < 0.0001) and many genomes included in our search are of poor quality (only 210 out of 775 genomes included have a N50 > 10 Kb). However, our phylogenetic analyses reveal widespread polyphyly of PDV fragments at the lepidopteran family, genus and even species level, indicating that a given lepidopteran species or multiple species within a family or a genus have received PDV fragments from various wasp donor species. We found that the number of PDV fragments uncovered per lepidopteran family was positively correlated with the number of genomes available in each family, with no apparent biological factor further explaining the distribution of these fragments in Lepidoptera. It is however noteworthy that some species have remarkably high numbers of PDV fragments (e.g., 81 and 80 PDV fragments in 2 *Blastobasis* species and 1756 PDV fragments in *Y. scabrella*) that were acquired from multiple donor wasp species (Figure 12 and Supplementary figure 2). We are unaware of how frequently these species are targeted by parasitoid wasps in the wild. By contrast, all lepidopteran species that are known hosts of CtBV (host: *Sesamia nonagrioides*), CcBV (host: *Manduca sexta*), CvBV (host: *Plutella xylostella*), HdIV (host: *Helicoverpa armigera*) and DsIV (host: *Plutella xylostella*) are devoid of HIM-mediated PDV insertions. It will be interesting to assess what factors may foster HIM-mediated germline integration of PDV circles in different hosts or non-hosts lepidopteran species using a dedicated and comprehensive sampling of parasitoid wasp and lepidopteran species.

This study is only a first step in the large-scale characterization of wasp-to-host HT of PDV circles. We chose to only focus on J1- or J2-bordered PDV fragments because these fragments contain the molecular signature typical of well-characterized chromosomal integrations occurring during parasitism. We could thus formulate explicit predictions regarding expected patterns of sequencing depth and number of chimeric reads. However, our similarity search uncovered many more PDV-like sequences not bordered by the J1 or J2 motifs of the HIM in lepidopteran genomes. For example, our initial blastn search yielded 2,214 hits longer than 300 bp with an evalue < 0.0001 to HdIV circles devoid of HIM. This number was even much higher (n = 19,558) when using CtBV devoid of HIM as queries. Though these sequences likely underwent some form of HT involving lepidopterans and parasitoid wasps, another full dedicated study will be necessary to determine and quantify which scenarios best explain these transfers. For example, it is possible that some of these sequences originate from donor wasp species in which these PDV circles bear a HIM motif but that this motif is not present in the homologous PDV circles of the wasps we used as queries to perform our searches (*H. didymator* or *C. typhae*). Alternatively, the fact that nucleocapsids of PDVs are able to enter the nuclei may also favour integration of circles by a more general mechanism such as DNA repair. Although we did not observe a significant level of circle integration not involving HIM sites during parasitism, such rare events could occur in the germline at the time scale of evolution. It is also possible that many of these sequences correspond to transposable elements present in HdIV and CtBV (Dupuy *et al*. 2011) and that these TEs were horizontally transferred either together with PDV circles or independently, as proposed by (Heringer and Kuhn 2022). Another limitation of this study is that it is focused only on lepidopterans, the major hosts of PDV-encoding braconid and campoplegine parasitoid wasps (Gauld 1988). Yet, some wasps within the two families are known to parasitize non-lepidopteran hosts (Robin *et al*. 2019) and it will be interesting to extend the search for HIM-mediated HT of PDV circles to a larger diversity of hosts. Extending the search to species not known to be hosts of parasitoid wasps may also reveal unexpected wasp-host interactions.

### A major route of horizontal transfer of genetic material among insects

Horizontal transfer of genetic material is widespread and a major force shaping prokaryote evolution (Soucy *et al*. 2015). In eukaryotes, the importance of HT is increasingly recognized (Husnik and McCutcheon 2018; Sibbald *et al*. 2020; Van Etten and Bhattacharya 2020), but the extent to which it influenced genome evolution remains debated, especially in animals and other multicellular taxa (Martin 2017; Salzberg 2017). Several studies reported spectacular individual cases of HT of genes in various animals, often with functional evidence supporting an important role played by horizontally transferred genes in the recipient species (Moran and Jarvik 2010; Acuna *et al*. 2012; Wybouw *et al*. 2014; Danchin *et al*. 2016; Leclercq *et al*. 2016; Wybouw *et al*. 2016; Gasmi *et al*. 2021; Xia *et al*. 2021; Cummings *et al*. 2022). Large-scale analyses of HT of genes and TEs in insects tend to show that these transfers occurred recurrently in most insect lineages and likely had important consequences on insect genome evolution (Peccoud *et al*. 2017; Li *et al*. 2022). So far however, the mechanisms through which these transfers occurred remain unclear and no mechanism dedicated to HT is known in animals. Together with previous studies on CcBV (Chevignon *et al*. 2018), DsIV (Ze-hua Wang *et al*. 2021), CvBV (Zehua Wang *et al*. 2021) and CtBV (Muller *et al*. 2021), our work on HdIV shows that parasitoid wasps use massive HIM-mediated HT of PDV circles to hijack host somatic cells, ensuring parasitism success. We also show that though not dedicated to germline integration, this mechanism fostered many accidental HT of PDV sequences in the germline of a large number of lepidopteran hosts that survived to parasitism (or to the injection of PDV only) and transmitted these sequences to their offspring. HIM-mediated HT of PDV sequences may thus be viewed as a major route of HT in insects. The extent to which it may have facilitated the horizontal transfer of non-PDV sequences integrated into PDV DNA circles or co-packaged with PDV circles into PDV viral particles remains to be assessed. Two studies have provided functional evidence that a PDV gene (*Sl* gasmin), acquired by a noctuid moth through HT, plays an essential role in the antibacterial immune response of the moth (Gasmi *et al*. 2015; Di Lelio *et al*. 2019). Our work suggests that many PDV genes have been acquired by lepidopterans through HIM-mediated HT. It opens new avenues to further quantify this phenomenon and the impact it had on insect evolution.

## Material and Methods

### Insects rearing, parasitization and DNA sequencing

The *Spodoptera frugiperda* laboratory colony is maintained on a semi-synthetic maize diet under stable conditions (24±2°C; 75–65% relative humidity; 16 h light: 8 h dark photoperiod). The wasp *Hyposoter didymator* laboratory colony is reared on *S. frugiperda* at 26±2°C with a 16 h light: 8 h dark photoperiod. Fourth instar *S. frugiperda* larvae were each parasitized by exposing them to one *H. didymator* female wasp until one oviposition event was observed. Two tissues were sampled: the hemolymph, which is the most targeted by BVs (Muller *et al*. 2021), and the fat body, because it is easy to isolate large quantity of this tissue, in turn facilitating DNA extractions. Hemolymph and fat body of 20 parasitized caterpillars were collected after 24h or 72h after oviposition (*i*.*e*., post-parasitism (pp)). Hemolymph was collected from the caterpillar proleg and stocked into 1.5 mL of ATL buffer from the Qiagen DNeasy Blood & Tissue Kit. The fat body was collected, washed with PBS buffer and also stored into 1.5 mL of ATL buffer after caterpillars were dissected and the digestive tract removed. The genomic DNA from the four caterpillar pools was extracted using the DNeasy Blood & Tissue Kit. DNA of the 4 samples from the parasitized caterpillars *S. frugiperda* was quantified on a Qubit apparatus and was sent to Novogene for Illumina sequencing in a 2×150bp paired-end mode.

### Reference genomes used to characterize somatic insertions of Hyposoter didymator Ichnovirus

The genome of *S. frugiperda* (corn strain) was sequenced by (Gimenez *et al*. 2020) using the long-read, PacBio RSII (Pacific Biosciences) technology and assembled with Platanus. It is available on the BIPAA platform (https://bipaa.genouest.org/is/; Rennes, France) and under accession number PRJNA662887 in NCBI. This genome is 384.46Mb, it has an N50 (*i*.*e*., minimum contig length that covers 50 percent of the genome) of 13.15Mb and it is composed of 125 contigs.

The genome of *H. didymator* was sequenced by Legeai *et al*. (2020) using the short-read Illumina HiSeq technology and assembled with Platanus. It is available in the NCBI database under accession number PRJNA589497. The genome is 226,9Mb, it has an N50 of 3,3 Mb and it is composed of 131,161 sequences. HdIV regions were annotated by Legeai *et al*. (2020) using the genome annotation editor Apollo browser. 57 viral segments localized in 66 viral loci were annotated in 32 scaffolds.

### Measuring sequencing depth on *Hyposoter didymator* and *Spodoptera frugiperda* genomes

The four datasets of Illumina reads we obtained (two parasitized caterpillar tissues at two timepoints post-parasitism; SRA accession numbers provided in Table 1) were quality-trimmed with Trimmomatic v.0.38 (LEADING:20 TRAILING:20 SLIDINGWINDOW:4:15 MINLEN:36 options) (Bolger *et al*. 2014) and the quality was evaluated with FastQC v.0.11.8 (Wingett and Andrews 2018). To get statistics on sequencing depth, we aligned the four datasets of trimmed paired-end reads using Bowtie2 v.2.3.4.1 (Langmead and Salzberg 2012) in end-to-end mode on both *S. frugiperda* and *H. didymator* reference genomes. We converted the output SAM files into sorted BAM files with samtools v.1.9. The sequencing depth for each of our samples was estimated with bedtools genomecov v2.26.0 (Quinlan and Hall 2010) on both reference genomes (-ibam and -d options). To get the sequencing depth of *H. didymator* proviral segments, we used bedtools coverage v2.26.0 (-d option) and provided as input a bed file containing the proviral segment coordinates and the bam file of the Illumina reads aligned on the *H. didymator* genome.

### Characterizing integrations of wasp ichnovirus DNA circles in somatic host genomes

To identify integrations of HdIV DNA circles in the *S. frugiperda* somatic genomes, we searched for chimeric reads, *i*.*e*., reads for which a portion aligns exclusively on HdIV and another portion aligns exclusively on the *S. frugiperda* genome. Reverse and forward raw fastq files were converted into fasta files. The resulting fasta files were aligned on the wasp genome with blastn version 2.6.0 (-task megablast -max_target_seqs 1 -outfmt 6). Reads aligning on the wasp genome were extracted from the fasta file using seqtk subseq (https://github.com/lh3/seqtk) and aligned with blastn (same options as above) on the *S. frugiperda* genome.

To further find chimeric reads most likely to result from the integration of HdIV into *S. frugiperda* genome, we used the approach described in Muller *et al*. (2021). Briefly, we used the tabulated outputs of the two successive blastn searches as entries of an R pipeline initially written to identify artificial chimeras generated during deep sequencing library preparation (Peccoud *et al*. 2018). This script identifies reads as chimeric reads if (i) at least 90 % of the read aligns on one or the other genome, (ii) at least 16 bp align *only* on one of the two genomes, (iii) with a maximal of 20 bp overlap between the two read portions aligning on a different genome and (iv) a maximal of random 5 bp insert at the chimeric junction. Then, we estimated the number of independent integration events (IEs) by counting as one event all reads with identical or nearly identical coordinates in the two genomes. To account for possible sequencing errors and/or alignment differences between reads covering the same HdIV – *S. frugiperda* junction (resulting from the fact that the length of the chimeric read region corresponding to each genome varies between reads), we allowed the coordinate of chimeric reads to vary by 5 - bp in the two genomes, *i*.*e*., all reads aligning at the same position +/- 5 bp in the two genomes were considered as reflecting the same integration events.

To be able to compare numbers of integration events between samples, we normalized values by the number of sequenced reads mapping to the *S. frugiperda* genome, as in Muller *et al*. (2021). We calculated a scaling factor by dividing the number of reads mapped on the *S. frugiperda* genome by 1,000,000. Then, to obtain the number of IEs per million reads mapping on *S. frugiperda* for each HdIV circle, we devided the absolute number of IEs by this scaling factor. We used samtools view (version 1.9, option -c -F 4) to determine the number of reads mapped on the *S. frugiperda* genome.

### Searching for polydnavirus DNA circles in lepidopteran genomes

To search for HIM-mediated horizontal transfer of PDV circles in lepidopteran genomes available in Genbank, we used the seven HIM-containing HdIV circles and the 16 HIM-containing Cotesia typhae BV (CtBV) circles previously described in Muller *et al*. (2021) as queries to perform similarity searches using blastn (-task blastn) on lepidopteran genomes. All HdIV and CtBV DNA circle sequences are provided in Supplementary Dataset 6 and the HIM coordinates within these circles are provided in Supplementary Dataset 7. The sequences of each HIM motifs of HdIV and CtBV circles are also provided as a multiple alignment in Supplementary Dataset 8 as well as in Figure 4. A total of 844 lepidopteran genomes were available in Genbank as of December 2021. When more than one genome per species was available, we only retained the largest one, resulting in 775 genomes that were submitted to our search. Accession numbers, size and N50 of these genomes are provided in Supplementary Table 6. Blastn outputs were filtered in R (R Core Team) based on hit size, e-value and coordinates. Some PDV circles are similar to each other. For example, Hd12 and Hd16 are similar to each other over most of their length. To avoid counting multiple times the same lepidopteran genome region as resulting from HIM-mediated horizontal transfer, alignment coordinates of blastn hits were thus merged using bedtools v2.26.0 merge (-s -c 4 -o distinct) (Quinlan and Hall 2010). Merging was performed independently for IV-like and BV-like sequences. The script written for this part of the study is available on GitHub: https://github.com/HeloiseMuller/Chimera/tree/master/scriptsArticles/Heisserer2022.

### Sequencing depth surrounding junctions between PDV DNA circles and lepidopteran genomes

To compute sequencing depths on junctions between PDV DNA circles and lepidopteran genomes, raw Illumina reads were downloaded from Genbank for nine species in which more than four junctions were found. Accession numbers of these reads are provided in Supplementary Table 7. Reads were mapped on the genome of the nine species using bowtie2 (default options) (Langmead and Salzberg 2012). Read depth over 300 bp upstream and downstream of each junction was obtained using bedtools v2.26.0 coverage (Quinlan and Hall 2010).

### Search for orthologous PDV fragments in lepidopteran genomes

We extracted all PDV fragments ending in the HIM together with 1000 bp flanking the J1 or J2 motif of the HIM. We filtered out fragments showing similarity to PDV sequences over > 100 bp in their flank, which may result from integration of PDV circles next to (or within) each other. We then used blastn to align all PDV fragments plus flanks on themselves. Self blastn hits as well as hits involving sequences corresponding to different circles were filtered out. We retained blastn hits involving the PDV fragments plus at least 200 bp of flanking regions in both sequences as candidate orthologous sequences. These candidate orthologous sequences were then all submitted to manual inspection to only retain PDV sequences integrated at the very same position in two species.

### PCR verifications

To experimentally verified that PDV are present in natural populations of butterflies, twelve individuals of *Pieris napi* and twelve individuals of *P. rapae* were collected in 2018 in three sites in France: S1, Pénerf (Morbihan) 47.511463, -2,622753; S2, Vernou-sur Brenne (Indre-et-Loire) 47.415418 0.858978; S3, Tours (Indre-et-Loire) 47.355069 0,702746. PDV junctions were identified in the genome of *P. napi* and *P. rapae* using similarity searches (blastn). This study was performed in 2018, before the large-scale bioinformatic study reported in this article. It was performed on two different assemblies of the *P. napi* genome (CAJQFU010000000 and DWAF00000000) while the large-scale bioinformatic study only included one (DWAF00000000). While ichnovirus circles used as queries for the blastn searches were those of *Hypososter didymator* (the same as those used in the large-scale bioinformatic study), bracovirus circles were those of Cotesia congregata bracovirus (instead of CtBV in the large-scale bioinformatic study). This explains why only one junction (out of six) verified by PCR was also found in the large-scale bioinformatic search. PCR primers were designed on each side of each PDV-butterfly junction. DNA was extracted from all 24 butterfly and a standard PCR protocol was used to screen for the presence/absence of six junctions in these individuals. Junctions found in at least two individuals were Sanger-sequenced and are provided in Supplementary Dataset 2, together with the genomic coordinates of each junction in the genomes of *P. napi* and *P. rapae*.

### Phylogenetic analyses

Sequences homologous to PDV DNA circles and starting or ending within HIM were extracted from lepidopteran genomes using seqtk subseq (https://github.com/lh3/seqtk) and are provided in Supplementary Table 1 and 2. These sequences were then aligned, together with homologous DNA circles from various parasitoid wasps (*D. semiclausum, H. didymator* and *H. fugitivus* for IV and *Cotesia sesamiae, C. vestalis* and *C. congregata* for BV) using MUSCLE (Edgar 2004). Given the large number of IV junctions found in some species, we clustered these sequences for all species in which more than five sequences were found for a given junction using Trimal (-maxidentity 0.95) (Capella-Gutierrez *et al*. 2009). Alignments were then trimmed with Trimal (-automated1) and submitted to Neighbor-joining analysis in MEGA 11 (Maximum Composite Likelihood model of substitution, uniform rates among sites and lineages, pairwise deletion) (Tamura *et al*. 2021).

## List of supplementary material

**Supplementary Figure 1**. Map of chimeric reads along all Hyposoter didymator ichnovirus (HdIV) circles containing a HIM. The vast majority of chimeric reads map to the HIM region (white), indicating HIM-mediated chromosomal integration. On the left are maps of full length HdIV proviral segments. The maps on the right correspond to a zoom on the HIM region of each proviral segment.

**Supplementary Figure 2**. Unrooted phylogenies of HIM-bordered PDV fragments found in lepidopteran genomes and their homologous PDV circles from various wasp species (in black). A) IV fragments homologous to the J1 junction of the Hyposoter didymator IV 12 and/or Hyposoter didymator IV16 circle. B) BV fragments homologous to the J2 junction of the Cotesia typhae BV 2 circle. In A), prior to alignment, the numerous fragments found in *Ypsolopha scabrella, Blastobasis adustella* and *Blastobasis lacticolella* were clusterized at 95% nucleotide identity threshold. Species names are colored according to lepidopteran family and the number associated to each species name is a unique identifier that allows retrieving the sequence in Supplementary Table 1 (column “Number”). Numbers on branches are bootstrap values higher than 70%. The trees were built using the Maximum Composite Likelihood model of substitution available in MEGA 11 (Tamura *et al*. 2021) with 100 bootstrap replicates.

**Supplementary Figure 3**. Hypothesis that thermodynamic constraints might explain the similarity of the HIM-mediated mechanism of ichnovirus and bracovirus integration. A) Transposon excision from the genome is the first step of transposition and involves the binding of recombinases monomers to their specific binding sites at the extremities of the transposable element that come into contact following the dimerisation of the recombinase protein. The flexibility of the genomic DNA is required to form the synaptic complex. B) in the case of polydnavirus circles integration, flexibilty is also required to form the synaptic complex which is probably conferred by the loop, which sequence is lost following integration. This results in an integrated copy of the circle in parasitized host DNA which has lost the loop sequence. For a transposon, the loss of a sequence would result in the inability to transpose again while for ichnovirus and bracovirus circles this does not prevent the expression of genes involved in parasitism success. Moreover, the integrated copies are not transmitted since parasitized Lepidoptera do not reach the adult stage, only the original copy present in the wasp genome is vertically transmitted to the wasp progeny. It is noteworthy that given the HIM structure; the length of the loop is minimal resulting only in the loss of a short sequence. The structural similarity between and ichnovirus and bracovirus HIMs could stem from similar constraints to transfer DNA from one organism to another resulting in convergent evolution. Another possibility is that HIMs were horizontally transferred between ichnoviruses and bracoviruses, which could explain the low level of similarity observed in primary sequence between the two PDV types (Figure 4).

**Supplementary Dataset 1**. Examples of full-length IV circles integrated into lepidopteran genomes. This file contains 7 alignments of full-length PDV circles found in lepidopteran genomes. For each alignment the PDV circle integrated into the lepidopteran genome is aligned onto a circle from HdIV or CtBV which has been linearized at the J1 and J2 motifs. HdIV and CtBV circles are linearized within the HIM and bordered by the J1 and J2 motif. J1 and J2 motifs of each HdIV and CtBV circle are shown in Figure 4. Each alignment is separated by a blank sequence. The HdIV16-like full length circle found in the genome of *Ypsolopha scabrella* (CAJUZE010000643.1:290500-30298) contains a nearly full length LTR retrotransposon.

**Supplementary Dataset 2. Sequences and multiple alignments of the six polydnavirus-butterfly junctions that were PCR verified and Sanger-sequenced in populations of *Pieris napi* and *Pieris rapae***. For each junction, the reference sequence initially identified in butterfly genomes is provided, together with accession number and coordinates. The polydnavirus region is highlighted in yellow and the butterfly region is highlighted in blue. Below we two to four sequences of junctions obtained using Sanger sequencing in our lab in various butterfly individuals, as well as a multiple alignment of these sequences and the reference sequence.

**Supplementary Dataset 3**. Examples of J1- or J2-bordered IV fragments shared at orthologous locus between at least two lepidopteran species. This file contains 12 alignments, each made of four sequences. The two top sequences are the orthologous J1- or J2-bordered IV fragments, together with their flanking region, found in two lepidopteran species. The third sequence is the longest IV fragment from these two species (sometimes the fragments are the same size in the two species, in which case we randomly selected one of them), provided to facilitate localization of the IV-host genome boundaries. The fourth sequence is the homologous HdIV circle. Each alignment of orthologous locus is separated by a blank sequence. J1 and J2 motifs of each HdIV circle are shown in Figure 4. Numbers and letters provided at the beginning of the name of each sequence correspond to those in Table 2.

**Supplementary Dataset 4**. Examples of J1- or J2-bordered BV fragments shared at orthologous locus between at least two lepidopteran species. This file contains 5 alignments, each made of four sequences. The two top sequences are the orthologous J1- or J2-bordered BV fragments, together with their flanking region, found in two lepidopteran species. The third sequence is the longest BV fragment from these two species (sometimes the fragments are the same size in the two species, in which case we randomly selected one of them), provided to facilitate localization of the BV-host genome boundaries. The fourth sequence is the homologous CtBV circle. Each alignment of orthologous locus is separated by a blank sequence. J1 and J2 motifs of each HdIV circle are shown in Figure 4. Numbers and letters provided at the beginning of the name of each sequence correspond to those in Table 2.

**Supplementary Dataset 5**. Examples of paralogous J1- or J2-bordered IV fragments in lepidopteran genomes. This file contains 54 alignments, each made of four sequences. The two top sequences are the paralogous J1- or J2-bordered IV fragments, together with their flanking region, found in the same lepidopteran species. The third sequence is the longest IV fragment from these two loci (sometimes the fragments are the same size in the two loci, in which case we randomly selected one of them), provided to facilitate localization of the IV-host genome boundaries. The fourth sequence is the homologous HdIV circle. Each alignment of paralogous locus is separated by a blank sequence. Numbers and letters provided at the beginning of the name of each sequence correspond to those in Supplementary Table 5.

**Supplementary Dataset 6**. Sequence of all HdIV and CtBV DNA circles in fasta format.

**Supplementary Dataset 7**. Coordinates of HIM in HIM-containing HdIV and CtBV DNA circles (Supplementary Dataset 6).

**Supplementary Dataset 8**. Multiple alignments of the sequences of each HIM motifs of HdIV and CtBV.

**Supplementary Table 1**. Characteristics of blastn hits retrieved using HIM-containing HdIV circles as queries on 775 lepidopteran genomes. Only hits longer than 300 bp with an e-value < 0.0001 were retained. J1 or J2-bordered PDV fragments found in lepidopteran genomes are provided in the last column.

**Supplementary Table 2:** Characteristics of blastn hits retrieved using HIM-containing CtBV circles as queries on 775 lepidopteran genomes. Only hits longer than 300 bp with an e-value < 0.0001 were retained. J1 or J2-bordered PDV fragments found in lepidopteran genomes are provided in the last column.

**Supplementary Table 3**. Numbers of J1- or J2-bordered IV fragments found in lepidopteran species and characteristics of lepidopteran genomes in which they were found

**Supplementary Table 4**. Numbers of J1- or J2-bordered BV fragments found in lepidopteran species and characteristics of lepidopteran genomes in which they were found.

**Supplementary Table 5**. Characteristics of J1- or J2-bordered IV fragments shared at paralogous loci within lepidopteran species. *Paralogous J1- or J2-bordered IV fragments were identified as pairs of sequences, number refer to their number label in sequence alignments provided in Supplementary Dataset 5. Group letters indicate pairs of sequences belonging to the same paralogous group (e.g., IV fragments 44, 72 and 73 all have a flanking region in common in *Y. scabrella* and are thus likely genomic duplicates of a single original germline IV integration event). Ali. size indicate the length of the IV + flank alignment provided in Supplementary Dataset 5. IV size indicate the size of the largest IV fragment among a group of orthologous fragments.

**Supplementary Table 6**. Accession number and characteristics of the 775 genomes screened for the presence of J1- or J2-bordered PDV fragments in this study.

**Supplementary Table 7**. Accession numbers of illumina reads used to compute average sequencing depth and number of chimeric reads over PDV-host junctions (Figures 9 and 10).

## Acknowledgements

This work was supported by Agence Nationale de la Recherche, project ANR-18-CE02-0021-01TranspHorizon.

## Data availability

All data produced during this study are available in public repositories. Genbank accession numbers of raw reads produced from *Spodoptera frugiperda* larvae parasitized by *Hyposoter didymator* are provided in Table 1. Scripts are provided in GitHub: https://github.com/HeloiseMuller/Chimera/tree/master/scriptsArticles/Heisserer2022. All wasp PDV circles and all lepidopteran PDV fragments bordered by HIM are provided in Supplementary Datasets 6 and Supplementary Tables 1 and 2. Accession numbers of lepidopteran genomes surveyed in this study are provided in Supplementary Table 7.

