## Supplementary Dataset 2 for "Somatic chromosomal integration of polydnavirus during parasitism triggered their germline infiltration in multiple lepidopteran families"

**Supplementary Dataset 2. Sequences and multiple alignments of the six polydnavirus-butterfly junctions that were PCR verified and Sanger-sequenced in populations of *Pieris napi* and *Pieris rapae*.** For each junction, the reference sequence initially identified in butterfly genomes is provided, together with accession number and coordinates. The polydnavirus region is highlighted in yellow and the butterfly region is highlighted in blue. Below we two to four sequences of junctions obtained using Sanger sequencing in our lab in various butterfly individuals, as well as a multiple alignement of these sequences and the reference sequence.

Polydnavirus

Butterfly

>P. napi CcBV1 jonction1 – NCBI accession number (start-end coordinates): DWAF01107089.1 (77-796)

GTATAGCAAACCGTAGAGATCATCAACCCGTTACGTAAACGAATGTATAGTCAAGGTCATGGACTCGGGGTGGTTTTTGTTTTGAATTTGGTGTGACGTCACTAACGTTTGATCAAGTTCTCGCTTCCCGTCCTAAAAAAAATATTTCAGAACGGGTTCTAAGTCGATTTTCAGACCGCGTACATCCCTGTCTATAAGGTTCTAGTGTTTTTTTCTTAAAANGACAAAACTTAAAAGGCATGCCTCGTATCACTCTCATCAAATTTAAATTGAAAACTATATTTTGGGATATCAGTATTCATCTGAGAACATTAGCAGATAATTTTTTAAATGGCATACTAGAAAAATATTTTTTTGTGTGAGAACCGTCAAACCATAGACCGAGACACCCAATACTTGCCAAGTATTTCCAGTAATAAACTGCCAAGACAAATACATCTGTATCAGCGCTGATTATGATAATATGAGCAGATTGTATCCTTACTGCGTGAAATGTATGGACGAACATTCTTGAATGAGCTTCTGAATGGTTTGATTGTAATTCAAGTAAGTGCCTCACAAATAGAAGACCCTTGCAACTTGAAACATTAAGTTGGATCATCGAATTCTCCACCAATATACATCTCACAGTCTTCATTGACTTGGGCATATTTTGAAGCATTAGCGCACAGAAATCGATCGAGTTGCAATTTATAAATTTAATTAGGGCTACAGTAGAAA

>P. napi CcBV1 jonction1 sample5

CCGTTACGTAAACGAATGTATAGTCATAGACTCGGGGTGGTTTTTGTTTTGAATTTGGTGTGACGTCCTAAAAAAATATTTCAGAACGGGTCTAAGTCGCTTAATTTTCAGATCGCGTACATCCCTGTCTATAAGGTTCTAGTGTTTTTTTCTTAAAACGACAAAACTTAAAAGGCATGCCTCGTATCACTCTCATCAAATTTAAATTGAAAGCTATATTTTAGGATATCAGTATTAATCTGAGAACATTAGCAGATAATTTTTTAAATGGCATACTAGAAAAATAT

>P. napi CcBV1 jonction1 sample7 lower PCR band

CCGTTACGTAAACGAATGTATAGTCAAGGTCATGGACTCGGGGTGGTTTTTGTTTTGAATTTGGTGTGACGTCACTAACGTTTGCTCAAGTTCTCGCTTCCCGTCCTAAAAAAATATTTCAGAACGGGTTTTAAGTCGCTTAATTTTCAGATCGCGTACATCCCTGTCTATAAGGTTCTAGTGTTTTTTTTTTTAAATGACAAAACTTAAAAGGCATGCCTCGTATTATTATAATTTTTAAATGGCATACTAGAAAAATATTTTTGTGTGAGAACCGTCA

>P. napi CcBV1 jonction1 sample7 upper PCR band

CCGTTACGTAAACGAATGTATAGTCAAGGTCATGGACTCGGGGTGGTTTTTGTTTTGAATTTGGTGTGACGTCACTAACGTTTACTCAAGTTCTCGGTTCCCGTCCTAAAAAAAATATTTCAGAACGGGGTCTAAGTCGCTTAATTTTCAGATCGCGTCCATCCCTGTCTATAAGGTTATAGTGTTTTTTTCTTAAAACGACAAAACTTAAAAGGCATGCCTCGTATCACTCTCATCAAATTTAAATTGAAAGCTATATTTTAGGATATCAGTATTAATCTGAGAACATTAGCAGATAATTTTTTAAATGGCATACTAGAAAAATATTTTTTTGTGGAGAACCG

>P. napi CcBV1 jonction1 sample22

ACGAACAATATCACGTCAAGGACTTGGGGTGCCGTTACGTAAACGAATGTATAGTCATAGACTCGGGGTGGTTTTTGTTTTGAATTTGGTGTGACGTCCTAAAAAAATATTTCAGAACGGGTCTAAGTCGCTTAATTTTCAGATCGCGTACATCCCTGTCTATAAGGTTTTAGTGTTTTTTTCTTAAAACGACAAAACTTAAAAGGCATGCCTCGTATCACTCTCATCAAATTTAAATTGAAAGCTATATTTTAGGATATCAGTATTAATCCGAGAACATTAGCAGATAATTTTTTACAATGGCATACTAGAAAAAT

Multiple alignment

```
                                         10        20        30        40        50        60
                                          |         |         |         |         |         |
PnapiCcBV1jonction1sample5       -------------------------------CCGTTACGTAAACGAATGTATAGTCA---
PnapiCcBV1jonction1sample22      ACGAACAATATCACGTCAAGGACTTGGGGTGCCGTTACGTAAACGAATGTATAGTCA---
PnapiCcBV1jonction1sample7uppe   -------------------------------CCGTTACGTAAACGAATGTATAGTCAAGG
PnapiCcBV1jonction1squenceNCBI   --GTATAGCAAACCGTAGAGATCATCAAC--CCGTTACGTAAACGAATGTATAGTCAAGG
PnapiCcBV1jonction1sample7lowe   -------------------------------CCGTTACGTAAACGAATGTATAGTCAAGG
                                                                **************************
Prim.cons.                       ACG2A2A22A222CGT22AG22C2T2222TGCCGTTACGTAAACGAATGTATAGTCAAGG

                                         70        80        90       100       110       120
                                          |         |         |         |         |         |
PnapiCcBV1jonction1sample5       ---TAGACTCGGGGTGGTTTTTGTTTTGAATTTGGTGTGACGTCCT--------------
PnapiCcBV1jonction1sample22      ---TAGACTCGGGGTGGTTTTTGTTTTGAATTTGGTGTGACGTCCT--------------
PnapiCcBV1jonction1sample7uppe   TCATGGACTCGGGGTGGTTTTTGTTTTGAATTTGGTGTGACGTCACTAACGTTTACTCAA
PnapiCcBV1jonction1squenceNCBI   TCATGGACTCGGGGTGGTTTTTGTTTTGAATTTGGTGTGACGTCACTAACGTTTGATCAA
PnapiCcBV1jonction1sample7lowe   TCATGGACTCGGGGTGGTTTTTGTTTTGAATTTGGTGTGACGTCACTAACGTTTGCTCAA
                                    * ***************************************
Prim.cons.                       TCATGGACTCGGGGTGGTTTTTGTTTTGAATTTGGTGTGACGTCACTAACGTTTGCTCAA

                                        130       140       150       160       170       180
                                          |         |         |         |         |         |
PnapiCcBV1jonction1sample5       -------------------AAAAAAATATTTCAGAACGGG-TCTAAGTCGCTTAATTTTC
PnapiCcBV1jonction1sample22      -------------------AAAAAAATATTTCAGAACGGG-TCTAAGTCGCTTAATTTTC
PnapiCcBV1jonction1sample7uppe   GTTCTCGGTTCCCGTCCTAAAAAAAATATTTCAGAACGGGGTCTAAGTCGCTTAATTTTC
PnapiCcBV1jonction1squenceNCBI   GTTCTCGCTTCCCGTCCTAAAAAAAATATTTCAGAACGGGTTCTAAGTCG----ATTTTC
PnapiCcBV1jonction1sample7lowe   GTTCTCGCTTCCCGTCCT-AAAAAAATATTTCAGAACGGGTTTTAAGTCGCTTAATTTTC
                                                    ********************* * *******    ******
Prim.cons.                       GTTCTCGCTTCCCGTCCTAAAAAAAATATTTCAGAACGGGTTCTAAGTCGCTTAATTTTC

                                        190       200       210       220       230       240
                                          |         |         |         |         |         |
PnapiCcBV1jonction1sample5       AGATCGCGTACATCCCTGTCTATAAGGTTCTAGTGTTTTTTTCTTAAAACGACAAAACTT
PnapiCcBV1jonction1sample22      AGATCGCGTACATCCCTGTCTATAAGGTTTTAGTGTTTTTTTCTTAAAACGACAAAACTT
PnapiCcBV1jonction1sample7uppe   AGATCGCGTCCATCCCTGTCTATAAGGTTATAGTGTTTTTTTCTTAAAACGACAAAACTT
PnapiCcBV1jonction1squenceNCBI   AGACCGCGTACATCCCTGTCTATAAGGTTCTAGTGTTTTTTTCTTAAAANGACAAAACTT
PnapiCcBV1jonction1sample7lowe   AGATCGCGTACATCCCTGTCTATAAGGTTCTAGTGTTTTTTTTTTTAAATGACAAAACTT
                                 *** ***** ******************* ************ ** *** **********
Prim.cons.                       AGATCGCGTACATCCCTGTCTATAAGGTTCTAGTGTTTTTTTCTTAAAACGACAAAACTT

                                        250       260       270       280       290       300
                                          |         |         |         |         |         |
PnapiCcBV1jonction1sample5       AAAAGGCATGCCTCGTATCACTCTCATCAAATTTAAATTGAAAGCTATATTTTAGGATAT
PnapiCcBV1jonction1sample22      AAAAGGCATGCCTCGTATCACTCTCATCAAATTTAAATTGAAAGCTATATTTTAGGATAT
PnapiCcBV1jonction1sample7uppe   AAAAGGCATGCCTCGTATCACTCTCATCAAATTTAAATTGAAAGCTATATTTTAGGATAT
PnapiCcBV1jonction1squenceNCBI   AAAAGGCATGCCTCGTATCACTCTCATCAAATTTAAATTGAAAACTATATTTTGGGATAT
PnapiCcBV1jonction1sample7lowe   AAAAGGCATGCCTCGTATTATTATAATT---TTTAAATGGCATACTAGA----AAAATAT
                                 ****************** * * * **    ******* * *  *** *       ****
Prim.cons.                       AAAAGGCATGCCTCGTATCACTCTCATCAAATTTAAATTGAAAGCTATATTTTAGGATAT

                                        310       320       330       340       350       360
                                          |         |         |         |         |         |
PnapiCcBV1jonction1sample5       CAGTATTAATCTGAGAACATTAGCAGATAATTTTTTA-AATGGCATACTAGAAAAATAT-
PnapiCcBV1jonction1sample22      CAGTATTAATCCGAGAACATTAGCAGATAATTTTTTACAATGGCATACTAGAAAAAT---
PnapiCcBV1jonction1sample7uppe   CAGTATTAATCTGAGAACATTAGCAGATAATTTTTTA-AATGGCATACTAGAAAAATATT
PnapiCcBV1jonction1squenceNCBI   CAGTATTCATCTGAGAACATTAGCAGATAATTTTTTA-AATGGCATACTAGAAAAATATT
PnapiCcBV1jonction1sample7lowe   T----TTTGTGTGAGAACCGTCA-------------------------------------
                                      **  *  ******  *
Prim.cons.                       CAGTATTAATCTGAGAACATTAGCAGATAATTTTTTACAATGGCATACTAGAAAAATATT

                                        370       380       390       400       410       420
                                          |         |         |         |         |         |
PnapiCcBV1jonction1sample5       ------------------------------------------------------------
PnapiCcBV1jonction1sample22      ------------------------------------------------------------
PnapiCcBV1jonction1sample7uppe   TTTTTGTGGAGAACCG--------------------------------------------
PnapiCcBV1jonction1squenceNCBI   TTTTTGTGTGAGAACCGTCAAACCATAGACCGAGACACCCAATACTTGCCAAGTATTTCC
PnapiCcBV1jonction1sample7lowe   ------------------------------------------------------------

Prim.cons.                       TTTTTGTG2222A2C2GTCAAACCATAGACCGAGACACCCAATACTTGCCAAGTATTTCC

                                        430       440       450       460       470       480
                                          |         |         |         |         |         |
PnapiCcBV1jonction1sample5       ------------------------------------------------------------
PnapiCcBV1jonction1sample22      ------------------------------------------------------------
PnapiCcBV1jonction1sample7uppe   ------------------------------------------------------------
PnapiCcBV1jonction1squenceNCBI   AGTAATAAACTGCCAAGACAAATACATCTGTATCAGCGCTGATTATGATAATATGAGCAG
PnapiCcBV1jonction1sample7lowe   ------------------------------------------------------------

Prim.cons.                       AGTAATAAACTGCCAAGACAAATACATCTGTATCAGCGCTGATTATGATAATATGAGCAG

                                        490       500       510       520       530       540
                                          |         |         |         |         |         |
PnapiCcBV1jonction1sample5       ------------------------------------------------------------
PnapiCcBV1jonction1sample22      ------------------------------------------------------------
PnapiCcBV1jonction1sample7uppe   ------------------------------------------------------------
PnapiCcBV1jonction1squenceNCBI   ATTGTATCCTTACTGCGTGAAATGTATGGACGAACATTCTTGAATGAGCTTCTGAATGGT
PnapiCcBV1jonction1sample7lowe   ------------------------------------------------------------

Prim.cons.                       ATTGTATCCTTACTGCGTGAAATGTATGGACGAACATTCTTGAATGAGCTTCTGAATGGT

                                        550       560       570       580       590       600
                                          |         |         |         |         |         |
PnapiCcBV1jonction1sample5       ------------------------------------------------------------
PnapiCcBV1jonction1sample22      ------------------------------------------------------------
PnapiCcBV1jonction1sample7uppe   ------------------------------------------------------------
PnapiCcBV1jonction1squenceNCBI   TTGATTGTAATTCAAGTAAGTGCCTCACAAATAGAAGACCCTTGCAACTTGAAACATTAA
PnapiCcBV1jonction1sample7lowe   ------------------------------------------------------------

Prim.cons.                       TTGATTGTAATTCAAGTAAGTGCCTCACAAATAGAAGACCCTTGCAACTTGAAACATTAA

                                        610       620       630       640       650       660
                                          |         |         |         |         |         |
PnapiCcBV1jonction1sample5       ------------------------------------------------------------
PnapiCcBV1jonction1sample22      ------------------------------------------------------------
PnapiCcBV1jonction1sample7uppe   ------------------------------------------------------------
PnapiCcBV1jonction1squenceNCBI   GTTGGATCATCGAATTCTCCACCAATATACATCTCACAGTCTTCATTGACTTGGGCATAT
PnapiCcBV1jonction1sample7lowe   ------------------------------------------------------------

Prim.cons.                       GTTGGATCATCGAATTCTCCACCAATATACATCTCACAGTCTTCATTGACTTGGGCATAT

                                        670       680       690       700       710       720
                                          |         |         |         |         |         |
PnapiCcBV1jonction1sample5       ------------------------------------------------------------
PnapiCcBV1jonction1sample22      ------------------------------------------------------------
PnapiCcBV1jonction1sample7uppe   ------------------------------------------------------------
PnapiCcBV1jonction1squenceNCBI   TTTGAAGCATTAGCGCACAGAAATCGATCGAGTTGCAATTTATAAATTTAATTAGGGCTA
PnapiCcBV1jonction1sample7lowe   ------------------------------------------------------------

Prim.cons.                       TTTGAAGCATTAGCGCACAGAAATCGATCGAGTTGCAATTTATAAATTTAATTAGGGCTA


PnapiCcBV1jonction1sample5       ---------
PnapiCcBV1jonction1sample22      ---------
PnapiCcBV1jonction1sample7uppe   ---------
PnapiCcBV1jonction1squenceNCBI   CAGTAGAAA
PnapiCcBV1jonction1sample7lowe   ---------

Prim.cons.                       CAGTAGAAA
```

>P. napi CcBV17 jonction1 – NCBI accession number (start-end coordinates): CAJNIW010000003.1 (8549793-8551292)

TAAACATCAACAACGCTTGCCCAATATTCAATAAAAATTTACTATTAGTAATAATAAAAGATTTACTATAAAATATAACACAAAAAAACAAGTAAATAGTGCCATTATTCTTTAATTTTTTACCGTATAAGCTTCAATATAATCAATTGTTTTTTAGCACTGGATGGGCCCGAGAATGCAGTGGTACCAAACGATGCAAAAGTCTTGGTGTCTTCGCAAGTGGTTGCGAATCCAGACGGGAAAGTTCGGGAAAGTCACGAACGAAAAGTCCTTACCAGAAATATTACAGATAGAGTGAGGGAGACAGAAGAGAAAATACATACTGGCGATACTACGCACGAGGTTAGTTTCTTGCACAGTTTGTAATTTATCGCTTAAATTATAAGTATTTTTAAGAAAGATTTCTCATTGAAAAAGTTATAAGTTGTTAGAACACATCCAATAATAAGGTTATCGGAAATTATAAATAAATTGAATGAGAGATCATTTTAAGAGATTGAACAATTTAAAAAATATAATAATATTTAACTCATTAAACATTAAATAATAACAACAGCAATATTAAACGTTAACCCTGAAACGATATGGTAAATTTTGGCCAATTCAAGAAACGTTCAGGCGACACACATAACCCTATAGGTAGTATGTAAGTAAGCCATTTTTAATCTAGACGGATCTAACGTTAATAAGCTAAATGAAATACTGCTGGTCTGTTGGAGCAGAACCATAGTTTATTTATAAAAGTTGCCACATCGACATAATTAAGAATTTAAAAATACGGAATATTATCGTGGCTCTATACATAATTACAAGTATAAGGCCAGTCAGGACTATAATGTGCATTACCATATTCTGAATATAGTAAATCTCTAGAGATATTCAATTTAGTAAGCATGTGGTGCATTTAAATCACACTGCAATAAAGGCGCCAAAGTAGGGAAGTGCGTTTGACTTTCGTATAATATTAGTAATACTTGCTTGTAATATTAACGCGTAATATGAGTAATACTATTTATTTTTATATAAAAGTTACATTAGTACTTGGAAAGGAAATTAATAATTTTCTGTAAACGTTTTAATTACTTGACGCAAAAACTTACGCGCGAGTGGTACTATCAATAGTATAAAAACAACAAATGATTAAATCTTACCATTTTCTATACATATTATTAAATAAAAACTTAGCTTTGAACAGTAATGGTCGCGATCTTGAAATTTTTAGACTTAAAAAATATTCAGCTGTAGAGTCGATGACGTCACGACGGAATTAATGTCACGGTAAGTCCGTGTTCTCTATGGCGTCATAAAAAGTTCAATATAGAGGGCACGCTGAGTATTTAGCTCCAGTTTATGTGTATTCGTTACGTAGGTAAGGTCGTTGACTCCTATGATACGGCCCTGACCTTTATTATGTAGTCCTGATAAAGGTGAATTTACATTGTGAACATGTTGTTTACTTCTGAGGTATAGTTATTGAGTAAGGTACACACGAAGTGTATA

>P. napi CcBV17 jonction1 sample25

CGTAATATGAGTAATACCATTTCTTTTTAAGTTACATTAGTACTTGGAAAGGAAATCAATTATTTTCTGTAAACGTTTTAACTATTTGACGCAAAAACTTCGCGCGAGTGGTAGGATCAATAGTATAAAAACAACAAATGATTAAATCTTACCATTTTCTATACATATTTTTAAATAAAAACTTAGCTTTGAACAGTAATGGTCGCGATCTTGAAATTTTTAGACTTAAAAAATATTCAGCTGTAGAGTCGATGACGTCACGACGGAATTAATGTCACGGTAAGTCCGTGTTCTCTATGGCGTCATAAAAAGTTCAATATAGAGGGCACGCTGAGTATTTAGCTCCAGTTTATGTGTATTCGTTACGTAGGTAAGGTCGTTGACTCCTATGATACGGCCCTGACCTTTA

>P. napi CcBV17 jonction1 sample28

GTAATATGAGTAATACCATTTCTTTTTAAGTTACATTAGTACTTGGAAAGGAAATCAATTATTTTCTGTAAACGTTTTAACTATTTGACGCAAAAACTTCGCGCGAGTGGTAGGATCAATAGTATAAAAACAACAAATGATTAAATCTTACCATTTTCTATACATATTATTAAATAAAAACTTAGCTTTGAACAGTAATGGTCGCGATCTTGAAATTTTTAGACTTAAAAAATATTCAGCTGTAGAGTCGATGACGTCACGACGGAATTAATGTCACGGTAAGTCCGTGTTCTCTATGGCGTCATAAAAAGTTCAATATAGAGGGCACGCTGAGTATTTAGCTCCAGTTTATGTGTATTCGTTACGTAGGTAAGGTCGTTGACTCCTATGATACGGCCCTGACCTTTA

Multiple alignment

```
                                         10        20        30        40        50        60
                                          |         |         |         |         |         |
PnapiCcBV17jonction1sample25     ------------------------------------------------------------
PnapiCcBV17jonction1sample28     ------------------------------------------------------------
PnapiCcBV17jonction1squenceNCB   TAAACATCAACAACGCTTGCCCAATATTCAATAAAAATTTACTATTAGTAATAATAAAAG

Prim.cons.                       TAAACATCAACAACGCTTGCCCAATATTCAATAAAAATTTACTATTAGTAATAATAAAAG

                                         70        80        90       100       110       120
                                          |         |         |         |         |         |
PnapiCcBV17jonction1sample25     ------------------------------------------------------------
PnapiCcBV17jonction1sample28     ------------------------------------------------------------
PnapiCcBV17jonction1squenceNCB   ATTTACTATAAAATATAACACAAAAAAACAAGTAAATAGTGCCATTATTCTTTAATTTTT

Prim.cons.                       ATTTACTATAAAATATAACACAAAAAAACAAGTAAATAGTGCCATTATTCTTTAATTTTT

                                        130       140       150       160       170       180
                                          |         |         |         |         |         |
PnapiCcBV17jonction1sample25     ------------------------------------------------------------
PnapiCcBV17jonction1sample28     ------------------------------------------------------------
PnapiCcBV17jonction1squenceNCB   TACCGTATAAGCTTCAATATAATCAATTGTTTTTTAGCACTGGATGGGCCCGAGAATGCA

Prim.cons.                       TACCGTATAAGCTTCAATATAATCAATTGTTTTTTAGCACTGGATGGGCCCGAGAATGCA

                                        190       200       210       220       230       240
                                          |         |         |         |         |         |
PnapiCcBV17jonction1sample25     ------------------------------------------------------------
PnapiCcBV17jonction1sample28     ------------------------------------------------------------
PnapiCcBV17jonction1squenceNCB   GTGGTACCAAACGATGCAAAAGTCTTGGTGTCTTCGCAAGTGGTTGCGAATCCAGACGGG

Prim.cons.                       GTGGTACCAAACGATGCAAAAGTCTTGGTGTCTTCGCAAGTGGTTGCGAATCCAGACGGG

                                        250       260       270       280       290       300
                                          |         |         |         |         |         |
PnapiCcBV17jonction1sample25     ------------------------------------------------------------
PnapiCcBV17jonction1sample28     ------------------------------------------------------------
PnapiCcBV17jonction1squenceNCB   AAAGTTCGGGAAAGTCACGAACGAAAAGTCCTTACCAGAAATATTACAGATAGAGTGAGG

Prim.cons.                       AAAGTTCGGGAAAGTCACGAACGAAAAGTCCTTACCAGAAATATTACAGATAGAGTGAGG

                                        310       320       330       340       350       360
                                          |         |         |         |         |         |
PnapiCcBV17jonction1sample25     ------------------------------------------------------------
PnapiCcBV17jonction1sample28     ------------------------------------------------------------
PnapiCcBV17jonction1squenceNCB   GAGACAGAAGAGAAAATACATACTGGCGATACTACGCACGAGGTTAGTTTCTTGCACAGT

Prim.cons.                       GAGACAGAAGAGAAAATACATACTGGCGATACTACGCACGAGGTTAGTTTCTTGCACAGT

                                        370       380       390       400       410       420
                                          |         |         |         |         |         |
PnapiCcBV17jonction1sample25     ------------------------------------------------------------
PnapiCcBV17jonction1sample28     ------------------------------------------------------------
PnapiCcBV17jonction1squenceNCB   TTGTAATTTATCGCTTAAATTATAAGTATTTTTAAGAAAGATTTCTCATTGAAAAAGTTA

Prim.cons.                       TTGTAATTTATCGCTTAAATTATAAGTATTTTTAAGAAAGATTTCTCATTGAAAAAGTTA

                                        430       440       450       460       470       480
                                          |         |         |         |         |         |
PnapiCcBV17jonction1sample25     ------------------------------------------------------------
PnapiCcBV17jonction1sample28     ------------------------------------------------------------
PnapiCcBV17jonction1squenceNCB   TAAGTTGTTAGAACACATCCAATAATAAGGTTATCGGAAATTATAAATAAATTGAATGAG

Prim.cons.                       TAAGTTGTTAGAACACATCCAATAATAAGGTTATCGGAAATTATAAATAAATTGAATGAG

                                        490       500       510       520       530       540
                                          |         |         |         |         |         |
PnapiCcBV17jonction1sample25     ------------------------------------------------------------
PnapiCcBV17jonction1sample28     ------------------------------------------------------------
PnapiCcBV17jonction1squenceNCB   AGATCATTTTAAGAGATTGAACAATTTAAAAAATATAATAATATTTAACTCATTAAACAT

Prim.cons.                       AGATCATTTTAAGAGATTGAACAATTTAAAAAATATAATAATATTTAACTCATTAAACAT

                                        550       560       570       580       590       600
                                          |         |         |         |         |         |
PnapiCcBV17jonction1sample25     ------------------------------------------------------------
PnapiCcBV17jonction1sample28     ------------------------------------------------------------
PnapiCcBV17jonction1squenceNCB   TAAATAATAACAACAGCAATATTAAACGTTAACCCTGAAACGATATGGTAAATTTTGGCC

Prim.cons.                       TAAATAATAACAACAGCAATATTAAACGTTAACCCTGAAACGATATGGTAAATTTTGGCC

                                        610       620       630       640       650       660
                                          |         |         |         |         |         |
PnapiCcBV17jonction1sample25     ------------------------------------------------------------
PnapiCcBV17jonction1sample28     ------------------------------------------------------------
PnapiCcBV17jonction1squenceNCB   AATTCAAGAAACGTTCAGGCGACACACATAACCCTATAGGTAGTATGTAAGTAAGCCATT

Prim.cons.                       AATTCAAGAAACGTTCAGGCGACACACATAACCCTATAGGTAGTATGTAAGTAAGCCATT

                                        670       680       690       700       710       720
                                          |         |         |         |         |         |
PnapiCcBV17jonction1sample25     ------------------------------------------------------------
PnapiCcBV17jonction1sample28     ------------------------------------------------------------
PnapiCcBV17jonction1squenceNCB   TTTAATCTAGACGGATCTAACGTTAATAAGCTAAATGAAATACTGCTGGTCTGTTGGAGC

Prim.cons.                       TTTAATCTAGACGGATCTAACGTTAATAAGCTAAATGAAATACTGCTGGTCTGTTGGAGC

                                        730       740       750       760       770       780
                                          |         |         |         |         |         |
PnapiCcBV17jonction1sample25     ------------------------------------------------------------
PnapiCcBV17jonction1sample28     ------------------------------------------------------------
PnapiCcBV17jonction1squenceNCB   AGAACCATAGTTTATTTATAAAAGTTGCCACATCGACATAATTAAGAATTTAAAAATACG

Prim.cons.                       AGAACCATAGTTTATTTATAAAAGTTGCCACATCGACATAATTAAGAATTTAAAAATACG

                                        790       800       810       820       830       840
                                          |         |         |         |         |         |
PnapiCcBV17jonction1sample25     ------------------------------------------------------------
PnapiCcBV17jonction1sample28     ------------------------------------------------------------
PnapiCcBV17jonction1squenceNCB   GAATATTATCGTGGCTCTATACATAATTACAAGTATAAGGCCAGTCAGGACTATAATGTG

Prim.cons.                       GAATATTATCGTGGCTCTATACATAATTACAAGTATAAGGCCAGTCAGGACTATAATGTG

                                        850       860       870       880       890       900
                                          |         |         |         |         |         |
PnapiCcBV17jonction1sample25     ------------------------------------------------------------
PnapiCcBV17jonction1sample28     ------------------------------------------------------------
PnapiCcBV17jonction1squenceNCB   CATTACCATATTCTGAATATAGTAAATCTCTAGAGATATTCAATTTAGTAAGCATGTGGT

Prim.cons.                       CATTACCATATTCTGAATATAGTAAATCTCTAGAGATATTCAATTTAGTAAGCATGTGGT

                                        910       920       930       940       950       960
                                          |         |         |         |         |         |
PnapiCcBV17jonction1sample25     ------------------------------------------------------------
PnapiCcBV17jonction1sample28     ------------------------------------------------------------
PnapiCcBV17jonction1squenceNCB   GCATTTAAATCACACTGCAATAAAGGCGCCAAAGTAGGGAAGTGCGTTTGACTTTCGTAT

Prim.cons.                       GCATTTAAATCACACTGCAATAAAGGCGCCAAAGTAGGGAAGTGCGTTTGACTTTCGTAT

                                        970       980       990      1000      1010      1020
                                          |         |         |         |         |         |
PnapiCcBV17jonction1sample25     --------------------------------CGTAATATGAGTAATACCATTTCTTTTT
PnapiCcBV17jonction1sample28     ---------------------------------GTAATATGAGTAATACCATTTCTTTTT
PnapiCcBV17jonction1squenceNCB   AATATTAGTAATACTTGCTTGTAATATTAACGCGTAATATGAGTAATACTATTTATTTTT
                                                                  **************** **** *****
Prim.cons.                       AATATTAGTAATACTTGCTTGTAATATTAACGCGTAATATGAGTAATACCATTTCTTTTT

                                       1030      1040      1050      1060      1070      1080
                                          |         |         |         |         |         |
PnapiCcBV17jonction1sample25     A------AGTTACATTAGTACTTGGAAAGGAAATCAATTATTTTCTGTAAACGTTTTAAC
PnapiCcBV17jonction1sample28     A------AGTTACATTAGTACTTGGAAAGGAAATCAATTATTTTCTGTAAACGTTTTAAC
PnapiCcBV17jonction1squenceNCB   ATATAAAAGTTACATTAGTACTTGGAAAGGAAATTAATAATTTTCTGTAAACGTTTTAAT
                                 *      *************************** *** ********************
Prim.cons.                       ATATAAAAGTTACATTAGTACTTGGAAAGGAAATCAATTATTTTCTGTAAACGTTTTAAC


                                       1090      1100      1110      1120      1130      1140
                                          |         |         |         |         |         |
PnapiCcBV17jonction1sample25     TATTTGACGCAAAAACTT-CGCGCGAGTGGTAGGATCAATAGTATAAAAACAACAAATGA
PnapiCcBV17jonction1sample28     TATTTGACGCAAAAACTT-CGCGCGAGTGGTAGGATCAATAGTATAAAAACAACAAATGA
PnapiCcBV17jonction1squenceNCB   TACTTGACGCAAAAACTTACGCGCGAGTGGTACTATCAATAGTATAAAAACAACAAATGA
                                 ** *************** *************  **************************
Prim.cons.                       TATTTGACGCAAAAACTTACGCGCGAGTGGTAGGATCAATAGTATAAAAACAACAAATGA

                                       1150      1160      1170      1180      1190      1200
                                          |         |         |         |         |         |
PnapiCcBV17jonction1sample25     TTAAATCTTACCATTTTCTATACATATTTTTAAATAAAAACTTAGCTTTGAACAGTAATG
PnapiCcBV17jonction1sample28     TTAAATCTTACCATTTTCTATACATATTATTAAATAAAAACTTAGCTTTGAACAGTAATG
PnapiCcBV17jonction1squenceNCB   TTAAATCTTACCATTTTCTATACATATTATTAAATAAAAACTTAGCTTTGAACAGTAATG
                                 **************************** *******************************
Prim.cons.                       TTAAATCTTACCATTTTCTATACATATTATTAAATAAAAACTTAGCTTTGAACAGTAATG

                                       1210      1220      1230      1240      1250      1260
                                          |         |         |         |         |         |
PnapiCcBV17jonction1sample25     GTCGCGATCTTGAAATTTTTAGACTTAAAAAATATTCAGCTGTAGAGTCGATGACGTCAC
PnapiCcBV17jonction1sample28     GTCGCGATCTTGAAATTTTTAGACTTAAAAAATATTCAGCTGTAGAGTCGATGACGTCAC
PnapiCcBV17jonction1squenceNCB   GTCGCGATCTTGAAATTTTTAGACTTAAAAAATATTCAGCTGTAGAGTCGATGACGTCAC
                                 ************************************************************
Prim.cons.                       GTCGCGATCTTGAAATTTTTAGACTTAAAAAATATTCAGCTGTAGAGTCGATGACGTCAC

                                       1270      1280      1290      1300      1310      1320
                                          |         |         |         |         |         |
PnapiCcBV17jonction1sample25     GACGGAATTAATGTCACGGTAAGTCCGTGTTCTCTATGGCGTCATAAAAAGTTCAATATA
PnapiCcBV17jonction1sample28     GACGGAATTAATGTCACGGTAAGTCCGTGTTCTCTATGGCGTCATAAAAAGTTCAATATA
PnapiCcBV17jonction1squenceNCB   GACGGAATTAATGTCACGGTAAGTCCGTGTTCTCTATGGCGTCATAAAAAGTTCAATATA
                                 ************************************************************
Prim.cons.                       GACGGAATTAATGTCACGGTAAGTCCGTGTTCTCTATGGCGTCATAAAAAGTTCAATATA

                                       1330      1340      1350      1360      1370      1380
                                          |         |         |         |         |         |
PnapiCcBV17jonction1sample25     GAGGGCACGCTGAGTATTTAGCTCCAGTTTATGTGTATTCGTTACGTAGGTAAGGTCGTT
PnapiCcBV17jonction1sample28     GAGGGCACGCTGAGTATTTAGCTCCAGTTTATGTGTATTCGTTACGTAGGTAAGGTCGTT
PnapiCcBV17jonction1squenceNCB   GAGGGCACGCTGAGTATTTAGCTCCAGTTTATGTGTATTCGTTACGTAGGTAAGGTCGTT
                                 ************************************************************
Prim.cons.                       GAGGGCACGCTGAGTATTTAGCTCCAGTTTATGTGTATTCGTTACGTAGGTAAGGTCGTT

                                       1390      1400      1410      1420      1430      1440
                                          |         |         |         |         |         |
PnapiCcBV17jonction1sample25     GACTCCTATGATACGGCCCTGACCTTTA--------------------------------
PnapiCcBV17jonction1sample28     GACTCCTATGATACGGCCCTGACCTTTA--------------------------------
PnapiCcBV17jonction1squenceNCB   GACTCCTATGATACGGCCCTGACCTTTATTATGTAGTCCTGATAAAGGTGAATTTACATT
                                 ****************************
Prim.cons.                       GACTCCTATGATACGGCCCTGACCTTTATTATGTAGTCCTGATAAAGGTGAATTTACATT

                                       1450      1460      1470      1480      1490      1500
                                          |         |         |         |         |         |
PnapiCcBV17jonction1sample25     ------------------------------------------------------------
PnapiCcBV17jonction1sample28     ------------------------------------------------------------
PnapiCcBV17jonction1squenceNCB   GTGAACATGTTGTTTACTTCTGAGGTATAGTTATTGAGTAAGGTACACACGAAGTGTATA

Prim.cons.                       GTGAACATGTTGTTTACTTCTGAGGTATAGTTATTGAGTAAGGTACACACGAAGTGTATA
```

>P. napi CcBV17 jonction2 – NCBI accession number (start-end coordinates): DWAF01019933.1 (542-1831)

TAGCCCGCGCGAGAGGATGTATCTATATTCGACATAAATACCGATCTGTGAAAATGATCGGCGACGCAACCGTTCGTACTTACTCCACATTCCACTTAATATCTGCTCCTTGAGACGCGATTAACGGATTCGAATTGAACCTTGACTTCAGTACGTTTTAGTACATGTTTAGAAAAACGGGAAATAACTAAAATAGTTAAACGACGGAAGCAGCTATTAAATGGAGTTAGTGAAGTGGCGGCATCGTAACACGAGTTTCGGTTCACGTGACACGTTGTNACCGATTTGCGTAGGTTTACTAAAGAATTTATAAGCAAAAATACGTCTAACTATAGAAACTTCGACAAAGTCGCCGATCCAAATCCGACTTTTTCTTAGCTTCTGTGGCATAGGACGGNTCCGTAAGTCACTGTCGGCCACAAACGAGACGTGTCTAAAGTACTCCGGTAATCGTACCGTTTCGCTCACTTATATTGTGTATGTTTATAAATAAGAGCGAGACGACGTGCGAGTTGGACGTGTATCTTTTAGATGAATAAGGCTAATTTACTTTATACCATTTAATTCTTGGGTAATTGTAATTTAATAATAATTCTAAAATAATGCTGTAAAGTTGTACGTAATAATATATTGTATCTATGATTGTGGTATACGTAATAATTTAAAAATTAAACAATTCTATAATGATTCATCTTGGATGTGATTAAGTAGGTAGTTCTTGAATACGAACCGCCGCTAGATGGCTCTGTTTTAATTATTGTCGTCATCTATCGGTCGGTAGAGATATAATTTACACAGCTAATGTACAGTCAAACAATTTCCACATTGTAACGCCACCTATTGGCGAGTAGTAGTAAACATAACACTGCCATCTATTGTACAATAGAATAAACTTCCTATTTGCTACTTTTCTATGTAAAAACGAGTGCAATAGTTTTGATGTCAATTGTGAAGGAAATAATTTTAGTCTNACACGAACATCGCAGCCTAAGAGAAATAAGAATGTAATTTATTTTATATCCTAACTGTAGATAAATAAAACGACTTTTATGTACAAATGTTGCCTTTTTTTTTACAAAGTTCCTCTTTTAGTTACAAAATGCACCGACCTCCCTTCAATAGTCTTTAACTAGACGTAGGACGGTTATATATCTGAATTGGCACATCCGGGGATCTGTTCAATGTCGAAATTTAAGAATNTAATTCCAAAAAAGATTGAAAAATTCTTTTCTACAAGCGAAACTGGCCATTATGTNGAACCTTGATTTGTATAATGCAGAAGNGTCTCAA

>P. napi CcBV17 jonction2 sample21

TATATCCTACTGTAGATAAATAAAACGACTTTTATGTACAAATGTTGCCTTTTCTTTACAAAGTTCCTCTTTTAGTTACAAAATGCACCGAACTCCCTTCAATAGTCTTTAACTAGACGTAGGACGGTTATATATCTGAATTGGCACATCCAGAGATCTGTTCAATGTCGAAATTTAAGAATTTAATTCCAAAAAAGATTGAAAAATTCTTTTCTACAAGCGAAACTGG

>P. napi CcBV17 jonction2 sample25

TTTAATCCTACTGTAGATAAATAAAACGACTTTTATGTACAAATGTTGCCTTTTTTTACAAAGTTCCTCTTTTAGTTACAAAATGCACCGACCTCCCTTCAATAGTCTTTAACTAGACGTAGGACGGTTATATATCTGAATTGGCACTTCCAGGGATCTGTTCAATGTCGAAATTTAAGAATTTAATTCCAAAAAAGATTGAAAAATTCTTTTCTACAAGCGAAACTGG

Multiple alignment

```
                                         10        20        30        40        50        60
                                          |         |         |         |         |         |
PnapiCcBV17jonction2squenceNCB   TAGCCCGCGCGAGAGGATGTATCTATATTCGACATAAATACCGATCTGTGAAAATGATCG
PnapiCcBV17jonction2sample25     ------------------------------------------------------------
PnapiCcBV17jonction2sample21     ------------------------------------------------------------

Prim.cons.                       TAGCCCGCGCGAGAGGATGTATCTATATTCGACATAAATACCGATCTGTGAAAATGATCG


                                         70        80        90       100       110       120
                                          |         |         |         |         |         |
PnapiCcBV17jonction2squenceNCB   GCGACGCAACCGTTCGTACTTACTCCACATTCCACTTAATATCTGCTCCTTGAGACGCGA
PnapiCcBV17jonction2sample25     ------------------------------------------------------------
PnapiCcBV17jonction2sample21     ------------------------------------------------------------

Prim.cons.                       GCGACGCAACCGTTCGTACTTACTCCACATTCCACTTAATATCTGCTCCTTGAGACGCGA

                                        130       140       150       160       170       180
                                          |         |         |         |         |         |
PnapiCcBV17jonction2squenceNCB   TTAACGGATTCGAATTGAACCTTGACTTCAGTACGTTTTAGTACATGTTTAGAAAAACGG
PnapiCcBV17jonction2sample25     ------------------------------------------------------------
PnapiCcBV17jonction2sample21     ------------------------------------------------------------

Prim.cons.                       TTAACGGATTCGAATTGAACCTTGACTTCAGTACGTTTTAGTACATGTTTAGAAAAACGG

                                        190       200       210       220       230       240
                                          |         |         |         |         |         |
PnapiCcBV17jonction2squenceNCB   GAAATAACTAAAATAGTTAAACGACGGAAGCAGCTATTAAATGGAGTTAGTGAAGTGGCG
PnapiCcBV17jonction2sample25     ------------------------------------------------------------
PnapiCcBV17jonction2sample21     ------------------------------------------------------------

Prim.cons.                       GAAATAACTAAAATAGTTAAACGACGGAAGCAGCTATTAAATGGAGTTAGTGAAGTGGCG

                                        250       260       270       280       290       300
                                          |         |         |         |         |         |
PnapiCcBV17jonction2squenceNCB   GCATCGTAACACGAGTTTCGGTTCACGTGACACGTTGTNACCGATTTGCGTAGGTTTACT
PnapiCcBV17jonction2sample25     ------------------------------------------------------------
PnapiCcBV17jonction2sample21     ------------------------------------------------------------

Prim.cons.                       GCATCGTAACACGAGTTTCGGTTCACGTGACACGTTGTNACCGATTTGCGTAGGTTTACT

                                        310       320       330       340       350       360
                                          |         |         |         |         |         |
PnapiCcBV17jonction2squenceNCB   AAAGAATTTATAAGCAAAAATACGTCTAACTATAGAAACTTCGACAAAGTCGCCGATCCA
PnapiCcBV17jonction2sample25     ------------------------------------------------------------
PnapiCcBV17jonction2sample21     ------------------------------------------------------------

Prim.cons.                       AAAGAATTTATAAGCAAAAATACGTCTAACTATAGAAACTTCGACAAAGTCGCCGATCCA

                                        370       380       390       400       410       420
                                          |         |         |         |         |         |
PnapiCcBV17jonction2squenceNCB   AATCCGACTTTTTCTTAGCTTCTGTGGCATAGGACGGNTCCGTAAGTCACTGTCGGCCAC
PnapiCcBV17jonction2sample25     ------------------------------------------------------------
PnapiCcBV17jonction2sample21     ------------------------------------------------------------

Prim.cons.                       AATCCGACTTTTTCTTAGCTTCTGTGGCATAGGACGGNTCCGTAAGTCACTGTCGGCCAC

                                        430       440       450       460       470       480
                                          |         |         |         |         |         |
PnapiCcBV17jonction2squenceNCB   AAACGAGACGTGTCTAAAGTACTCCGGTAATCGTACCGTTTCGCTCACTTATATTGTGTA
PnapiCcBV17jonction2sample25     ------------------------------------------------------------
PnapiCcBV17jonction2sample21     ------------------------------------------------------------

Prim.cons.                       AAACGAGACGTGTCTAAAGTACTCCGGTAATCGTACCGTTTCGCTCACTTATATTGTGTA

                                        490       500       510       520       530       540
                                          |         |         |         |         |         |
PnapiCcBV17jonction2squenceNCB   TGTTTATAAATAAGAGCGAGACGACGTGCGAGTTGGACGTGTATCTTTTAGATGAATAAG
PnapiCcBV17jonction2sample25     ------------------------------------------------------------
PnapiCcBV17jonction2sample21     ------------------------------------------------------------

Prim.cons.                       TGTTTATAAATAAGAGCGAGACGACGTGCGAGTTGGACGTGTATCTTTTAGATGAATAAG

                                        550       560       570       580       590       600
                                          |         |         |         |         |         |
PnapiCcBV17jonction2squenceNCB   GCTAATTTACTTTATACCATTTAATTCTTGGGTAATTGTAATTTAATAATAATTCTAAAA
PnapiCcBV17jonction2sample25     ------------------------------------------------------------
PnapiCcBV17jonction2sample21     ------------------------------------------------------------

Prim.cons.                       GCTAATTTACTTTATACCATTTAATTCTTGGGTAATTGTAATTTAATAATAATTCTAAAA


                                        610       620       630       640       650       660
                                          |         |         |         |         |         |
PnapiCcBV17jonction2squenceNCB   TAATGCTGTAAAGTTGTACGTAATAATATATTGTATCTATGATTGTGGTATACGTAATAA
PnapiCcBV17jonction2sample25     ------------------------------------------------------------
PnapiCcBV17jonction2sample21     ------------------------------------------------------------

Prim.cons.                       TAATGCTGTAAAGTTGTACGTAATAATATATTGTATCTATGATTGTGGTATACGTAATAA

                                        670       680       690       700       710       720
                                          |         |         |         |         |         |
PnapiCcBV17jonction2squenceNCB   TTTAAAAATTAAACAATTCTATAATGATTCATCTTGGATGTGATTAAGTAGGTAGTTCTT
PnapiCcBV17jonction2sample25     ------------------------------------------------------------
PnapiCcBV17jonction2sample21     ------------------------------------------------------------

Prim.cons.                       TTTAAAAATTAAACAATTCTATAATGATTCATCTTGGATGTGATTAAGTAGGTAGTTCTT

                                        730       740       750       760       770       780
                                          |         |         |         |         |         |
PnapiCcBV17jonction2squenceNCB   GAATACGAACCGCCGCTAGATGGCTCTGTTTTAATTATTGTCGTCATCTATCGGTCGGTA
PnapiCcBV17jonction2sample25     ------------------------------------------------------------
PnapiCcBV17jonction2sample21     ------------------------------------------------------------

Prim.cons.                       GAATACGAACCGCCGCTAGATGGCTCTGTTTTAATTATTGTCGTCATCTATCGGTCGGTA

                                        790       800       810       820       830       840
                                          |         |         |         |         |         |
PnapiCcBV17jonction2squenceNCB   GAGATATAATTTACACAGCTAATGTACAGTCAAACAATTTCCACATTGTAACGCCACCTA
PnapiCcBV17jonction2sample25     ------------------------------------------------------------
PnapiCcBV17jonction2sample21     ------------------------------------------------------------

Prim.cons.                       GAGATATAATTTACACAGCTAATGTACAGTCAAACAATTTCCACATTGTAACGCCACCTA

                                        850       860       870       880       890       900
                                          |         |         |         |         |         |
PnapiCcBV17jonction2squenceNCB   TTGGCGAGTAGTAGTAAACATAACACTGCCATCTATTGTACAATAGAATAAACTTCCTAT
PnapiCcBV17jonction2sample25     ------------------------------------------------------------
PnapiCcBV17jonction2sample21     ------------------------------------------------------------

Prim.cons.                       TTGGCGAGTAGTAGTAAACATAACACTGCCATCTATTGTACAATAGAATAAACTTCCTAT

                                        910       920       930       940       950       960
                                          |         |         |         |         |         |
PnapiCcBV17jonction2squenceNCB   TTGCTACTTTTCTATGTAAAAACGAGTGCAATAGTTTTGATGTCAATTGTGAAGGAAATA
PnapiCcBV17jonction2sample25     ------------------------------------------------------------
PnapiCcBV17jonction2sample21     ------------------------------------------------------------

Prim.cons.                       TTGCTACTTTTCTATGTAAAAACGAGTGCAATAGTTTTGATGTCAATTGTGAAGGAAATA

                                        970       980       990      1000      1010      1020
                                          |         |         |         |         |         |
PnapiCcBV17jonction2squenceNCB   ATTTTAGTCTNACACGAACATCGCAGCCTAAGAGAAATAAGAATGTAATTTATTTTATAT
PnapiCcBV17jonction2sample25     -----------------------------------------------------TTTA-AT
PnapiCcBV17jonction2sample21     -------------------------------------------------------TATAT
                                                                                        ** **
Prim.cons.                       ATTTTAGTCTNACACGAACATCGCAGCCTAAGAGAAATAAGAATGTAATTTATTTTATAT

                                       1030      1040      1050      1060      1070      1080
                                          |         |         |         |         |         |
PnapiCcBV17jonction2squenceNCB   CCTAACTGTAGATAAATAAAACGACTTTTATGTACAAATGTTGCCTTTTTTTTTACAAAG
PnapiCcBV17jonction2sample25     CCTA-CTGTAGATAAATAAAACGACTTTTATGTACAAATGTTGCCTTTTTTT--ACAAAG
PnapiCcBV17jonction2sample21     CCTA-CTGTAGATAAATAAAACGACTTTTATGTACAAATGTTGCCTTTTCTTT-ACAAAG
                                 **** ******************************************** **  ******
Prim.cons.                       CCTAACTGTAGATAAATAAAACGACTTTTATGTACAAATGTTGCCTTTTTTTTTACAAAG

                                       1090      1100      1110      1120      1130      1140
                                          |         |         |         |         |         |
PnapiCcBV17jonction2squenceNCB   TTCCTCTTTTAGTTACAAAATGCACCGACCTCCCTTCAATAGTCTTTAACTAGACGTAGG
PnapiCcBV17jonction2sample25     TTCCTCTTTTAGTTACAAAATGCACCGACCTCCCTTCAATAGTCTTTAACTAGACGTAGG
PnapiCcBV17jonction2sample21     TTCCTCTTTTAGTTACAAAATGCACCGAACTCCCTTCAATAGTCTTTAACTAGACGTAGG
                                 **************************** *******************************
Prim.cons.                       TTCCTCTTTTAGTTACAAAATGCACCGACCTCCCTTCAATAGTCTTTAACTAGACGTAGG


                                       1150      1160      1170      1180      1190      1200
                                          |         |         |         |         |         |
PnapiCcBV17jonction2squenceNCB   ACGGTTATATATCTGAATTGGCACATCCGGGGATCTGTTCAATGTCGAAATTTAAGAATN
PnapiCcBV17jonction2sample25     ACGGTTATATATCTGAATTGGCACTTCCAGGGATCTGTTCAATGTCGAAATTTAAGAATT
PnapiCcBV17jonction2sample21     ACGGTTATATATCTGAATTGGCACATCCAGAGATCTGTTCAATGTCGAAATTTAAGAATT
                                 ************************ *** * ****************************
Prim.cons.                       ACGGTTATATATCTGAATTGGCACATCCAGGGATCTGTTCAATGTCGAAATTTAAGAATT

                                       1210      1220      1230      1240      1250      1260
                                          |         |         |         |         |         |
PnapiCcBV17jonction2squenceNCB   TAATTCCAAAAAAGATTGAAAAATTCTTTTCTACAAGCGAAACTGGCCATTATGTNGAAC
PnapiCcBV17jonction2sample25     TAATTCCAAAAAAGATTGAAAAATTCTTTTCTACAAGCGAAACTGG--------------
PnapiCcBV17jonction2sample21     TAATTCCAAAAAAGATTGAAAAATTCTTTTCTACAAGCGAAACTGG--------------
                                 **********************************************
Prim.cons.                       TAATTCCAAAAAAGATTGAAAAATTCTTTTCTACAAGCGAAACTGGCCATTATGTNGAAC

                                       1270      1280      1290
                                          |         |         |
PnapiCcBV17jonction2squenceNCB   CTTGATTTGTATAATGCAGAAGNGTCTCAA
PnapiCcBV17jonction2sample25     ------------------------------
PnapiCcBV17jonction2sample21     ------------------------------

Prim.cons.                       CTTGATTTGTATAATGCAGAAGNGTCTCAA
```

>P. napi HdIV12 jonction1 – NCBI accession number (start-end coordinates): CAJQFU010000061.1 (476573-478187)

AACGTATTAAGGCGAAACGAAATTCGCGGGGCAGCTAGTAATTTATTCAAAAAAATAAAGAGTACGTGGTGAGCGGATGGGCACAGAAAGTAAGTTTCATATACTTACTGTGGATATCCTAAAAGATTCGCCAAGATTTATGTGAGAGAGAGAGTTCTTCTTGAGATATGAAATAGGTTTAGGAATAAGGGTTGGAATATAAGAGANGGAGAGCGTGAGCATCACGCCGAAACGGTCCTTAGGCCTAAAACAGCGTGGCAACAAACACGGTACTTTAATCGTTTAAAATAAGGTCAAAAGAAATCAATTAAACCGATGCATTTGAATAATCGAGTCATCGTAAGAATACTCTAGTTCGAGTTTTGCACGGAACTAATATANAGCATTCCTCAAAAGCGTTTCGATTTCAAACGTTCCAGAAAACACGGAGCAAAGTGNAGCGTTTGTGCTCGGGTTTTAGATTCGAAAAAAATAATTTGCAGCCCGTTCACTTTTTTATTCTTTTGCGACACTGCGTATACGGATTTGATTTGTTTCATGAAACCGTAAAGGAAANCTTGCGCTCGATGCCGAATCAAATGTCCGGCTATATTTCAATTATTCAAAGCTTTGATTTTCANTTAAGTAGACTGCGGCTGGTTGCGTTCTATGGAGAGACTATTGACATACTTTATATTTTTAATACGATTAAACGATAGGTATTATCTGTGGTNTTTAACGGTTATTTGTGTAGGATCGACCTTACAATACAATAAATATTTTGTGGCGCGTGTATAACACGCGATCGACACGTCACTCTCCCCCCTTAGTGATACCTAAAAATCGCTTCTGCGCAGGCAATGACACTTGTTCATGATGTATGCCGGTATCTGTACTTTAGAATATTTTCGGGAAATTATGAATATTTTATTTGAATTTTAAAGCCATACAAAGCGCACTAATTAAAAATTATGGAATTCACCATTATAGCCATAAAAAGAACGACAAAAAATAGTCTAACTTAATAATAAATTCTACAAACGATATCACATATAAAAGATAATTTTAGCCCTCCACCCTTAAAGTCTAGACTCTAGGCAATTGCTCTATATATTTCGCTACCTCGATTCACATAAAATTGTAGCAATAACCATAGAAGACCTTTCCATTATATCCCTTAACTAAACTACAGGGTGGGTTTTTGTTTATACAGCAAAAGTATAAATGTGATAATATTTTACANACGTCCGAAATTTGTGTAAATGGGGAATGTAAAACAAAGCGATACAGTCATGGTCATAAGACCGTACGGGGCAAAAATTCAAAATGGTGGACCGGGGNAAGGGGAGGGTGATACCAATGTATGCAGCGGAGAAGTGACTCTGATGCATAGTGGACTCGAGAAAAGTATGATTGTTTAAGAAGTTCTAGGTTTTGCTATGTGAAAATGAGANCTAGATTCGAGTTCCCCTAACCTTAGGTCTCTGTGAATGCGAGCGAAAAGCAAAAACACTTAACTGCTAACCTATGAGTGCACATTGGTTCTAGGTAGCAAAATTGATATCGGCAAATTGCCACGTAATGTCANGACATTAGAGCGACCTACCGTGACGGCGATAACGCTGTTATCGCTGAATTTAAATGGTATCAATGTGAATTTTCTGTGATCCAATTAAAAAAGTTATCNGGCTTAACTAACCAATCTTATGGGGTTCATTATGATATTAATCGTTCGGGGAAATAAAACAATACNCCATAGTTCACGACGTCATACCTACGGTGAAACACAGGCACACCTATGAAGATGAGTGTTCATTATCCAACGATTGTGTGGATGGATGAATCAATGTAAACCTAAGTTTCTCAAGTTCCGTGGTGAACTTCTTTGCGGACATGAGTGCACGCTACAACGCTGTGGTGGAATGCTCGGTGTAACGGACCCTTCACTGAATATAACATATGTAAGACTGCTGGTCACGTAGCAAGACTCCCTGCATGCTCCTCATGCCAGGCAGCCTGCAGTTACATTTTAAACTCTTTAGTTGCATTTTTGTACCATAGCGAGTATCTCCTGAAGCTTACCTGTATTTTTTTCCTATANGGAGCATGATTGTGCAAGCTAATCGGAAAGTTAACACAGTCTATGACATCGGTGCCATCTTCAACATTTTTCCCTTCTTCCGTTGAAAATAATTGGGTAATGCCGATATGTATTTTCCGTTAACATCATCAATATAGCCCAATTTGTATCGCAAAATACTTTTCTCTACCTAAATTCGATATCATTAGCGAGTTTTGTTAAAAACAAAAAAAATCCTGATAACTATAAAATGCTAACCGGAACACATTAGGTAGGTTTCGAATTAGTATAAATTGAGGGAAGGACCTATACATTNAATCATTAGCATCATTACCCCCGACATAATCACCAGGAGCTATACAGTCAATNAACACTAATACCCTCGACATAGTCACCATGTGGGGGAACAATAGTGGAAACTCCGGAGAAGCCGCTCGTCGGTACGCCGAGGAAGAGAAGTGCAGACAAGACAAGGAGCGTGATGAGAGGAAGCGTCAGCAGGAAGAGCAAGAGCGAATTGCGGAGCGCGTGAGGCAGTATG

>P. napi HdIV12 jonction1 sample5

TGCTCTATATATTTCGCTACCTCGATTCACATAAAATTGTAGCAATAACCATAGAAGACCTTTCCATTATATCCCTTAACTACAGGGTGGGTTTTTGTTTATACAGCAAAATTATAAATGTGATAATATTTTACACACGTCCGAAATTTGTGTAAATGGGGAATGTAAAACAAAGAGATACAGTCATGGTCATAAGACCGTACGGGGCAAAAATTCAAAATGGTGGACCGGGGAAAGGGGAGGGTGATACCAATGTATGCAGCGGAGAAGTGACTCTGATGCATAGTGGACTCGGGAGAAGTATGATTGTTTAAGAAGTTCTAGGTTTTGCTATGTGAAAATGAGACCTAGATTCGAGTTCCCCTAACCTTAGG

>P. napi HdIV12 jonction1 sample21

ATTTCGCTACCTCGATTCACATAAAATTGTAGCAATAACCATAGAAGACCTTTCCATTATATCCCTTAACTACAGGGTGGGTTTTTGATTATACAGCAAAAGTATAAATGTCATAATATTTTACACACGTCCGAAATTTGTGTAAATGGGGAATGTAAAGCAAAGCGATACAGTCATGGTCATAAGACCGTACGGGGAAAAAATTCAAAAGGGTGGGGTAAGGGGAGGGTGATACCAATGTATGCAGCGGAGAAGTGACTCTGATGCATAGTGGACTCGGGAAAAGTATGATTGTTTAAGAAGTTCTAGGTTTTGCTATGTGAAAATGAGTCCTAGATTCGAGTTCCCCTAACCTTAGG

Multiple alignment

```
                                         10        20        30        40        50        60
                                          |         |         |         |         |         |
PnapiHdIV12jonction1squenceNCB   AACGTATTAAGGCGAAACGAAATTCGCGGGGCAGCTAGTAATTTATTCAAAAAAATAAAG
PnapiHdIV12jonction1sample5      ------------------------------------------------------------
PnapiHdIV12jonction1sample21     ------------------------------------------------------------

Prim.cons.                       AACGTATTAAGGCGAAACGAAATTCGCGGGGCAGCTAGTAATTTATTCAAAAAAATAAAG

                                         70        80        90       100       110       120
                                          |         |         |         |         |         |
PnapiHdIV12jonction1squenceNCB   AGTACGTGGTGAGCGGATGGGCACAGAAAGTAAGTTTCATATACTTACTGTGGATATCCT
PnapiHdIV12jonction1sample5      ------------------------------------------------------------
PnapiHdIV12jonction1sample21     ------------------------------------------------------------

Prim.cons.                       AGTACGTGGTGAGCGGATGGGCACAGAAAGTAAGTTTCATATACTTACTGTGGATATCCT

                                        130       140       150       160       170       180
                                          |         |         |         |         |         |
PnapiHdIV12jonction1squenceNCB   AAAAGATTCGCCAAGATTTATGTGAGAGAGAGAGTTCTTCTTGAGATATGAAATAGGTTT
PnapiHdIV12jonction1sample5      ------------------------------------------------------------
PnapiHdIV12jonction1sample21     ------------------------------------------------------------

Prim.cons.                       AAAAGATTCGCCAAGATTTATGTGAGAGAGAGAGTTCTTCTTGAGATATGAAATAGGTTT

                                        190       200       210       220       230       240
                                          |         |         |         |         |         |
PnapiHdIV12jonction1squenceNCB   AGGAATAAGGGTTGGAATATAAGAGANGGAGAGCGTGAGCATCACGCCGAAACGGTCCTT
PnapiHdIV12jonction1sample5      ------------------------------------------------------------
PnapiHdIV12jonction1sample21     ------------------------------------------------------------

Prim.cons.                       AGGAATAAGGGTTGGAATATAAGAGANGGAGAGCGTGAGCATCACGCCGAAACGGTCCTT

                                        250       260       270       280       290       300
                                          |         |         |         |         |         |
PnapiHdIV12jonction1squenceNCB   AGGCCTAAAACAGCGTGGCAACAAACACGGTACTTTAATCGTTTAAAATAAGGTCAAAAG
PnapiHdIV12jonction1sample5      ------------------------------------------------------------
PnapiHdIV12jonction1sample21     ------------------------------------------------------------

Prim.cons.                       AGGCCTAAAACAGCGTGGCAACAAACACGGTACTTTAATCGTTTAAAATAAGGTCAAAAG


                                        310       320       330       340       350       360
                                          |         |         |         |         |         |
PnapiHdIV12jonction1squenceNCB   AAATCAATTAAACCGATGCATTTGAATAATCGAGTCATCGTAAGAATACTCTAGTTCGAG
PnapiHdIV12jonction1sample5      ------------------------------------------------------------
PnapiHdIV12jonction1sample21     ------------------------------------------------------------

Prim.cons.                       AAATCAATTAAACCGATGCATTTGAATAATCGAGTCATCGTAAGAATACTCTAGTTCGAG

                                        370       380       390       400       410       420
                                          |         |         |         |         |         |
PnapiHdIV12jonction1squenceNCB   TTTTGCACGGAACTAATATANAGCATTCCTCAAAAGCGTTTCGATTTCAAACGTTCCAGA
PnapiHdIV12jonction1sample5      ------------------------------------------------------------
PnapiHdIV12jonction1sample21     ------------------------------------------------------------

Prim.cons.                       TTTTGCACGGAACTAATATANAGCATTCCTCAAAAGCGTTTCGATTTCAAACGTTCCAGA

                                        430       440       450       460       470       480
                                          |         |         |         |         |         |
PnapiHdIV12jonction1squenceNCB   AAACACGGAGCAAAGTGNAGCGTTTGTGCTCGGGTTTTAGATTCGAAAAAAATAATTTGC
PnapiHdIV12jonction1sample5      ------------------------------------------------------------
PnapiHdIV12jonction1sample21     ------------------------------------------------------------

Prim.cons.                       AAACACGGAGCAAAGTGNAGCGTTTGTGCTCGGGTTTTAGATTCGAAAAAAATAATTTGC

                                        490       500       510       520       530       540
                                          |         |         |         |         |         |
PnapiHdIV12jonction1squenceNCB   AGCCCGTTCACTTTTTTATTCTTTTGCGACACTGCGTATACGGATTTGATTTGTTTCATG
PnapiHdIV12jonction1sample5      ------------------------------------------------------------
PnapiHdIV12jonction1sample21     ------------------------------------------------------------

Prim.cons.                       AGCCCGTTCACTTTTTTATTCTTTTGCGACACTGCGTATACGGATTTGATTTGTTTCATG

                                        550       560       570       580       590       600
                                          |         |         |         |         |         |
PnapiHdIV12jonction1squenceNCB   AAACCGTAAAGGAAANCTTGCGCTCGATGCCGAATCAAATGTCCGGCTATATTTCAATTA
PnapiHdIV12jonction1sample5      ------------------------------------------------------------
PnapiHdIV12jonction1sample21     ------------------------------------------------------------

Prim.cons.                       AAACCGTAAAGGAAANCTTGCGCTCGATGCCGAATCAAATGTCCGGCTATATTTCAATTA

                                        610       620       630       640       650       660
                                          |         |         |         |         |         |
PnapiHdIV12jonction1squenceNCB   TTCAAAGCTTTGATTTTCANTTAAGTAGACTGCGGCTGGTTGCGTTCTATGGAGAGACTA
PnapiHdIV12jonction1sample5      ------------------------------------------------------------
PnapiHdIV12jonction1sample21     ------------------------------------------------------------

Prim.cons.                       TTCAAAGCTTTGATTTTCANTTAAGTAGACTGCGGCTGGTTGCGTTCTATGGAGAGACTA

                                        670       680       690       700       710       720
                                          |         |         |         |         |         |
PnapiHdIV12jonction1squenceNCB   TTGACATACTTTATATTTTTAATACGATTAAACGATAGGTATTATCTGTGGTNTTTAACG
PnapiHdIV12jonction1sample5      ------------------------------------------------------------
PnapiHdIV12jonction1sample21     ------------------------------------------------------------

Prim.cons.                       TTGACATACTTTATATTTTTAATACGATTAAACGATAGGTATTATCTGTGGTNTTTAACG

                                        730       740       750       760       770       780
                                          |         |         |         |         |         |
PnapiHdIV12jonction1squenceNCB   GTTATTTGTGTAGGATCGACCTTACAATACAATAAATATTTTGTGGCGCGTGTATAACAC
PnapiHdIV12jonction1sample5      ------------------------------------------------------------
PnapiHdIV12jonction1sample21     ------------------------------------------------------------

Prim.cons.                       GTTATTTGTGTAGGATCGACCTTACAATACAATAAATATTTTGTGGCGCGTGTATAACAC

                                        790       800       810       820       830       840
                                          |         |         |         |         |         |
PnapiHdIV12jonction1squenceNCB   GCGATCGACACGTCACTCTCCCCCCTTAGTGATACCTAAAAATCGCTTCTGCGCAGGCAA
PnapiHdIV12jonction1sample5      ------------------------------------------------------------
PnapiHdIV12jonction1sample21     ------------------------------------------------------------

Prim.cons.                       GCGATCGACACGTCACTCTCCCCCCTTAGTGATACCTAAAAATCGCTTCTGCGCAGGCAA


                                        850       860       870       880       890       900
                                          |         |         |         |         |         |
PnapiHdIV12jonction1squenceNCB   TGACACTTGTTCATGATGTATGCCGGTATCTGTACTTTAGAATATTTTCGGGAAATTATG
PnapiHdIV12jonction1sample5      ------------------------------------------------------------
PnapiHdIV12jonction1sample21     ------------------------------------------------------------

Prim.cons.                       TGACACTTGTTCATGATGTATGCCGGTATCTGTACTTTAGAATATTTTCGGGAAATTATG

                                        910       920       930       940       950       960
                                          |         |         |         |         |         |
PnapiHdIV12jonction1squenceNCB   AATATTTTATTTGAATTTTAAAGCCATACAAAGCGCACTAATTAAAAATTATGGAATTCA
PnapiHdIV12jonction1sample5      ------------------------------------------------------------
PnapiHdIV12jonction1sample21     ------------------------------------------------------------

Prim.cons.                       AATATTTTATTTGAATTTTAAAGCCATACAAAGCGCACTAATTAAAAATTATGGAATTCA

                                        970       980       990      1000      1010      1020
                                          |         |         |         |         |         |
PnapiHdIV12jonction1squenceNCB   CCATTATAGCCATAAAAAGAACGACAAAAAATAGTCTAACTTAATAATAAATTCTACAAA
PnapiHdIV12jonction1sample5      ------------------------------------------------------------
PnapiHdIV12jonction1sample21     ------------------------------------------------------------

Prim.cons.                       CCATTATAGCCATAAAAAGAACGACAAAAAATAGTCTAACTTAATAATAAATTCTACAAA

                                       1030      1040      1050      1060      1070      1080
                                          |         |         |         |         |         |
PnapiHdIV12jonction1squenceNCB   CGATATCACATATAAAAGATAATTTTAGCCCTCCACCCTTAAAGTCTAGACTCTAGGCAA
PnapiHdIV12jonction1sample5      ------------------------------------------------------------
PnapiHdIV12jonction1sample21     ------------------------------------------------------------

Prim.cons.                       CGATATCACATATAAAAGATAATTTTAGCCCTCCACCCTTAAAGTCTAGACTCTAGGCAA

                                       1090      1100      1110      1120      1130      1140
                                          |         |         |         |         |         |
PnapiHdIV12jonction1squenceNCB   TTGCTCTATATATTTCGCTACCTCGATTCACATAAAATTGTAGCAATAACCATAGAAGAC
PnapiHdIV12jonction1sample5      -TGCTCTATATATTTCGCTACCTCGATTCACATAAAATTGTAGCAATAACCATAGAAGAC
PnapiHdIV12jonction1sample21     -----------ATTTCGCTACCTCGATTCACATAAAATTGTAGCAATAACCATAGAAGAC
                                            *************************************************
Prim.cons.                       TTGCTCTATATATTTCGCTACCTCGATTCACATAAAATTGTAGCAATAACCATAGAAGAC

                                       1150      1160      1170      1180      1190      1200
                                          |         |         |         |         |         |
PnapiHdIV12jonction1squenceNCB   CTTTCCATTATATCCCTTAACTAAACTACAGGGTGGGTTTTTGTTTATACAGCAAAAGTA
PnapiHdIV12jonction1sample5      CTTTCCATTATATCCCTTAACT-----ACAGGGTGGGTTTTTGTTTATACAGCAAAATTA
PnapiHdIV12jonction1sample21     CTTTCCATTATATCCCTTAACT-----ACAGGGTGGGTTTTTGATTATACAGCAAAAGTA
                                 **********************     **************** ************* **
Prim.cons.                       CTTTCCATTATATCCCTTAACTAAACTACAGGGTGGGTTTTTGTTTATACAGCAAAAGTA

                                       1210      1220      1230      1240      1250      1260
                                          |         |         |         |         |         |
PnapiHdIV12jonction1squenceNCB   TAAATGTGATAATATTTTACANACGTCCGAAATTTGTGTAAATGGGGAATGTAAAACAAA
PnapiHdIV12jonction1sample5      TAAATGTGATAATATTTTACACACGTCCGAAATTTGTGTAAATGGGGAATGTAAAACAAA
PnapiHdIV12jonction1sample21     TAAATGTCATAATATTTTACACACGTCCGAAATTTGTGTAAATGGGGAATGTAAAGCAAA
                                 ******* ************* ********************************* ****
Prim.cons.                       TAAATGTGATAATATTTTACACACGTCCGAAATTTGTGTAAATGGGGAATGTAAAACAAA

                                       1270      1280      1290      1300      1310      1320
                                          |         |         |         |         |         |
PnapiHdIV12jonction1squenceNCB   GCGATACAGTCATGGTCATAAGACCGTACGGGGCAAAAATTCAAAATGGTGGACCGGGGN
PnapiHdIV12jonction1sample5      GAGATACAGTCATGGTCATAAGACCGTACGGGGCAAAAATTCAAAATGGTGGACCGGGGA
PnapiHdIV12jonction1sample21     GCGATACAGTCATGGTCATAAGACCGTACGGGGAAAAAATTCAAAAGGGT-----GGGGT
                                 * ******************************* ************ ***     ****
Prim.cons.                       GCGATACAGTCATGGTCATAAGACCGTACGGGGCAAAAATTCAAAATGGTGGACCGGGG3

                                       1330      1340      1350      1360      1370      1380
                                          |         |         |         |         |         |
PnapiHdIV12jonction1squenceNCB   AAGGGGAGGGTGATACCAATGTATGCAGCGGAGAAGTGACTCTGATGCATAGTGGACTCG
PnapiHdIV12jonction1sample5      AAGGGGAGGGTGATACCAATGTATGCAGCGGAGAAGTGACTCTGATGCATAGTGGACTCG
PnapiHdIV12jonction1sample21     AAGGGGAGGGTGATACCAATGTATGCAGCGGAGAAGTGACTCTGATGCATAGTGGACTCG
                                 ************************************************************
Prim.cons.                       AAGGGGAGGGTGATACCAATGTATGCAGCGGAGAAGTGACTCTGATGCATAGTGGACTCG


                                       1390      1400      1410      1420      1430      1440
                                          |         |         |         |         |         |
PnapiHdIV12jonction1squenceNCB   AGAAAAGTATGATTGTTTAAGAAGTTCTAGGTTTTGCTATGTGAAAATGAGANCTAGATT
PnapiHdIV12jonction1sample5      GGAGAAGTATGATTGTTTAAGAAGTTCTAGGTTTTGCTATGTGAAAATGAGACCTAGATT
PnapiHdIV12jonction1sample21     GGAAAAGTATGATTGTTTAAGAAGTTCTAGGTTTTGCTATGTGAAAATGAGTCCTAGATT
                                  ** ***********************************************  *******
Prim.cons.                       GGAAAAGTATGATTGTTTAAGAAGTTCTAGGTTTTGCTATGTGAAAATGAGACCTAGATT

                                       1450      1460      1470      1480      1490      1500
                                          |         |         |         |         |         |
PnapiHdIV12jonction1squenceNCB   CGAGTTCCCCTAACCTTAGGTCTCTGTGAATGCGAGCGAAAAGCAAAAACACTTAACTGC
PnapiHdIV12jonction1sample5      CGAGTTCCCCTAACCTTAGG----------------------------------------
PnapiHdIV12jonction1sample21     CGAGTTCCCCTAACCTTAGG----------------------------------------
                                 ********************
Prim.cons.                       CGAGTTCCCCTAACCTTAGGTCTCTGTGAATGCGAGCGAAAAGCAAAAACACTTAACTGC

                                       1510      1520      1530      1540      1550      1560
                                          |         |         |         |         |         |
PnapiHdIV12jonction1squenceNCB   TAACCTATGAGTGCACATTGGTTCTAGGTAGCAAAATTGATATCGGCAAATTGCCACGTA
PnapiHdIV12jonction1sample5      ------------------------------------------------------------
PnapiHdIV12jonction1sample21     ------------------------------------------------------------

Prim.cons.                       TAACCTATGAGTGCACATTGGTTCTAGGTAGCAAAATTGATATCGGCAAATTGCCACGTA

                                       1570      1580      1590      1600      1610      1620
                                          |         |         |         |         |         |
PnapiHdIV12jonction1squenceNCB   ATGTCANGACATTAGAGCGACCTACCGTGACGGCGATAACGCTGTTATCGCTGAATTTAA
PnapiHdIV12jonction1sample5      ------------------------------------------------------------
PnapiHdIV12jonction1sample21     ------------------------------------------------------------

Prim.cons.                       ATGTCANGACATTAGAGCGACCTACCGTGACGGCGATAACGCTGTTATCGCTGAATTTAA

                                       1630      1640      1650      1660      1670      1680
                                          |         |         |         |         |         |
PnapiHdIV12jonction1squenceNCB   ATGGTATCAATGTGAATTTTCTGTGATCCAATTAAAAAAGTTATCNGGCTTAACTAACCA
PnapiHdIV12jonction1sample5      ------------------------------------------------------------
PnapiHdIV12jonction1sample21     ------------------------------------------------------------

Prim.cons.                       ATGGTATCAATGTGAATTTTCTGTGATCCAATTAAAAAAGTTATCNGGCTTAACTAACCA

                                       1690      1700      1710      1720      1730      1740
                                          |         |         |         |         |         |
PnapiHdIV12jonction1squenceNCB   ATCTTATGGGGTTCATTATGATATTAATCGTTCGGGGAAATAAAACAATACNCCATAGTT
PnapiHdIV12jonction1sample5      ------------------------------------------------------------
PnapiHdIV12jonction1sample21     ------------------------------------------------------------

Prim.cons.                       ATCTTATGGGGTTCATTATGATATTAATCGTTCGGGGAAATAAAACAATACNCCATAGTT

                                       1750      1760      1770      1780      1790      1800
                                          |         |         |         |         |         |
PnapiHdIV12jonction1squenceNCB   CACGACGTCATACCTACGGTGAAACACAGGCACACCTATGAAGATGAGTGTTCATTATCC
PnapiHdIV12jonction1sample5      ------------------------------------------------------------
PnapiHdIV12jonction1sample21     ------------------------------------------------------------

Prim.cons.                       CACGACGTCATACCTACGGTGAAACACAGGCACACCTATGAAGATGAGTGTTCATTATCC

                                       1810      1820      1830      1840      1850      1860
                                          |         |         |         |         |         |
PnapiHdIV12jonction1squenceNCB   AACGATTGTGTGGATGGATGAATCAATGTAAACCTAAGTTTCTCAAGTTCCGTGGTGAAC
PnapiHdIV12jonction1sample5      ------------------------------------------------------------
PnapiHdIV12jonction1sample21     ------------------------------------------------------------

Prim.cons.                       AACGATTGTGTGGATGGATGAATCAATGTAAACCTAAGTTTCTCAAGTTCCGTGGTGAAC

                                       1870      1880      1890      1900      1910      1920
                                          |         |         |         |         |         |
PnapiHdIV12jonction1squenceNCB   TTCTTTGCGGACATGAGTGCACGCTACAACGCTGTGGTGGAATGCTCGGTGTAACGGACC
PnapiHdIV12jonction1sample5      ------------------------------------------------------------
PnapiHdIV12jonction1sample21     ------------------------------------------------------------

Prim.cons.                       TTCTTTGCGGACATGAGTGCACGCTACAACGCTGTGGTGGAATGCTCGGTGTAACGGACC


                                       1930      1940      1950      1960      1970      1980
                                          |         |         |         |         |         |
PnapiHdIV12jonction1squenceNCB   CTTCACTGAATATAACATATGTAAGACTGCTGGTCACGTAGCAAGACTCCCTGCATGCTC
PnapiHdIV12jonction1sample5      ------------------------------------------------------------
PnapiHdIV12jonction1sample21     ------------------------------------------------------------

Prim.cons.                       CTTCACTGAATATAACATATGTAAGACTGCTGGTCACGTAGCAAGACTCCCTGCATGCTC

                                       1990      2000      2010      2020      2030      2040
                                          |         |         |         |         |         |
PnapiHdIV12jonction1squenceNCB   CTCATGCCAGGCAGCCTGCAGTTACATTTTAAACTCTTTAGTTGCATTTTTGTACCATAG
PnapiHdIV12jonction1sample5      ------------------------------------------------------------
PnapiHdIV12jonction1sample21     ------------------------------------------------------------

Prim.cons.                       CTCATGCCAGGCAGCCTGCAGTTACATTTTAAACTCTTTAGTTGCATTTTTGTACCATAG

                                       2050      2060      2070      2080      2090      2100
                                          |         |         |         |         |         |
PnapiHdIV12jonction1squenceNCB   CGAGTATCTCCTGAAGCTTACCTGTATTTTTTTCCTATANGGAGCATGATTGTGCAAGCT
PnapiHdIV12jonction1sample5      ------------------------------------------------------------
PnapiHdIV12jonction1sample21     ------------------------------------------------------------

Prim.cons.                       CGAGTATCTCCTGAAGCTTACCTGTATTTTTTTCCTATANGGAGCATGATTGTGCAAGCT

                                       2110      2120      2130      2140      2150      2160
                                          |         |         |         |         |         |
PnapiHdIV12jonction1squenceNCB   AATCGGAAAGTTAACACAGTCTATGACATCGGTGCCATCTTCAACATTTTTCCCTTCTTC
PnapiHdIV12jonction1sample5      ------------------------------------------------------------
PnapiHdIV12jonction1sample21     ------------------------------------------------------------

Prim.cons.                       AATCGGAAAGTTAACACAGTCTATGACATCGGTGCCATCTTCAACATTTTTCCCTTCTTC

                                       2170      2180      2190      2200      2210      2220
                                          |         |         |         |         |         |
PnapiHdIV12jonction1squenceNCB   CGTTGAAAATAATTGGGTAATGCCGATATGTATTTTCCGTTAACATCATCAATATAGCCC
PnapiHdIV12jonction1sample5      ------------------------------------------------------------
PnapiHdIV12jonction1sample21     ------------------------------------------------------------

Prim.cons.                       CGTTGAAAATAATTGGGTAATGCCGATATGTATTTTCCGTTAACATCATCAATATAGCCC

                                       2230      2240      2250      2260      2270      2280
                                          |         |         |         |         |         |
PnapiHdIV12jonction1squenceNCB   AATTTGTATCGCAAAATACTTTTCTCTACCTAAATTCGATATCATTAGCGAGTTTTGTTA
PnapiHdIV12jonction1sample5      ------------------------------------------------------------
PnapiHdIV12jonction1sample21     ------------------------------------------------------------

Prim.cons.                       AATTTGTATCGCAAAATACTTTTCTCTACCTAAATTCGATATCATTAGCGAGTTTTGTTA

                                       2290      2300      2310      2320      2330      2340
                                          |         |         |         |         |         |
PnapiHdIV12jonction1squenceNCB   AAAACAAAAAAAATCCTGATAACTATAAAATGCTAACCGGAACACATTAGGTAGGTTTCG
PnapiHdIV12jonction1sample5      ------------------------------------------------------------
PnapiHdIV12jonction1sample21     ------------------------------------------------------------

Prim.cons.                       AAAACAAAAAAAATCCTGATAACTATAAAATGCTAACCGGAACACATTAGGTAGGTTTCG

                                       2350      2360      2370      2380      2390      2400
                                          |         |         |         |         |         |
PnapiHdIV12jonction1squenceNCB   AATTAGTATAAATTGAGGGAAGGACCTATACATTNAATCATTAGCATCATTACCCCCGAC
PnapiHdIV12jonction1sample5      ------------------------------------------------------------
PnapiHdIV12jonction1sample21     ------------------------------------------------------------

Prim.cons.                       AATTAGTATAAATTGAGGGAAGGACCTATACATTNAATCATTAGCATCATTACCCCCGAC

                                       2410      2420      2430      2440      2450      2460
                                          |         |         |         |         |         |
PnapiHdIV12jonction1squenceNCB   ATAATCACCAGGAGCTATACAGTCAATNAACACTAATACCCTCGACATAGTCACCATGTG
PnapiHdIV12jonction1sample5      ------------------------------------------------------------
PnapiHdIV12jonction1sample21     ------------------------------------------------------------

Prim.cons.                       ATAATCACCAGGAGCTATACAGTCAATNAACACTAATACCCTCGACATAGTCACCATGTG


                                       2470      2480      2490      2500      2510      2520
                                          |         |         |         |         |         |
PnapiHdIV12jonction1squenceNCB   GGGGAACAATAGTGGAAACTCCGGAGAAGCCGCTCGTCGGTACGCCGAGGAAGAGAAGTG
PnapiHdIV12jonction1sample5      ------------------------------------------------------------
PnapiHdIV12jonction1sample21     ------------------------------------------------------------

Prim.cons.                       GGGGAACAATAGTGGAAACTCCGGAGAAGCCGCTCGTCGGTACGCCGAGGAAGAGAAGTG

                                       2530      2540      2550      2560      2570      2580
                                          |         |         |         |         |         |
PnapiHdIV12jonction1squenceNCB   CAGACAAGACAAGGAGCGTGATGAGAGGAAGCGTCAGCAGGAAGAGCAAGAGCGAATTGC
PnapiHdIV12jonction1sample5      ------------------------------------------------------------
PnapiHdIV12jonction1sample21     ------------------------------------------------------------

Prim.cons.                       CAGACAAGACAAGGAGCGTGATGAGAGGAAGCGTCAGCAGGAAGAGCAAGAGCGAATTGC

                                       2590      2600
                                          |         |
PnapiHdIV12jonction1squenceNCB   GGAGCGCGTGAGGCAGTATG
PnapiHdIV12jonction1sample5      --------------------
PnapiHdIV12jonction1sample21     --------------------

Prim.cons.                       GGAGCGCGTGAGGCAGTATG
```

>P. napi HdIV12 jonction2 – NCBI accession number (start-end coordinates): DWAF01023688.1 (637-2062)

AAAGTTACTTCAGTTTTGTTCATGGACAGGAGAAACCGTCGATATTCACTATCAAAATGTAAATATCGGTTCACGCTGAAGGGGGCCCCACACGGTAGACGAAAACCACGTTTCGCAAAGTCATTTACTGTGAAACTTTGTGACTTCTTTTCTTCTTGTAACGCCATTGTGACTCCATGAAGTTGTGCGGCCTTTTGTTTATGTTTCTTGCTGAAGTGTTGAACGTTAATTGATGAGCAGTCGTTTCGACTGCTTATTTTATAGGTGTATTGTTGCCGGACAAAAGTCAACACTGATAACCTAGAAGTTATTTAAAAAAAAATCGGAATTCCCTCAAAAATATAACTTATCAGCAAGTGGCATGCTTCGTAGTAAACAACTGATTAGGTACATGAATGATTCTAGGTATAAAAAACAATGCTGAGACCTCGTCAGCTGAACTTTCGGTGAACTACGGTGAACTTTACTCAACCGAATACTGATAACGCAGAAGACGTAATAGACATAAACAAAATTCATCAGATTTTCCTGACTAATATAACCGTTTGGTTATCGGCCGACTTTGTGATAAGCAAGCGACTTGGGAAATTGGCTGTTGACAAAACAACTTCAATTCTAGCAGTAAGGGTCTCGTGAGTACCCGCTATGAAGGCATGCTTAGATTATACAAAAGTGGCGAAGCGATGTATGGCCTACGAGATGCGGGTGACCGTTCCATGTCAACAAAGTAGATAAGTTTACAAGAAGTATGTTTGTTCTAGGCGTTCCAAGTGTCGTTCTGCCAGAACGTGAGCTGTAGGTTTCTGATAAGTTGGCGCTACCTTAAATGCTGATTCTGTGGCCCCAGCGATTCATGTTACCTGCTGCTTTGACCCATATTTTTAGCACCTTAGAGCATGCACCTACAGAAAGGGTCGAAATTTTCGAAAATGCATCTGACCTCTATGCACTCTAGTATCTTCATCACTGAATGCGAAAATATCAACCTTGCCTAGCCTAGATGCTTAATTTTGAAGTTATAGACCGCACGGTAACGTGGCACCTACTGTACATTGTTGCGTTACGTTGAACTGTCATTGGTCAAGAGACGTATTGCATTTCTGTGATTCCTCAGCTGTCGACCAATTATAATCAAGTTGCGTTTCGTTTTTTGTTTTTTTTTATGTAATAGAAGGCGATGGATATTAAGGGATACCGCAGCCTGAGGACACTCATATTGCGAGAAAGCTCGCAAGTGCGTTGCCGGCCTTTTAATAATTGGTACGCTTTTCTTGAAGGACCCTAAGTCTAATTGGTACTGACCTTGTCGGAAATACTTCAGTAGGCAAAAAACTACCTTAAGTTTTTGGAACGAAGTTCCTAATCGCGCGCTGTGAAAGGGGGCTGGACGGAAAAAATTCTTACGAAAAGTTGTCACGACACTTTTTTGCTATTTGCTATTGTA

>P. napi HdIV12 jonction2 sample5

TTCTGATAAGTTGGCGCTACCTTGGATGCTGATTCTGTGGCCCCAGCGATTCATGTTACCTGCTGCTTTGACCCATATTTTTAGCACCTTAGAGCATGCACCTACAGAACAGGTCGAAATTTTCGAAAATGCATCTGACCTCTATGCACTGTAGTATCTTCATCACTGAATCCGAAAATATCAACCTTGCCTAGCCTAGATGCTTAATTTTGAAGTTATAGACCGCACGGTAACGTGGCACCTACTGTACATTGTTGCGTTACGTTGAACTGTCATTGGTCAAGAGACGTATTGCAA

>P. napi HdIV12 jonction2 sample25

CTGTAGGTTTCTGATAAGTTGTCGCTACCTTGGATGCTGATTCTGCGGCCCCAGCGATTCATGTTACCTGCTGCTTTGACCCATATTTTTAGCACCTTAGAGCATGCACCTACAGGACAGGTCGAAATTTTCGAAAATGCATCTGACCTCTATGCACTCTAGTATCTTCATCACTGAATGCGAAAATATCAACCTTGCCTAGCCTAGATGCTTAATTTTGAAGTTATAGACCGCACGGTAACGTGGCACCTACTGTACATTGTTGCGTTACGTTGAACTGTCATTGGTCAAGAGACGTATTGCAA

Multiple alignment

```
                                         10        20        30        40        50        60
                                          |         |         |         |         |         |
PnapiHdIV12jonction2sample5      ------------------------------------------------------------
PnapiHdIV12jonction2sample25     ------------------------------------------------------------
PnapiHdIV12jonction2squenceNCB   AAAGTTACTTCAGTTTTGTTCATGGACAGGAGAAACCGTCGATATTCACTATCAAAATGT

Prim.cons.                       AAAGTTACTTCAGTTTTGTTCATGGACAGGAGAAACCGTCGATATTCACTATCAAAATGT

                                         70        80        90       100       110       120
                                          |         |         |         |         |         |
PnapiHdIV12jonction2sample5      ------------------------------------------------------------
PnapiHdIV12jonction2sample25     ------------------------------------------------------------
PnapiHdIV12jonction2squenceNCB   AAATATCGGTTCACGCTGAAGGGGGCCCCACACGGTAGACGAAAACCACGTTTCGCAAAG

Prim.cons.                       AAATATCGGTTCACGCTGAAGGGGGCCCCACACGGTAGACGAAAACCACGTTTCGCAAAG

                                        130       140       150       160       170       180
                                          |         |         |         |         |         |
PnapiHdIV12jonction2sample5      ------------------------------------------------------------
PnapiHdIV12jonction2sample25     ------------------------------------------------------------
PnapiHdIV12jonction2squenceNCB   TCATTTACTGTGAAACTTTGTGACTTCTTTTCTTCTTGTAACGCCATTGTGACTCCATGA

Prim.cons.                       TCATTTACTGTGAAACTTTGTGACTTCTTTTCTTCTTGTAACGCCATTGTGACTCCATGA

                                        190       200       210       220       230       240
                                          |         |         |         |         |         |
PnapiHdIV12jonction2sample5      ------------------------------------------------------------
PnapiHdIV12jonction2sample25     ------------------------------------------------------------
PnapiHdIV12jonction2squenceNCB   AGTTGTGCGGCCTTTTGTTTATGTTTCTTGCTGAAGTGTTGAACGTTAATTGATGAGCAG

Prim.cons.                       AGTTGTGCGGCCTTTTGTTTATGTTTCTTGCTGAAGTGTTGAACGTTAATTGATGAGCAG

                                        250       260       270       280       290       300
                                          |         |         |         |         |         |
PnapiHdIV12jonction2sample5      ------------------------------------------------------------
PnapiHdIV12jonction2sample25     ------------------------------------------------------------
PnapiHdIV12jonction2squenceNCB   TCGTTTCGACTGCTTATTTTATAGGTGTATTGTTGCCGGACAAAAGTCAACACTGATAAC

Prim.cons.                       TCGTTTCGACTGCTTATTTTATAGGTGTATTGTTGCCGGACAAAAGTCAACACTGATAAC

                                        310       320       330       340       350       360
                                          |         |         |         |         |         |
PnapiHdIV12jonction2sample5      ------------------------------------------------------------
PnapiHdIV12jonction2sample25     ------------------------------------------------------------
PnapiHdIV12jonction2squenceNCB   CTAGAAGTTATTTAAAAAAAAATCGGAATTCCCTCAAAAATATAACTTATCAGCAAGTGG

Prim.cons.                       CTAGAAGTTATTTAAAAAAAAATCGGAATTCCCTCAAAAATATAACTTATCAGCAAGTGG

                                        370       380       390       400       410       420
                                          |         |         |         |         |         |
PnapiHdIV12jonction2sample5      ------------------------------------------------------------
PnapiHdIV12jonction2sample25     ------------------------------------------------------------
PnapiHdIV12jonction2squenceNCB   CATGCTTCGTAGTAAACAACTGATTAGGTACATGAATGATTCTAGGTATAAAAAACAATG

Prim.cons.                       CATGCTTCGTAGTAAACAACTGATTAGGTACATGAATGATTCTAGGTATAAAAAACAATG

                                        430       440       450       460       470       480
                                          |         |         |         |         |         |
PnapiHdIV12jonction2sample5      ------------------------------------------------------------
PnapiHdIV12jonction2sample25     ------------------------------------------------------------
PnapiHdIV12jonction2squenceNCB   CTGAGACCTCGTCAGCTGAACTTTCGGTGAACTACGGTGAACTTTACTCAACCGAATACT

Prim.cons.                       CTGAGACCTCGTCAGCTGAACTTTCGGTGAACTACGGTGAACTTTACTCAACCGAATACT

                                        490       500       510       520       530       540
                                          |         |         |         |         |         |
PnapiHdIV12jonction2sample5      ------------------------------------------------------------
PnapiHdIV12jonction2sample25     ------------------------------------------------------------
PnapiHdIV12jonction2squenceNCB   GATAACGCAGAAGACGTAATAGACATAAACAAAATTCATCAGATTTTCCTGACTAATATA

Prim.cons.                       GATAACGCAGAAGACGTAATAGACATAAACAAAATTCATCAGATTTTCCTGACTAATATA

                                        550       560       570       580       590       600
                                          |         |         |         |         |         |
PnapiHdIV12jonction2sample5      ------------------------------------------------------------
PnapiHdIV12jonction2sample25     ------------------------------------------------------------
PnapiHdIV12jonction2squenceNCB   ACCGTTTGGTTATCGGCCGACTTTGTGATAAGCAAGCGACTTGGGAAATTGGCTGTTGAC

Prim.cons.                       ACCGTTTGGTTATCGGCCGACTTTGTGATAAGCAAGCGACTTGGGAAATTGGCTGTTGAC

                                        610       620       630       640       650       660
                                          |         |         |         |         |         |
PnapiHdIV12jonction2sample5      ------------------------------------------------------------
PnapiHdIV12jonction2sample25     ------------------------------------------------------------
PnapiHdIV12jonction2squenceNCB   AAAACAACTTCAATTCTAGCAGTAAGGGTCTCGTGAGTACCCGCTATGAAGGCATGCTTA

Prim.cons.                       AAAACAACTTCAATTCTAGCAGTAAGGGTCTCGTGAGTACCCGCTATGAAGGCATGCTTA

                                        670       680       690       700       710       720
                                          |         |         |         |         |         |
PnapiHdIV12jonction2sample5      ------------------------------------------------------------
PnapiHdIV12jonction2sample25     ------------------------------------------------------------
PnapiHdIV12jonction2squenceNCB   GATTATACAAAAGTGGCGAAGCGATGTATGGCCTACGAGATGCGGGTGACCGTTCCATGT

Prim.cons.                       GATTATACAAAAGTGGCGAAGCGATGTATGGCCTACGAGATGCGGGTGACCGTTCCATGT

                                        730       740       750       760       770       780
                                          |         |         |         |         |         |
PnapiHdIV12jonction2sample5      ------------------------------------------------------------
PnapiHdIV12jonction2sample25     ------------------------------------------------------------
PnapiHdIV12jonction2squenceNCB   CAACAAAGTAGATAAGTTTACAAGAAGTATGTTTGTTCTAGGCGTTCCAAGTGTCGTTCT

Prim.cons.                       CAACAAAGTAGATAAGTTTACAAGAAGTATGTTTGTTCTAGGCGTTCCAAGTGTCGTTCT

                                        790       800       810       820       830       840
                                          |         |         |         |         |         |
PnapiHdIV12jonction2sample5      ---------------------TTCTGATAAGTTGGCGCTACCTTGGATGCTGATTCTGTG
PnapiHdIV12jonction2sample25     -------------CTGTAGGTTTCTGATAAGTTGTCGCTACCTTGGATGCTGATTCTGCG
PnapiHdIV12jonction2squenceNCB   GCCAGAACGTGAGCTGTAGGTTTCTGATAAGTTGGCGCTACCTTAAATGCTGATTCTGTG
                                                      ************* *********  ************ *
Prim.cons.                       GCCAGAACGTGAGCTGTAGGTTTCTGATAAGTTGGCGCTACCTTGGATGCTGATTCTGTG

                                        850       860       870       880       890       900
                                          |         |         |         |         |         |
PnapiHdIV12jonction2sample5      GCCCCAGCGATTCATGTTACCTGCTGCTTTGACCCATATTTTTAGCACCTTAGAGCATGC
PnapiHdIV12jonction2sample25     GCCCCAGCGATTCATGTTACCTGCTGCTTTGACCCATATTTTTAGCACCTTAGAGCATGC
PnapiHdIV12jonction2squenceNCB   GCCCCAGCGATTCATGTTACCTGCTGCTTTGACCCATATTTTTAGCACCTTAGAGCATGC
                                 ************************************************************
Prim.cons.                       GCCCCAGCGATTCATGTTACCTGCTGCTTTGACCCATATTTTTAGCACCTTAGAGCATGC


                                        910       920       930       940       950       960
                                          |         |         |         |         |         |
PnapiHdIV12jonction2sample5      ACCTACAGAACAGGTCGAAATTTTCGAAAATGCATCTGACCTCTATGCACTGTAGTATCT
PnapiHdIV12jonction2sample25     ACCTACAGGACAGGTCGAAATTTTCGAAAATGCATCTGACCTCTATGCACTCTAGTATCT
PnapiHdIV12jonction2squenceNCB   ACCTACAGAAAGGGTCGAAATTTTCGAAAATGCATCTGACCTCTATGCACTCTAGTATCT
                                 ******** *  *************************************** ********
Prim.cons.                       ACCTACAGAACAGGTCGAAATTTTCGAAAATGCATCTGACCTCTATGCACTCTAGTATCT

                                        970       980       990      1000      1010      1020
                                          |         |         |         |         |         |
PnapiHdIV12jonction2sample5      TCATCACTGAATCCGAAAATATCAACCTTGCCTAGCCTAGATGCTTAATTTTGAAGTTAT
PnapiHdIV12jonction2sample25     TCATCACTGAATGCGAAAATATCAACCTTGCCTAGCCTAGATGCTTAATTTTGAAGTTAT
PnapiHdIV12jonction2squenceNCB   TCATCACTGAATGCGAAAATATCAACCTTGCCTAGCCTAGATGCTTAATTTTGAAGTTAT
                                 ************ ***********************************************
Prim.cons.                       TCATCACTGAATGCGAAAATATCAACCTTGCCTAGCCTAGATGCTTAATTTTGAAGTTAT

                                       1030      1040      1050      1060      1070      1080
                                          |         |         |         |         |         |
PnapiHdIV12jonction2sample5      AGACCGCACGGTAACGTGGCACCTACTGTACATTGTTGCGTTACGTTGAACTGTCATTGG
PnapiHdIV12jonction2sample25     AGACCGCACGGTAACGTGGCACCTACTGTACATTGTTGCGTTACGTTGAACTGTCATTGG
PnapiHdIV12jonction2squenceNCB   AGACCGCACGGTAACGTGGCACCTACTGTACATTGTTGCGTTACGTTGAACTGTCATTGG
                                 ************************************************************
Prim.cons.                       AGACCGCACGGTAACGTGGCACCTACTGTACATTGTTGCGTTACGTTGAACTGTCATTGG

                                       1090      1100      1110      1120      1130      1140
                                          |         |         |         |         |         |
PnapiHdIV12jonction2sample5      TCAAGAGACGTATTGCAA------------------------------------------
PnapiHdIV12jonction2sample25     TCAAGAGACGTATTGCAA------------------------------------------
PnapiHdIV12jonction2squenceNCB   TCAAGAGACGTATTGCATTTCTGTGATTCCTCAGCTGTCGACCAATTATAATCAAGTTGC
                                 *****************
Prim.cons.                       TCAAGAGACGTATTGCAATTCTGTGATTCCTCAGCTGTCGACCAATTATAATCAAGTTGC

                                       1150      1160      1170      1180      1190      1200
                                          |         |         |         |         |         |
PnapiHdIV12jonction2sample5      ------------------------------------------------------------
PnapiHdIV12jonction2sample25     ------------------------------------------------------------
PnapiHdIV12jonction2squenceNCB   GTTTCGTTTTTTGTTTTTTTTTATGTAATAGAAGGCGATGGATATTAAGGGATACCGCAG

Prim.cons.                       GTTTCGTTTTTTGTTTTTTTTTATGTAATAGAAGGCGATGGATATTAAGGGATACCGCAG

                                       1210      1220      1230      1240      1250      1260
                                          |         |         |         |         |         |
PnapiHdIV12jonction2sample5      ------------------------------------------------------------
PnapiHdIV12jonction2sample25     ------------------------------------------------------------
PnapiHdIV12jonction2squenceNCB   CCTGAGGACACTCATATTGCGAGAAAGCTCGCAAGTGCGTTGCCGGCCTTTTAATAATTG

Prim.cons.                       CCTGAGGACACTCATATTGCGAGAAAGCTCGCAAGTGCGTTGCCGGCCTTTTAATAATTG

                                       1270      1280      1290      1300      1310      1320
                                          |         |         |         |         |         |
PnapiHdIV12jonction2sample5      ------------------------------------------------------------
PnapiHdIV12jonction2sample25     ------------------------------------------------------------
PnapiHdIV12jonction2squenceNCB   GTACGCTTTTCTTGAAGGACCCTAAGTCTAATTGGTACTGACCTTGTCGGAAATACTTCA

Prim.cons.                       GTACGCTTTTCTTGAAGGACCCTAAGTCTAATTGGTACTGACCTTGTCGGAAATACTTCA

                                       1330      1340      1350      1360      1370      1380
                                          |         |         |         |         |         |
PnapiHdIV12jonction2sample5      ------------------------------------------------------------
PnapiHdIV12jonction2sample25     ------------------------------------------------------------
PnapiHdIV12jonction2squenceNCB   GTAGGCAAAAAACTACCTTAAGTTTTTGGAACGAAGTTCCTAATCGCGCGCTGTGAAAGG

Prim.cons.                       GTAGGCAAAAAACTACCTTAAGTTTTTGGAACGAAGTTCCTAATCGCGCGCTGTGAAAGG

                                       1390      1400      1410      1420      1430      1440
                                          |         |         |         |         |         |
PnapiHdIV12jonction2sample5      ------------------------------------------------------------
PnapiHdIV12jonction2sample25     ------------------------------------------------------------
PnapiHdIV12jonction2squenceNCB   GGGCTGGACGGAAAAAATTCTTACGAAAAGTTGTCACGACACTTTTTTGCTATTTGCTAT

Prim.cons.                       GGGCTGGACGGAAAAAATTCTTACGAAAAGTTGTCACGACACTTTTTTGCTATTTGCTAT


PnapiHdIV12jonction2sample5      ----
PnapiHdIV12jonction2sample25     ----
PnapiHdIV12jonction2squenceNCB   TGTA

Prim.cons.                       TGTA
```

>P. rapae CcBV1 jonction2 – NCBI accession number (start-end coordinates): LWME01000268.1 (268186-269456)

CAACGAAGATAAAACTTAATCTATCTGACCAGACCTCGGAAAAACTAACTCGCAAAATTTAAGGAAAAAATATGTTGATAAGATAACAAACTGGGTTTTAATTGGCACGCAATAATATTGTGATAAAATAATTTAAAAACAAACGAGTTTTTTCGCGTTTAGAATTCTAACTTTTGACAACGATTAGGACACTGGTTACATCTTTAAAGGTTATCTTCGCGCAACATTAGGAATAAAAGTACCTAACTAGGTAATTTAGTTACGTTTTTTTTTAATATTCACACGTAGAAAGCTTAGCGAATCTTTGCACGTTTTTTTCAATTCGTTTAGAATAATTATTTATTGTCACTATGACCTGTCAACTGGTCAGTTGTTAACAGTACAGACAGTACAATCTTCAACCTTCTCATCAAACTCCATCATTCAGTTCCACGTGGACTGATCTCCAGCTTCATCAACAGGACGTTACTTAATCAATCACGAAGCCGGTGTAAAATAATTGTGTACGGTATACTGTGAAACATACATCATTAAATGTTCATTAAGAAATATACACTGTTCGCATTAACCAACAAACTAATAAAAAACACGCAGCTTTCCATAATTTTAACATCAGTTATTAATAAAGCGATTATTTCTCCGGTTTTGTGTATAACTTACAATGCCTAGTTTATGTGCATGTGTATGTTTCATATTATACAAAATACCTACTATGTAATACAATACAGAAAGCAAAAACAGATTACATAGATTAACAAATTATGTATTAAAAACAGAAAACAANCAGGCGCATTAAAAGACAAAGATACATTGAAAAAAAAAACAATAGAATAAAATAAAAACACTGAAATTTGAGATTTAAAAGCAACAAATAATACTAAGAATTTAAAACTACTACTTAAAAAAATCTGATTCCCGCTTCTCCAAGTANGTACTTCTTGAAATCTCTCTTTTTCTTAAATAATACGACAATTTTAAGATGTTTTATTAAAATTGACGTATTATTTCTTACACCCTTATTTCTTATACTTATATCTTTCACAATAAATATTATCGCTACTATATAATGCACTTAGTACTTTTAAACGCGGCCCGCGGACGTATTTNCACGCTTAAGCTTGGCCCTCAACATGTNAAAGGTTGCCCATACCTGATCTAGAAGGTATGACTTAACATCAGGCGAACCACCTGACTCTATGCCCTCAGTTATATATTTGAATGTCGTTGATCTTTTGCCATAATGTTTTCCTCTCGATGTATTTCTTCACCGTAAAACCCTC

>P. rapae CcBV1 jonction2 sample15

GGAAAAACTAACTCGCAAAATTTAAGGAAAAAATATGTTGATAAGATAACAAACTGGGTTTTAATTGGCACGCAATAATATTGTGATAAAATAATTTAAAAACAAACGAGTTTTTTCGCGTTTAGAATTCTAACTTTTGACAACGATTAGGACACTGGTTACATCTTTAAAGGTTATCTTCGCGCAACATTAGGAATAAAAGTACCTAACTAGGTAATTTAGTTACGTTTTTTTTTAATATTCACACGTAGAAAGCTTAGCGAATCTTTGCACGTTTTTTTCAATTCGTTTAGAATAATTATTTATTGTCACTATGACCTGTCAACTGGTCAGTTGTTAACAGTACAGACAGTACAATCTTCAACCTTCTCATCAAACTCCATCATTCAGTTCCACGTGGAC

>P. rapae CcBV1 jonction2 sample30

GACCAGATCCTCGGAAAAACTAACTCGCAAAATTTAAGGAAAAAATATGTTGATAAGATAACAAACTGGGTTTTAATTGGCACGCAATAATATTGTGATAAAATAATTTAAAAACAAACGAGTTTTTTCGCGTTTAGAATTCTAATTTTTGACAACGATTAGGACACTGGTTACATCTTTAAAGGTTATCTTCGCGCAACATTAGGAATAAAAGTACCTAACTAGGTAATTTAGTTACGTTTTTTTTTAATATTCACACGTAGAAAGCTTAGCGAATCTTTGCACGTTTTTTTCAATTCGTTTAGAATAATTATTTATTGTCACTATGACCTGTCAACTGGTCAGTTGTTAACAGTACAGACAGTACAATCTTCAACCTTCTCATCAAACTCCATCATTCAGTTCCACGTGGAC

Multiple alignment

```
                                         10        20        30        40        50        60
                                          |         |         |         |         |         |
PrapaeCcBV1jonction2squenceNCB   CAACGAAGATAAAACTTAATCTATCTGACCAGA-CCTCGGAAAAACTAACTCGCAAAATT
PrapaeCcBV1jonction2sample15     --------------------------------------GGAAAAACTAACTCGCAAAATT
PrapaeCcBV1jonction2sample30     --------------------------GACCAGATCCTCGGAAAAACTAACTCGCAAAATT
                                                                       **********************
Prim.cons.                       CAACGAAGATAAAACTTAATCTATCTGACCAGATCCTCGGAAAAACTAACTCGCAAAATT

                                         70        80        90       100       110       120
                                          |         |         |         |         |         |
PrapaeCcBV1jonction2squenceNCB   TAAGGAAAAAATATGTTGATAAGATAACAAACTGGGTTTTAATTGGCACGCAATAATATT
PrapaeCcBV1jonction2sample15     TAAGGAAAAAATATGTTGATAAGATAACAAACTGGGTTTTAATTGGCACGCAATAATATT
PrapaeCcBV1jonction2sample30     TAAGGAAAAAATATGTTGATAAGATAACAAACTGGGTTTTAATTGGCACGCAATAATATT
                                 ************************************************************
Prim.cons.                       TAAGGAAAAAATATGTTGATAAGATAACAAACTGGGTTTTAATTGGCACGCAATAATATT

                                        130       140       150       160       170       180
                                          |         |         |         |         |         |
PrapaeCcBV1jonction2squenceNCB   GTGATAAAATAATTTAAAAACAAACGAGTTTTTTCGCGTTTAGAATTCTAACTTTTGACA
PrapaeCcBV1jonction2sample15     GTGATAAAATAATTTAAAAACAAACGAGTTTTTTCGCGTTTAGAATTCTAACTTTTGACA
PrapaeCcBV1jonction2sample30     GTGATAAAATAATTTAAAAACAAACGAGTTTTTTCGCGTTTAGAATTCTAATTTTTGACA
                                 *************************************************** ********
Prim.cons.                       GTGATAAAATAATTTAAAAACAAACGAGTTTTTTCGCGTTTAGAATTCTAACTTTTGACA

                                        190       200       210       220       230       240
                                          |         |         |         |         |         |
PrapaeCcBV1jonction2squenceNCB   ACGATTAGGACACTGGTTACATCTTTAAAGGTTATCTTCGCGCAACATTAGGAATAAAAG
PrapaeCcBV1jonction2sample15     ACGATTAGGACACTGGTTACATCTTTAAAGGTTATCTTCGCGCAACATTAGGAATAAAAG
PrapaeCcBV1jonction2sample30     ACGATTAGGACACTGGTTACATCTTTAAAGGTTATCTTCGCGCAACATTAGGAATAAAAG
                                 ************************************************************
Prim.cons.                       ACGATTAGGACACTGGTTACATCTTTAAAGGTTATCTTCGCGCAACATTAGGAATAAAAG

                                        250       260       270       280       290       300
                                          |         |         |         |         |         |
PrapaeCcBV1jonction2squenceNCB   TACCTAACTAGGTAATTTAGTTACGTTTTTTTTTAATATTCACACGTAGAAAGCTTAGCG
PrapaeCcBV1jonction2sample15     TACCTAACTAGGTAATTTAGTTACGTTTTTTTTTAATATTCACACGTAGAAAGCTTAGCG
PrapaeCcBV1jonction2sample30     TACCTAACTAGGTAATTTAGTTACGTTTTTTTTTAATATTCACACGTAGAAAGCTTAGCG
                                 ************************************************************
Prim.cons.                       TACCTAACTAGGTAATTTAGTTACGTTTTTTTTTAATATTCACACGTAGAAAGCTTAGCG

                                        310       320       330       340       350       360
                                          |         |         |         |         |         |
PrapaeCcBV1jonction2squenceNCB   AATCTTTGCACGTTTTTTTCAATTCGTTTAGAATAATTATTTATTGTCACTATGACCTGT
PrapaeCcBV1jonction2sample15     AATCTTTGCACGTTTTTTTCAATTCGTTTAGAATAATTATTTATTGTCACTATGACCTGT
PrapaeCcBV1jonction2sample30     AATCTTTGCACGTTTTTTTCAATTCGTTTAGAATAATTATTTATTGTCACTATGACCTGT
                                 ************************************************************
Prim.cons.                       AATCTTTGCACGTTTTTTTCAATTCGTTTAGAATAATTATTTATTGTCACTATGACCTGT

                                        370       380       390       400       410       420
                                          |         |         |         |         |         |
PrapaeCcBV1jonction2squenceNCB   CAACTGGTCAGTTGTTAACAGTACAGACAGTACAATCTTCAACCTTCTCATCAAACTCCA
PrapaeCcBV1jonction2sample15     CAACTGGTCAGTTGTTAACAGTACAGACAGTACAATCTTCAACCTTCTCATCAAACTCCA
PrapaeCcBV1jonction2sample30     CAACTGGTCAGTTGTTAACAGTACAGACAGTACAATCTTCAACCTTCTCATCAAACTCCA
                                 ************************************************************
Prim.cons.                       CAACTGGTCAGTTGTTAACAGTACAGACAGTACAATCTTCAACCTTCTCATCAAACTCCA

                                        430       440       450       460       470       480
                                          |         |         |         |         |         |
PrapaeCcBV1jonction2squenceNCB   TCATTCAGTTCCACGTGGACTGATCTCCAGCTTCATCAACAGGACGTTACTTAATCAATC
PrapaeCcBV1jonction2sample15     TCATTCAGTTCCACGTGGAC----------------------------------------
PrapaeCcBV1jonction2sample30     TCATTCAGTTCCACGTGGAC----------------------------------------
                                 ********************
Prim.cons.                       TCATTCAGTTCCACGTGGACTGATCTCCAGCTTCATCAACAGGACGTTACTTAATCAATC

                                        490       500       510       520       530       540
                                          |         |         |         |         |         |
PrapaeCcBV1jonction2squenceNCB   ACGAAGCCGGTGTAAAATAATTGTGTACGGTATACTGTGAAACATACATCATTAAATGTT
PrapaeCcBV1jonction2sample15     ------------------------------------------------------------
PrapaeCcBV1jonction2sample30     ------------------------------------------------------------

Prim.cons.                       ACGAAGCCGGTGTAAAATAATTGTGTACGGTATACTGTGAAACATACATCATTAAATGTT

                                        550       560       570       580       590       600
                                          |         |         |         |         |         |
PrapaeCcBV1jonction2squenceNCB   CATTAAGAAATATACACTGTTCGCATTAACCAACAAACTAATAAAAAACACGCAGCTTTC
PrapaeCcBV1jonction2sample15     ------------------------------------------------------------
PrapaeCcBV1jonction2sample30     ------------------------------------------------------------

Prim.cons.                       CATTAAGAAATATACACTGTTCGCATTAACCAACAAACTAATAAAAAACACGCAGCTTTC

                                        610       620       630       640       650       660
                                          |         |         |         |         |         |
PrapaeCcBV1jonction2squenceNCB   CATAATTTTAACATCAGTTATTAATAAAGCGATTATTTCTCCGGTTTTGTGTATAACTTA
PrapaeCcBV1jonction2sample15     ------------------------------------------------------------
PrapaeCcBV1jonction2sample30     ------------------------------------------------------------

Prim.cons.                       CATAATTTTAACATCAGTTATTAATAAAGCGATTATTTCTCCGGTTTTGTGTATAACTTA

                                        670       680       690       700       710       720
                                          |         |         |         |         |         |
PrapaeCcBV1jonction2squenceNCB   CAATGCCTAGTTTATGTGCATGTGTATGTTTCATATTATACAAAATACCTACTATGTAAT
PrapaeCcBV1jonction2sample15     ------------------------------------------------------------
PrapaeCcBV1jonction2sample30     ------------------------------------------------------------

Prim.cons.                       CAATGCCTAGTTTATGTGCATGTGTATGTTTCATATTATACAAAATACCTACTATGTAAT

                                        730       740       750       760       770       780
                                          |         |         |         |         |         |
PrapaeCcBV1jonction2squenceNCB   ACAATACAGAAAGCAAAAACAGATTACATAGATTAACAAATTATGTATTAAAAACAGAAA
PrapaeCcBV1jonction2sample15     ------------------------------------------------------------
PrapaeCcBV1jonction2sample30     ------------------------------------------------------------

Prim.cons.                       ACAATACAGAAAGCAAAAACAGATTACATAGATTAACAAATTATGTATTAAAAACAGAAA

                                        790       800       810       820       830       840
                                          |         |         |         |         |         |
PrapaeCcBV1jonction2squenceNCB   ACAANCAGGCGCATTAAAAGACAAAGATACATTGAAAAAAAAAACAATAGAATAAAATAA
PrapaeCcBV1jonction2sample15     ------------------------------------------------------------
PrapaeCcBV1jonction2sample30     ------------------------------------------------------------

Prim.cons.                       ACAANCAGGCGCATTAAAAGACAAAGATACATTGAAAAAAAAAACAATAGAATAAAATAA

                                        850       860       870       880       890       900
                                          |         |         |         |         |         |
PrapaeCcBV1jonction2squenceNCB   AAACACTGAAATTTGAGATTTAAAAGCAACAAATAATACTAAGAATTTAAAACTACTACT
PrapaeCcBV1jonction2sample15     ------------------------------------------------------------
PrapaeCcBV1jonction2sample30     ------------------------------------------------------------

Prim.cons.                       AAACACTGAAATTTGAGATTTAAAAGCAACAAATAATACTAAGAATTTAAAACTACTACT

                                        910       920       930       940       950       960
                                          |         |         |         |         |         |
PrapaeCcBV1jonction2squenceNCB   TAAAAAAATCTGATTCCCGCTTCTCCAAGTANGTACTTCTTGAAATCTCTCTTTTTCTTA
PrapaeCcBV1jonction2sample15     ------------------------------------------------------------
PrapaeCcBV1jonction2sample30     ------------------------------------------------------------

Prim.cons.                       TAAAAAAATCTGATTCCCGCTTCTCCAAGTANGTACTTCTTGAAATCTCTCTTTTTCTTA

                                        970       980       990      1000      1010      1020
                                          |         |         |         |         |         |
PrapaeCcBV1jonction2squenceNCB   AATAATACGACAATTTTAAGATGTTTTATTAAAATTGACGTATTATTTCTTACACCCTTA
PrapaeCcBV1jonction2sample15     ------------------------------------------------------------
PrapaeCcBV1jonction2sample30     ------------------------------------------------------------

Prim.cons.                       AATAATACGACAATTTTAAGATGTTTTATTAAAATTGACGTATTATTTCTTACACCCTTA

                                       1030      1040      1050      1060      1070      1080
                                          |         |         |         |         |         |
PrapaeCcBV1jonction2squenceNCB   TTTCTTATACTTATATCTTTCACAATAAATATTATCGCTACTATATAATGCACTTAGTAC
PrapaeCcBV1jonction2sample15     ------------------------------------------------------------
PrapaeCcBV1jonction2sample30     ------------------------------------------------------------

Prim.cons.                       TTTCTTATACTTATATCTTTCACAATAAATATTATCGCTACTATATAATGCACTTAGTAC


                                       1090      1100      1110      1120      1130      1140
                                          |         |         |         |         |         |
PrapaeCcBV1jonction2squenceNCB   TTTTAAACGCGGCCCGCGGACGTATTTNCACGCTTAAGCTTGGCCCTCAACATGTNAAAG
PrapaeCcBV1jonction2sample15     ------------------------------------------------------------
PrapaeCcBV1jonction2sample30     ------------------------------------------------------------

Prim.cons.                       TTTTAAACGCGGCCCGCGGACGTATTTNCACGCTTAAGCTTGGCCCTCAACATGTNAAAG

                                       1150      1160      1170      1180      1190      1200
                                          |         |         |         |         |         |
PrapaeCcBV1jonction2squenceNCB   GTTGCCCATACCTGATCTAGAAGGTATGACTTAACATCAGGCGAACCACCTGACTCTATG
PrapaeCcBV1jonction2sample15     ------------------------------------------------------------
PrapaeCcBV1jonction2sample30     ------------------------------------------------------------

Prim.cons.                       GTTGCCCATACCTGATCTAGAAGGTATGACTTAACATCAGGCGAACCACCTGACTCTATG

                                       1210      1220      1230      1240      1250      1260
                                          |         |         |         |         |         |
PrapaeCcBV1jonction2squenceNCB   CCCTCAGTTATATATTTGAATGTCGTTGATCTTTTGCCATAATGTTTTCCTCTCGATGTA
PrapaeCcBV1jonction2sample15     ------------------------------------------------------------
PrapaeCcBV1jonction2sample30     ------------------------------------------------------------

Prim.cons.                       CCCTCAGTTATATATTTGAATGTCGTTGATCTTTTGCCATAATGTTTTCCTCTCGATGTA

                                       1270      1280
                                          |         |
PrapaeCcBV1jonction2squenceNCB   TTTCTTCACCGTAAAACCCTC
PrapaeCcBV1jonction2sample15     ---------------------
PrapaeCcBV1jonction2sample30     ---------------------

Prim.cons.                       TTTCTTCACCGTAAAACCCTC
```
