## Supplementary figures and images for "Somatic chromosomal integration of polydnavirus during parasitism triggered their germline infiltration in multiple lepidopteran families"

### Supplementary Figure 1

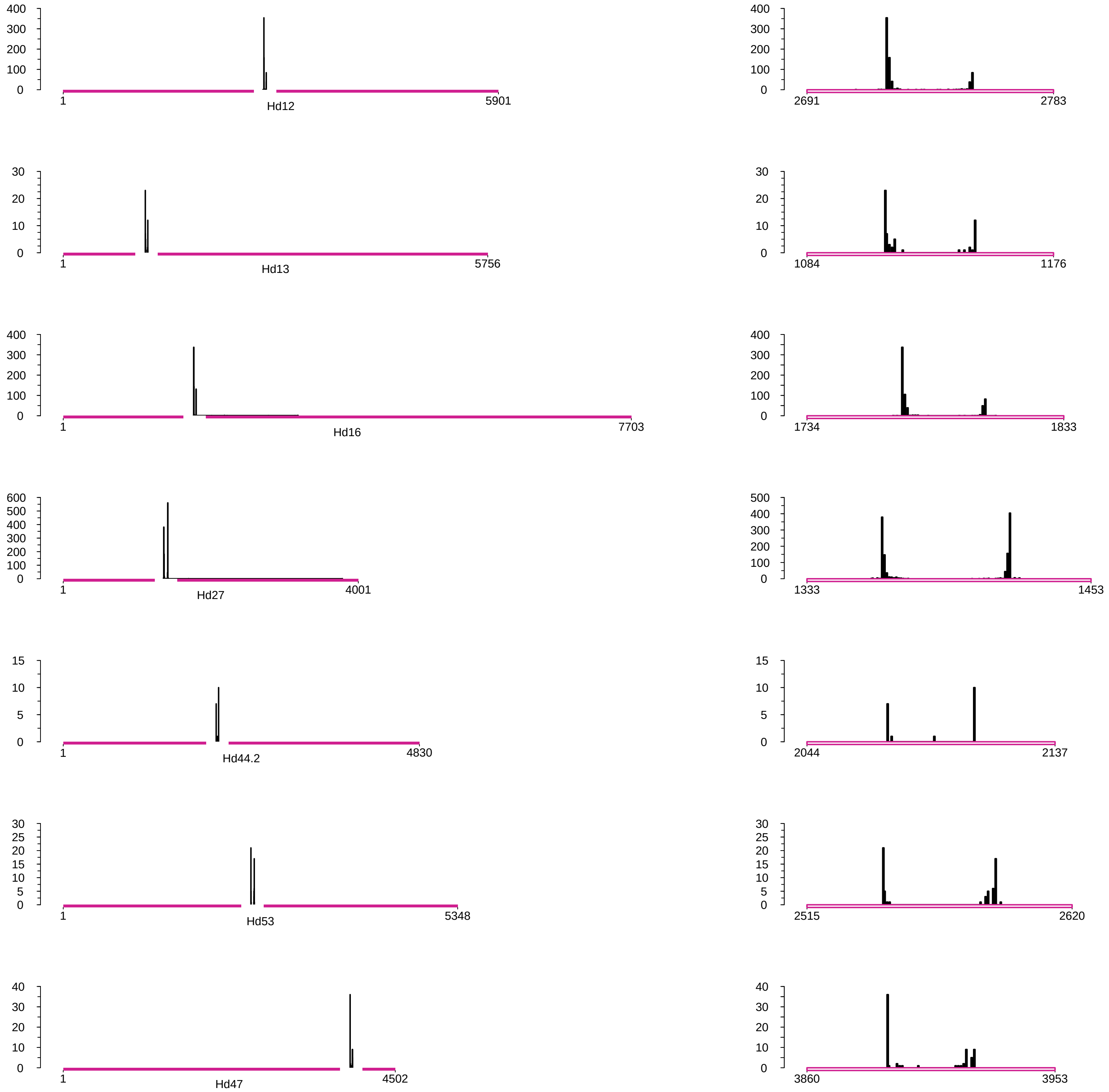

### Supplementary Figure 2

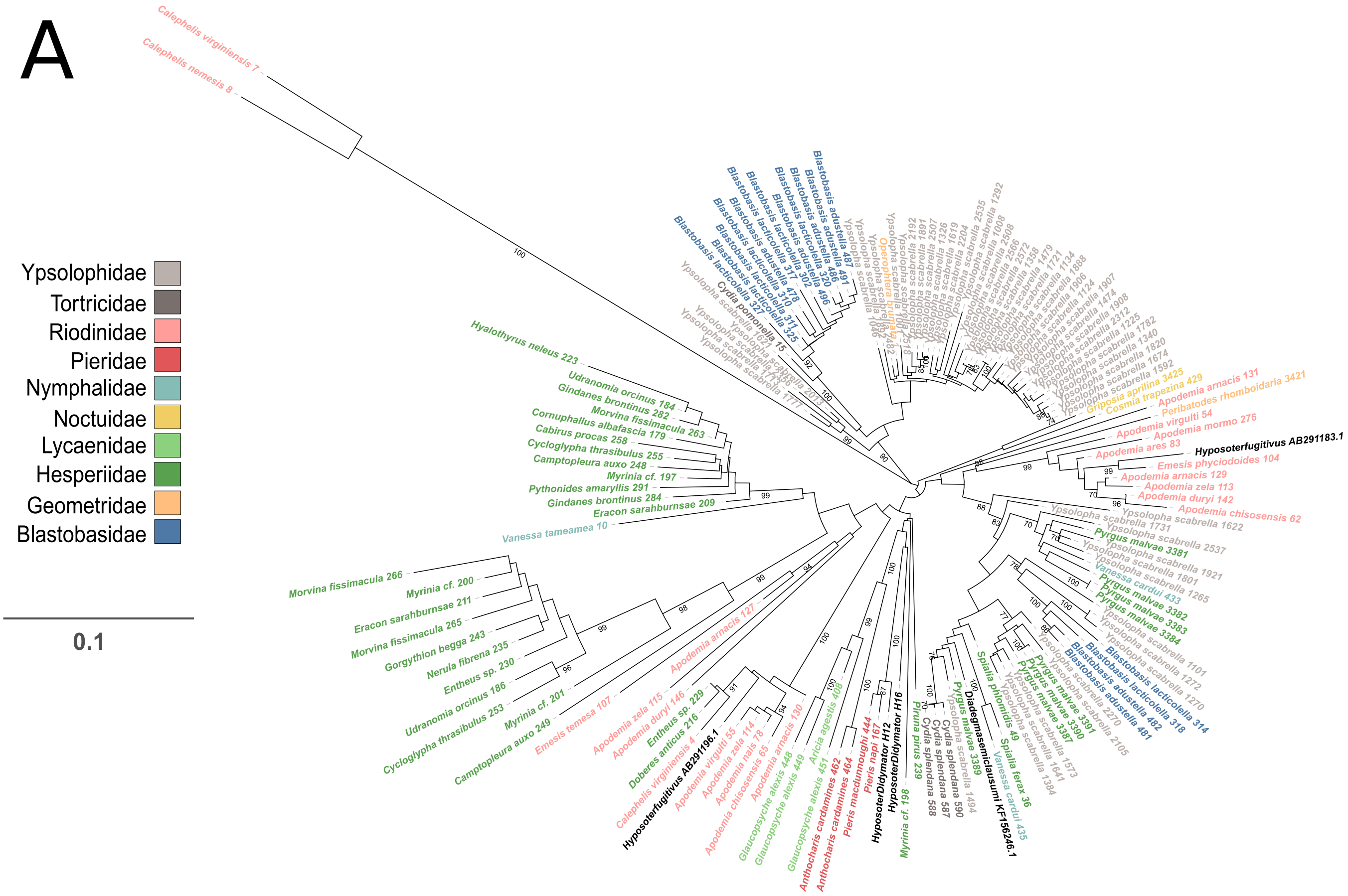

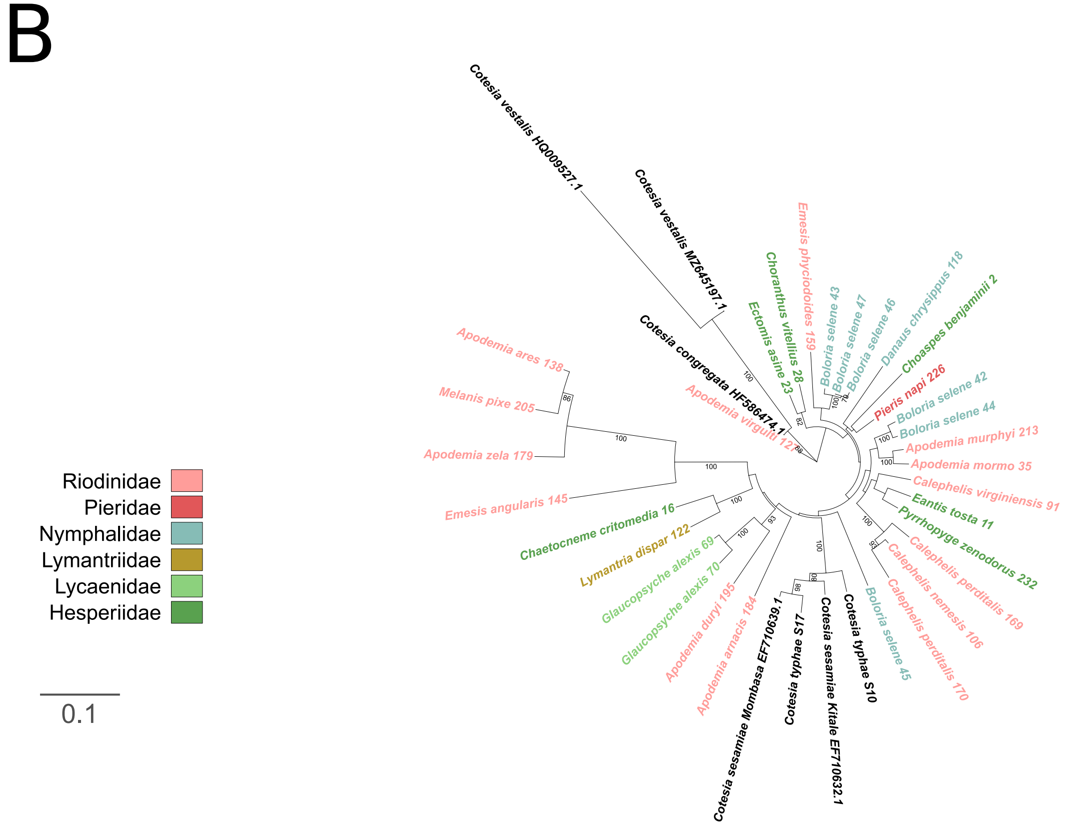

### Supplementary Figure 3

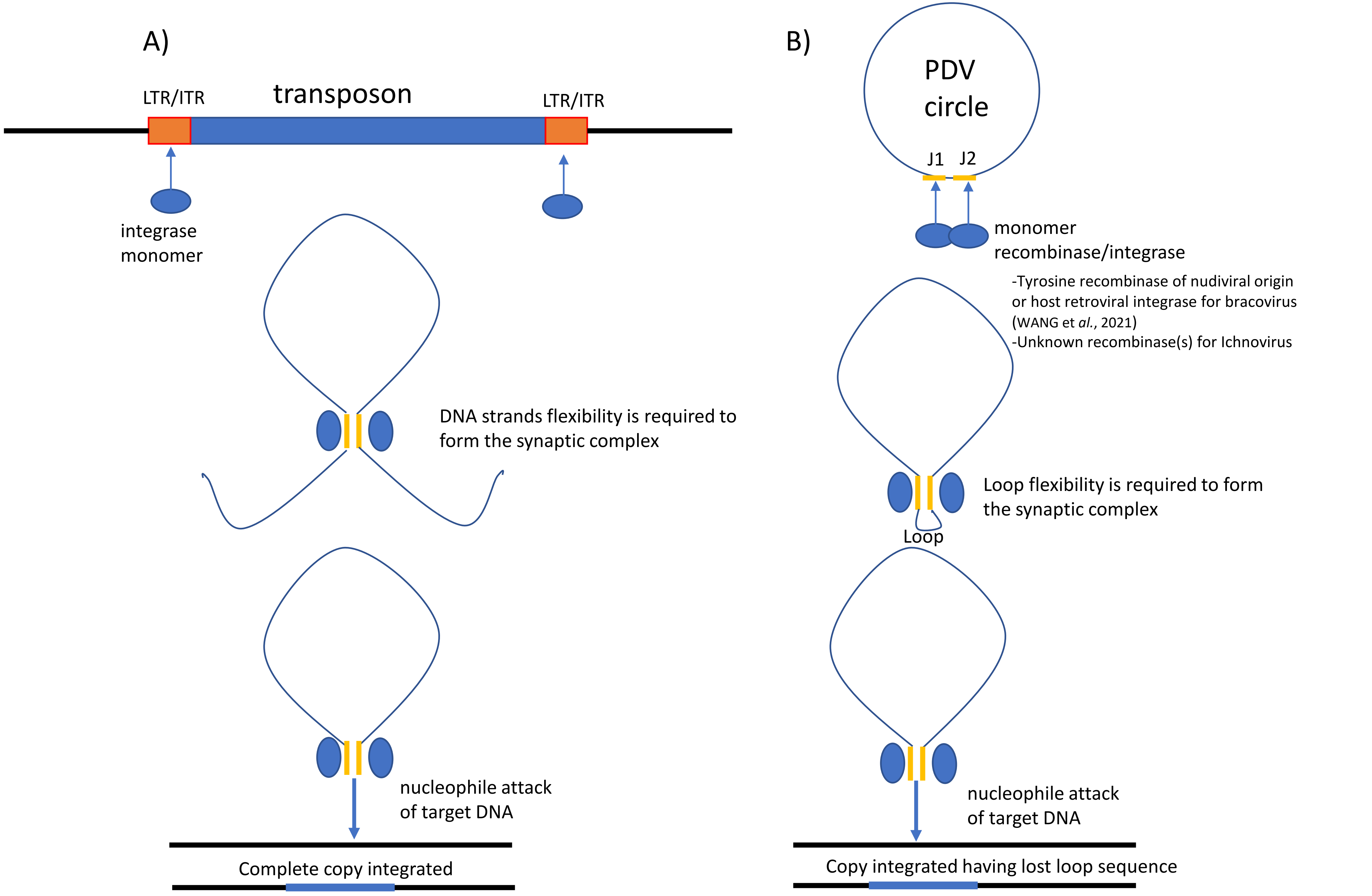
